# Early Developmental Origins of Cortical Disorders Modeled in Human Neural Stem Cells

**DOI:** 10.1101/2024.06.14.598925

**Authors:** Xoel Mato-Blanco, Suel-Kee Kim, Alexandre Jourdon, Shaojie Ma, Andrew T.N. Tebbenkamp, Fuchen Liu, Alvaro Duque, Flora M. Vaccarino, Nenad Sestan, Carlo Colantuoni, Pasko Rakic, Gabriel Santpere, Nicola Micali

## Abstract

The implications of the early phases of human telencephalic development, involving neural stem cells (NSCs), in the etiology of cortical disorders remain elusive. Here, we explored the expression dynamics of cortical and neuropsychiatric disorder-associated genes in datasets generated from human NSCs across telencephalic fate transitions in vitro and in vivo. We identified risk genes expressed in brain organizers and sequential gene regulatory networks across corticogenesis revealing disease-specific critical phases, when NSCs are more vulnerable to gene dysfunctions, and converging signaling across multiple diseases. Moreover, we simulated the impact of risk transcription factor (TF) depletions on different neural cell types spanning the developing human neocortex and observed a spatiotemporal-dependent effect for each perturbation. Finally, single-cell transcriptomics of newly generated autism-affected patient-derived NSCs in vitro revealed recurrent alterations of TFs orchestrating brain patterning and NSC lineage commitment. This work opens new perspectives to explore human brain dysfunctions at the early phases of development.

**One-sentence summary:** The temporal analysis of gene regulatory networks in human neural stem cells reveals multiple early critical phases associated with cortical disorders and neuropsychiatric traits.

## INTRODUCTION

The mammalian cerebral cortex emerges from the dorsal telencephalon, where organizing centers secrete morphogenic signals that instruct the spatiotemporal identity of neural stem cells (NSCs) ^1–5^. Radial glial (RG) cells serve as NSCs, initially generating excitatory glutamatergic neurons that migrate to the overlaying cortex and eventually becoming gliogenic ^6–10^. The cerebral cortex is also populated by inhibitory γ-aminobutyric acid (GABAergic) interneurons, mostly originating from ventral telencephalic NSCs, that migrate dorsally and integrate with the excitatory neurons ^11,12^.

Disruption of these events in humans by germline and/or somatic genetic mutations or environmental insults may cause malformations of cortical development (MCDs) and neurodevelopmental disorders (NDDs) ^13–19^, characterized by aberrant morphology of the cortex and neuropsychiatric manifestations ^20–23^. However, the intricate molecular architecture of these conditions, which can manifest overlapping symptoms across distinct disorders or divergent features within the same disease, remains unclear.

Dysfunctional neuronal mechanisms during late gestation or postnatal stages have been associated with the origin of cortical and neuropsychiatric disorders ^24–31^. However, several works suggest disruption of the spatiotemporal identity of the NSCs as earlier fetal risk events ^32–38^. Despite their importance, the implications of these early events involving human NSCs in the etiology of brain diseases and the impact of their dysfunctions on the brain in postnatal life is still poorly understood. As it is challenging to explore these phases of human brain development, it is necessary to employ pluripotent stem cells (PSCs), which faithfully model many aspects of species-specific corticogenesis in vivo including sequential generation of neuronal subtypes and glia ^39–43^.

Here, we curated gene lists associated with MCDs and NDDs and explored their expression across the in vitro and in vivo progression of human NSCs (hNSCs), exploiting our previously reported transcriptomic data and other brain datasets. We identified “critical phases” for each disorder, defined as distinct NSC states during human telencephalic development that are more vulnerable to gene disruptions. Moreover, we identified putative transcription factor (TF) networks along hNSC progression, revealing convergent genes across different diseases and unique interactions associated with a specific disorder, and further in silico simulated the impact of their depletions on each neural cell type across human corticogenesis. Using multiple ASD patient-derived NSC lines, we modeled the influence of intrinsic genetic background on individual early brain development, and unveiled frequent alteration of brain patterning and NSC fate regulators across donors. We created a resource to explore the expression dynamics of disease genes and gene sets in the neural cells of the datasets used in this study which is accessible at <u>NeMO/genes</u>.

## RESULTS

### Enrichment of cortical disorder risk genes across human neuronal differentiation in vitro

We compiled lists of genes associated with MCDs, including microcephaly (MIC), lissencephaly (LIS), cobblestone (COB), heterotopia (HET), polymicrogyria (POLYM), congenital hydrocephalus (HC), focal cortical dysplasia (FCD), mTORopathies (mTOR), NDDs including schizophrenia (SCZ), attention deficit hyperactivity disorder (ADHD), major depressive disorder (MDD), bipolar disorder (BD), autism spectrum disorder (ASD), obsessive-compulsive disorder (OCD), Tourette syndrome (TS), developmental delay (DD), and neurodegenerative disorders such as Alzheimer’s (AD) and Parkinson’s diseases (PD) (see methods; Supplementary Table 1).

Cortical disorders are commonly associated with neuronal dysfunctions ^24,26,27,31^. Using a bulk RNA-sequencing (RNA-seq) dataset generated from human induced PSC (hiPSC) lines traversing neurogenesis ^44^, we explored gene expression dynamics utilizing GWCoGAPS (genome-wide Coordinated Gene Activity in Pattern Sets) analysis, which defined expression patterns (p1-12) depicting the transcriptomic progression of cells across differentiation (Supplementary Fig. 1a and b) ^45,46^. We checked enrichment of disorder-associated risk genes in the GWCoGAPS patterns and found that most of the risk gene lists were enriched in neurons, consistent with prior works ^24,26,27^, except for MIC-associated genes which were enriched in progenitors (Supplementary Fig. 1c). However, we have recently shown that many neuropsychiatric disorder-associated genes, while globally enriched in excitatory and inhibitory neurons, are also expressed in early telencephalic organizers (also called patterning centers, PCs) and/or dorsal and ventral NSCs during primate corticogenesis ^37^. Thus, we investigated the dynamics of these risk genes in earlier cell states in vitro.

We have previously established an in vitro system where hPSC-derived forebrain NSCs were serially passaged, and exposed to varying doses of FGF2 to create a gradient of differentiation potential (Supplementary Fig. 2a) ^40^. This assay is hereafter referred to as the “NSC progression protocol”. We demonstrated that hNSCs, transitioning from pluripotency, spontaneously recapitulate a state resembling the cortical hem, the dorsomedial-posterior organizer of the telencephalon, before generating neurons. RNA-seq analysis along this time-course identified 24 GWCoGAPS patterns (p1– p24), describing transcriptomic dynamics across the hNSC progression (Supplementary Fig. 2b; <u>NeMO/CoGAPS</u>). Here, we focused on those GWCoGAPS patterns most specific to discrete NSC states triggered by FGF2 doses and passage (PS, Fig. 1a). By performing projection analysis ^47^ of single-cell and microdissection expression data of primate developing neocortex ^37,40,48–50^, we identified the in vivo equivalent cells for the in vitro hNSCs of each passage, and further annotated hNSCs as neuroepithelial/organizer progenitors at early passages (PS2-3), excitatory neurogenic RG at mid passage (PS4), and late neuro-/glio-genic RG at late (PS6-8) passages (Fig. 1a, Supplementary Fig. 2b-d; methods), complementing our previous work ^40^. Hence, we interrogated the expression of the disease gene sets throughout the progression of hNSCs and found differential enrichment across different phases. For instance, MIC and HC associated genes were enriched at early neuroepithelial states, while LIS, mTOR, and SCZ risk genes were enriched in mid- and late-RG cells (Fig. 1b). Analysis of a similar dataset ^51^ yielded congruent findings, highlighting risk genes enriched in hNSCs and in the derived neurons and glia (Supplementary Fig. 1d-f). Thus, NSCs likely play critical roles in early cortical disorder events that can be dissected using this in vitro system.

**Fig. 1.**
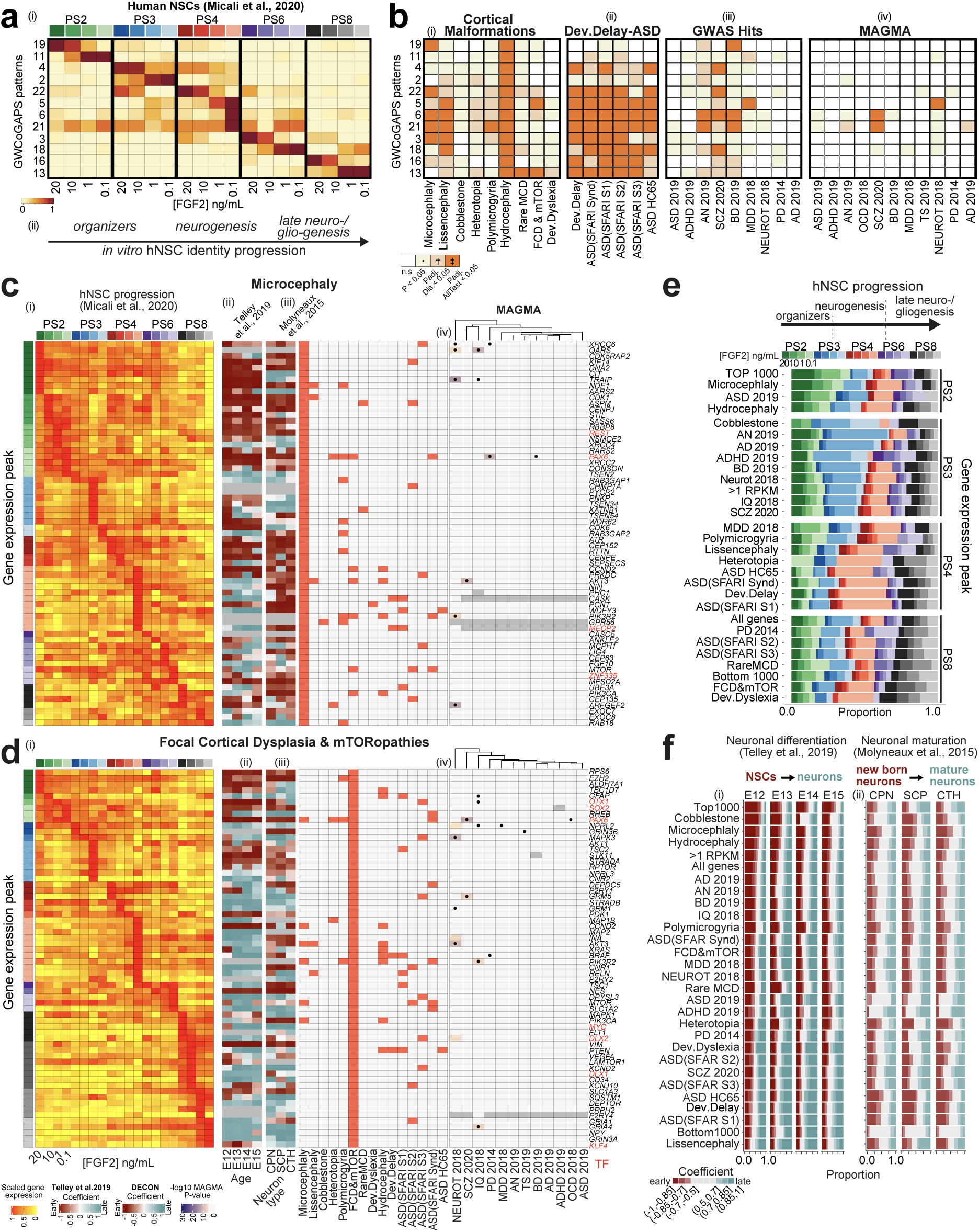
Expression dynamics of risk genes across cortical neurogenesis. **a)** Selected GWCoGAPS patterns dissecting hNSC progression across passages and FGF2 doses ^40^. (ii) Schema of hNSC progression. **b**) Enrichment of disease gene sets in the GWCoGAPS patterns. n.s.: not significant; P: uncorrected P-values at p<0.05; Padj.Dis: significance correcting by each disease independently; Padj. AllTest: significance after multiple-testing correction using the whole dataset. **c, d**) (i) Expression levels of risk genes in FGF2-regulated hNSC progression, ordered by the temporal peak of expression (left column colored by passage and FGF2 dose). (ii-iii) Slope of gene expression change across (ii) neuronal differentiation of age-specific RG cells and (iii) maturation of different neuronal classes from developing mouse cortex. (iv) Disease associations of each gene (left panel), and log10 p-value of the MAGMA gene-level test of association with each GWAS dataset (right panel). Black dots indicate a top-hit gene in the corresponding GWAS publication, based on genome-wide significant loci. **e**) Proportion of genes for each disease showing expression peaks at each passage and FGF2 condition. Additional categories: all genes in the dataset; genes with average expression of >1 log2 RPKM (RPKM>1); 1000 genes at the top and bottom ranks of expression, respectively (top and bottom 1000). Categories are ordered as: PS2 and PS3 high to low, PS4 and PS8 low to high. No diseases with most genes in PS6 were found. **f)** Proportion of genes classified in different bins of expression fold change in (i) differentiating NSCs and (ii) maturing neurons. Coefficient=slope of expression change as described in panels c and d.

### Expression dynamics of risk genes across human cortical NSCs progressing in vitro

We ordered the disease gene sets based on the average expression peak across the temporal progression of the in vitro hNSCs ^40^, revealing a specific distribution of the risk genes for each disorder (Fig. 1ci-di and Supplementary Fig. 3ai-vi; <u>NeMO/genes</u> and <u>NeMO/diseasegenesets</u>). For instance, MIC-specific risk genes involved in cell replication, such as *ASPM* and the centrosome component *CENPJ*, were highly expressed in the first passages (PS2-3), suggesting that the earliest founder cells of the telencephalon might be more vulnerable to perturbations of the cell cycle machinery than later RG cells (Fig. 1ci). HC-associated genes, also highly expressed in PS2-3, included well-known regional patterning regulators, such as *ARX, FGFR3*, *WNT3*, and *GLI3* ^37,52,53^, suggesting putative dysfunctions of patterning and/or fate commitment of the early NSCs at the origins of this disorder (Supplementary Fig. 3bi). These results denote a precise phase during early NSC expansion critical for MIC and HC. LIS-associated genes, including *DCX*, ranked higher in the differentiating neurogenic RG cells of PS4 with low FGF2 (Supplementary Fig. 3li), while FCD- and mTOR-associated risk genes, such as the mammalian target of rapamycin (*mTOR*) and its targets *DEPTOR* and *KLF4*, were highly expressed in late NSCs (Fig. 1di). Thus, mid-fetal neurogenesis might be critical for LIS, while FCD and mTOR might involve late progenitors, as previously reported ^54^. Thus, by systematically comparing the temporal maximum expression of risk genes across all the diseases, we distinguished “early-organizer-”, “mid-neurogenic-” and “late-neuro/gliogenic”-related disorders (Fig. 1e).

We further compared the distribution of genes associated with 269 diseases, including cancers, neuropsychiatric, and non-brain diseases (e.g. cardiovascular), from DisGeNET ^55^, across the progression of the hNSCs (Supplementary Fig. 4a). Most cortical disorder-related genes were highly enriched at PS4 low FGF2, indicating that this state reflects a neuronal-specific signature which is less represented in non-brain diseases. Altogether, these results suggest a precise expression dynamic of the genes associated with each brain disease across the progression of cortical hNSCs, revealing putative disease-specific “critical phases”, that is, temporal windows when NSCs are more vulnerable to dysfunctions that might lead to the onset of a disorder.

### Expression dynamics of risk genes across mammalian neurogenesis in vivo

To assess the implication of risk genes in cortical NSCs in vivo, we leveraged a mouse single-cell (sc)RNA-seq dataset, where cortical RG cells were profiled across sequential differentiation towards glutamatergic neurons from embryonic day (E) 12 to E15 ^56^. We determined a slope of expression change for each gene across differentiation and summarized the analysis showing expression fold change bins for each gene set (Fig. 1cii and dii, fi, Supplementary Fig. 3aii-vii). The result highlights expression of risk genes in the NSCs and during their differentiation for all the disorders. “Early” diseases, such as MIC and HC, exhibited a higher proportion of genes in RG cells from E12 to E15, unlike “later” disorders, e.g., ASD and LIS, which had a higher proportion of genes in the neuronal differentiation phase. These data confirm the involvement of risk genes in NSCs and their transition into neurons during corticogenesis.

We next exploited the DeCoN resource ^57^, where mouse cortico-thalamic (CThPN), sub-cortical and cortico-spinal motor (ScPN), and interhemispheric callosal (CPN) projection neurons were profiled by RNA-seq from E15.5 to postnatal day (P)1. We determined a slope of expression change for risk genes from immature to mature state for each neuron subtype, and summarized their distribution in disease-specific bins (Fig. 1ciii and diii, 1fii, Supplementary Fig. 3aiii-viii). Gene sets exhibited greater expression differences in RG-to-neuron comparisons than in neuronal maturation stage comparisons (Fig. 1f). Risk genes associated with disorders such as MIC, COB, HC, and ASD (ASD HC65 and SFARI S1) showed high enrichment in immature neurons, suggesting a transient implication in the terminal differentiation process. In contrast, diseases such as ADHD and SCZ exhibited more genes with higher expression in mature neurons. Most diseases showed consistent patterns among all neuronal subtypes; however, some disease exhibited subtype-specific biases. For instance, a larger proportion of COB-associated genes exhibited decreasing expression specifically across SCP neuronal maturation. Together, these data suggest a prominent function of cortical disorder-associated risk genes during the progression of RG cells and neuronal transition. While each diseases encompassed genes with preferentially increased or decreased expression trends in RG-to-neuron and/or immature-to-mature neuron transitions, all diseases included genes in both trajectories (Fig. 1f), suggesting multiple critical phases in the pathogeny of cortical disorders.

Finally, we assessed the enrichment of two well-known ASD-associated risk genes, the RNA-binding protein FMRP and the chromatin remodeling factor CHD8, and their targets identified from embryonic and adult mouse brain datasets, in our disease gene sets and DisGeNET (Supplementary Fig. 4a and b; Supplementary Table 2). Consistent with previous findings, we observed enrichment of FMRP and CHD8 targets among ASD- and Developmental Delay (DD)-associated genes. However, this enrichment extended to other disorders, including SCZ, depending on the cell types where the targets have been identified. For instance, embryonic FMRP targets were enriched in HET and HC genes, while targets determined in the adult brain were enriched in LIS. Similarly, CHD8 targets were associated which an extended range of diseases including non-GWAS ASD gene sets, MIC, and cancers. This analysis denotes a transcriptional intersection of FMRP and CHD8 with their targets aross hNSC progression, suggesting that they may play roles in both disease-specific and shared developmental pathways, including, but not restricted to, ASD.

### Disease risk genes expressed in brain organizers

We have previously described six hiPSC lines differentiated into forebrain NSCs showing variable propensity to form dorsoposterior (DP) or anteroventral (AV) telencephalic organizer states in their transition from pluripotency, and consequentially exhibiting divergent bias towards excitatory glutamatergic or inhibitory GABAergic neuronal lineages, respectively ^40^ (Supplementary Fig. 20ci). This was further supported in this current study by projecting the GWCoGAPS (p1-30) defined in the RNA-seq analysis of those lines onto the PCs from our recent macaque developing brain dataset ^37^ (Supplementary Fig. 5a; <u>NeMO/CoGAPSII</u>). The analysis distinguished dorsal-versus ventral-biased lines expressing genes of the cortical hem or rostral patterning center (RPC), respectively, at day in vitro (DIV) 8. Leveraging this in vitro dataset, we assessed the enrichment of each disease risk gene list throughout the differentiation trajectory of the hNSC lines, at DIV 8, when the organizer states emerge, and later at DIV17 and 30 in neurons ^40^, an assessment hereafter referred to as the “neuronal differentiation protocol” (Fig. 2a; Supplementary Table 3). Diseases were sorted by the proportion of dorsal-biased genes at DIV8. For instance, HET risk genes showed a dorsal bias in each phase, in contrast to COB, which was enriched in genes expressed early in the AV organizer and later in excitatory neurons, suggesting multiple spatiotemporal risk phases. Genes associated with ASD, ADHD, FCD, or SCZ were present early in both ventral- and dorsal-biased NSCs, and later in the derived neurons, consistent with our previous report ^37^.

**Fig. 2.**
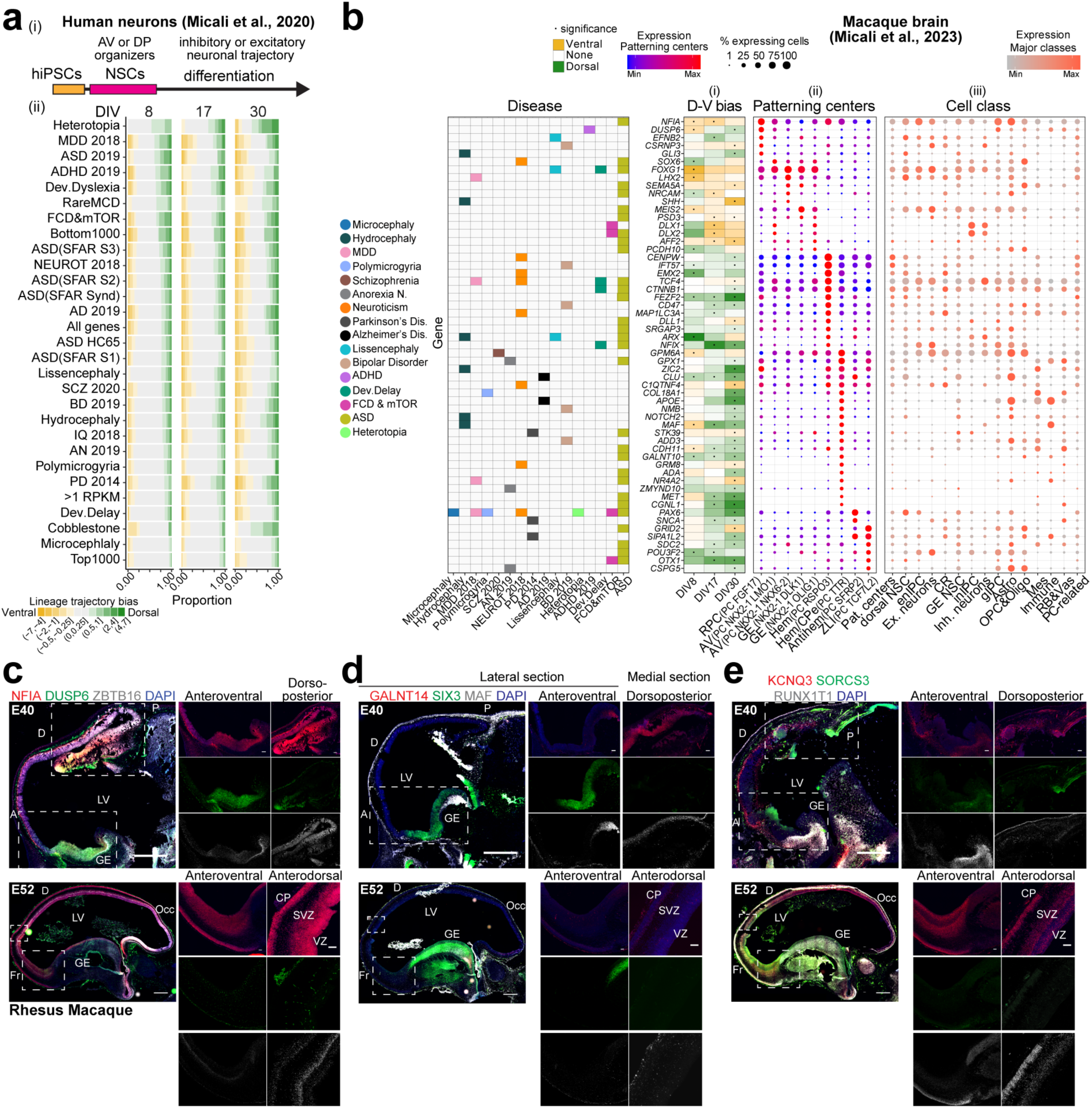
Expression of risk genes in brain organizers. **a)** (i) Neuronal differentiation protocol. (ii) Proportion of disease-associated genes with dorsal or ventral bias across differentiation of the 6 hNSC lines ^40^. Fold change between dorsal and ventral expression binned in categories. All genes, RPKM>1, top and bottom 1000 categories as in Fig. 1. **b)** (i) PC markers with dorsal or ventral expression bias in the 6 hNSC lines at DIV 8-30 and (ii-iii) gene expression in (ii) PC clusters and (iii) other cell subtypes from the macaque dataset ^37^. Filtered PC markers with significant dorsoventral bias and disease association (left panel) are displayed. **c-e)** RNAscope of macaque sagittal fetal brains. Scale bar: 500 µm (panoramic), 100 µm (zoom-in).

To determine whether the risk genes expressed in hNSCs in vitro at the organizer state (DIV8) were present in the telencephalic organizers in vivo, we assessed their expression in macaque ^37^ and mouse ^58^ developing brain scRNA-seq datasets, where the PCs have been profiled (Fig. 2b, Supplementary Fig. 5b and c). For example, neural fate regulators such as *FOXG1* and *MEIS2*, associated with ASD, showed ventral bias in vitro and an equivalent expression pattern in the AV organizers [i.e., RPC (PC FGF17), AV (PC NKX2-1) or ganglionic eminence (GE, RG NKX2-1)] in vivo. In contrast, *EMX2*, associated with Neuroticism, showed a dorsal bias in vitro and expression in the cortical hem clusters [(PC RSPO3) and hem/choroid plexus epithelium (CPe, PC TTR)] in vivo. Other genes, e.g. *PAX6* and *OTX1*, were dorsal in vitro and expressed in the dorsal antihem (PC SFRP2) and zona limitans intrathalamica (ZLI, PC TCF7L2), respectively, in vivo. Notably, *MAF* and *ARX*, associated with HC, exhibited a shift of the DV bias over time in vitro. Moreover, many genes (e.g., *MEIS2*, *ARX*, *FOXG1*) became also expressed later in neurons, in vitro and in vivo, consistent with our previous report^37^. RNAscope analysis of developing macaque telencephalons validated the expression pattern of several genes identified from the macaque and mouse dataset analysis (Fig. 2b, Supplementary Fig. 5b and c), including *DUSP6* and *SIX3* in the AV organizer, *GALNT14* in the cortical hem, and *MAF* in both AV and DP organizers at E40, and in the cortex at E52, paralleling the trend observed in vitro (Fig. 2c-e). These data indicate that the telencephalic spatiotemporal expression pattern of cortical disease-associated genes is faithfully reproduced in this in vitro system. These risk genes can function in organizers and early NSCs, further supporting our hypothesis that altered patterning events and/or NSC progression are potential early causes of these cortical diseases.

### Disease networks across human NSC progression in vitro

We identified putative TF regulatory networks across the progression of the in vitro cortical hNSCs for each disease gene set, hereafter referred to as “disease regulons”. Using RcisTarget, which searches for overrepresented TF binding motifs in gene sets, we identified 31 disease regulons involving 29 core TFs, targeting from 5 to 485 genes in 9 diseases (Fig. 3a; Supplementary Table 4). Core TFs and targets were required to co-occur in a disease gene list; hence, only gene sets containing TFs could yield regulons. Moreover, gene sets of different sizes exhibited variable power to yield regulons. Permutation analysis revealed 22 highly disease-specific regulons (P<0.05) and 7 core TFs, including FOXG1 and DLX2, found more frequently by chance (P>0.1), commensurate to their small number of targets. Many disease regulons were enriched in targets significantly correlated (p-value < 0.05) with the expression level of the core TF along the progression of the hNSCs: 12 regulons were enriched in targets positively correlated, 6 in targets negatively correlated, and 4 in both types of targets (Fig. 3b and Supplementary Fig. 6; Supplementary Table 4). In several regulons enriched in positively correlated targets, some targets exhibited expression peak at different passages than their core TF, reflecting possible TF-target regulatory mechanisms, negative regulatory effects, or unspecific associations between TFs and their motifs. The characterization of these disease regulons was further extended to the mouse in vivo ^56,57^ and human in vitro ^40^ datasets, suggesting potential functions across RG cell progression and neuronal maturation, as well as during both dorsal and ventral neuronal specification, respectively (Supplementary Fig. 7a-d).

**Fig. 3.**
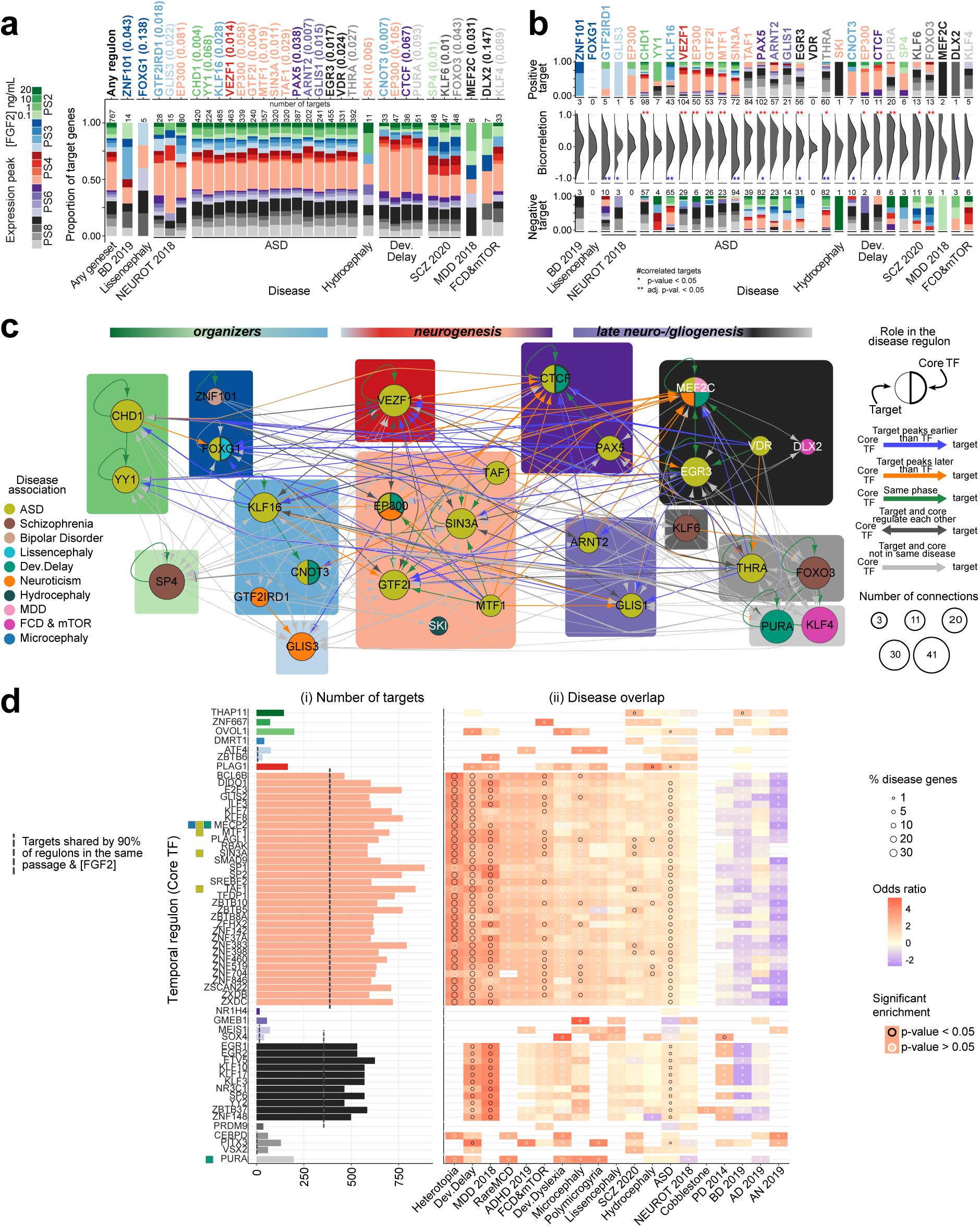
Sequential gene regulatory networks across hNSC progression. **a)** Proportion of target genes for each disease regulon with expression peak across passages and FGF2 conditions. Each core TF (top axis) is colored by its expression peak with p-value associated. The category “Any geneset/Any disease” includes all the genes of the disease regulons. Number of targets per regulon indicated on the top bar. **b)** Distribution of expression correlations between core TFs and their targets for each disease with positively and negatively correlated targets, and their number shown in the top and bottom summary bar plots, respectively. The color of the bars and TF labelling represent the expression peaks. Significantly high number of positively or negatively correlated targets are marked as ‘*’: p-value < 0.05, or ‘**’: corrected p-value < 0.05. **c)** Predicted gene regulatory network of the core TFs in the disease regulons with nodes representing genes colored by disease. For a TF found in multiple disease regulons, the thicker stroke indicates the disease in which it is a core TF; the node size indicates the connections to other core TFs. Background colors indicate gene expression peak in the in vitro hNSCs, same colors as in (a). Edge colors represent the expression peak relation between core TF and target, if they regulate each other, if associated with same or different disease. **d)** Temporal regulons ordered by the core TF peak expression across hNSC progression, same colors as in (a). Core TFs of disease regulons are indicated and colored by disease as in panel c. MECP2, although not a core TF in the disease regulons, is disease-associated. i) Number of targets for each regulon. ii) Overlap of temporal regulons with disease shown as odds ratio of the regulon gene enrichment in each disease (color of the grid), fraction of disease genes present in a regulon (dot size), and the significance of the enrichment (dot color).

We next investigated putative regulatory relationships among the 29 unique core TF networks that showed temporal progression along hNSC development (Fig. 3c). RcisTarget revealed core TFs connecting multiple regulons within the same NSC phase, spanning different NSC phases, or having reciprocal interaction. For instance, the multiple disease-associated TF MEF2C, peaking in PS8 RG cells, ranked among the most connected core TFs. In addition of being a core TF associated with Neuroticism, MEF2C was also predicted as target of other TFs associated with ASD, DD, and MDD, including CHD1 and TAF1, peaking at earlier cell states (Fig. 3c; Supplementary Table 4). Furthermore, allowing each core TF from any disease regulon to be a potential target in other regulons, we uncovered additional regulatory connections among diseases, and identified KLF4 as the core TF most interconnected with other core TFs. These data suggest intricate crosstalk between networks and convergence of risk genes within key hub TFs, supporting the hypothesis of multiple vulnerability phases throughout hNSC progression for a disease.

### Temporal networks across human NSC progression in vitro

Other genes, while currently not classified as risk factors, may still be involved in disease-regulons, as TFs or targets, and the pathobiology of a disease. Thus, we determined “temporal-specific” regulons for all the genes peaking at each hNSC phase employing RcisTarget, without requiring a core TF to be disease-associated or its targets to co-occur in a given disease gene list. We identified 60 temporal regulons and assessed the enrichment of disease genes in these networks (Fig. 3d and Supplementary Fig. 7e; Supplementary Table 5). Since the targets in the temporal regulons differ from those in the disease set, the analysis identified a distinct set of core TFs with little overlap with the disease core TFs, which included MTF1, SIN3A, TAF1, and PURA. Regulons detected in PS4 low FGF2 and PS8 high FGF2 showed a high number of targets and were shared among diseases. For example, the core TF MTF1 associated with ASD, putatively targeted genes at PS4 enriched in other disorders, including Fragile X Messenger Ribonucleoprotein 1 (*FMR1*) associated with HET, *AUTS2* and *MECP2* associated with DD, cannabinoid receptor 1 (*CNR1*) associated with FCD (Supplementary Table 5). However, the heightened connectivity observed at PS4 and PS8 could result from shared binding motifs between promiscuous core TFs. Thus, we clustered the binding motifs based on similarity and identified a prominent cluster of Krüppel-like family TF (KLF/SP) binding sites for the TFs of PS4 and PS8 (Supplementary Fig. 8 a-c). Although the potential targets of the TFs within these two NSC phases were mutually exclusive, diverse sets of disease-associated risk genes were enriched in KLF/Sp targets. By contrasting disease and temporal regulons, we identified multiple members of the KLF/SP family in both (Fig. 3a and d). Notably, neurogenic PS4 NSCs correspond to RG transitioning to neuronal intermediate precursor cells (nIPCs) and new-born neurons during macaque corticogenesis (Supplementary Fig. 2d), suggesting that this high connectivity and shared binding motifs across genes might be an intrinsic feature of this cell state. Taken together, our regulon analyses reveal molecular mechanisms shared among different diseases, highlighting target genes where multiple lesions might converge during NSC progression, as well as unique pathogenic signaling within the same disease ^59,60^. However, the convergent or divergent expression dynamics of TFs and targets also suggest varying severity for a perturbation during neurogenesis, depending on the TF-target’s role in a network and the timing they are functioning. This raised the need to investigate the effect of a gene network perturbation across different phases of neurogenesis.

### *In silico* assessment of the gene regulatory network disruption across human corticogenesis

We leveraged single-cell multiomic data of the developing human cortex of four donors, spanning post-conceptional week (PCW)16 to 24 ^61^. Using CellOracle, we reconstructed cell type-specific gene regulatory networks (GRN) ^62^, and simulated the effect of perturbations of disease-related TFs and chromatin modifiers, which in vivo could be caused by genetic mutations and/or environmental factors, across distinct phases of corticogenesis. To gain higher resolution in NSCs subpopulations, we reanalyzed the Trevino et al., 2021 dataset including markers from monkey ^37^ and mouse ^56^ studies (see methods). This led to distinguish multiple NSC states, including ventricular early (vRG E) and late (vRG L), outer early (oRG E) and late (oRG L), and truncated (tRG) radial glial subtypes, and their glutamatergic neuronal and glial cell progeny (Fig. 4a and Supplementary Fig. 9; Supplementary Table 6). Unlike RcisTarget, CellOracle constructs networks based on TF-target co-expression and chromatin co-accessibility. Furthermore, for a TF to be susceptible to perturbation, it must pass specific thresholds of expression levels and variance across cells (Supplementary Fig. 10). We applied CellOracle to each cell subtypes across RG progression and gliogenesis from all donors (PCW16-24) to derive transcriptomic trajectories spanning the entire spectrum of NSCs and glial cells. Additionally, we constructed GRNs across neurogenesis from individual donors at PCW20, 21 and 24, each containing complete RG-to-neuron trajectories, which served as biological replicates. We perturbed 141 TFs, of which 61 were disease-related and/or were core TFs in temporal regulons (Supplementary Fig. 10). Network centrality, reflecting the role and relevance of each TF across cell subtypes, varied among the GRN (Fig. 4b and Supplementary Fig. 11). For example, the eigenvector centrality of TCF4, implicated in multiple disorders (e.g. ASD and MDD), was prominent in vRG and oRG cells and glutamatergic neurons. However, the highest centrality was found for the SCZ-related core TF KLF6, in RG cells, neurons, and glial cells. Knock-out (KO) simulation of KLF6, which peaks at PS8 in vitro, predicted an impact on both neurogenic and gliogenic trajectories, prompting the transition of RG cells, multipotent glial precursors (mGPCs), and nIPCs to later states (Fig. 4c, Supplementary Fig. 12a and c). Similarly, perturbation of the core TF MEF2C, with high centrality in neurons (Fig. 4b), reduced vRG proportion, promoting the transition to more mature states and glutamatergic neurons (Fig. 4d, Supplementary Fig. 12a and b).

**Fig. 4.**
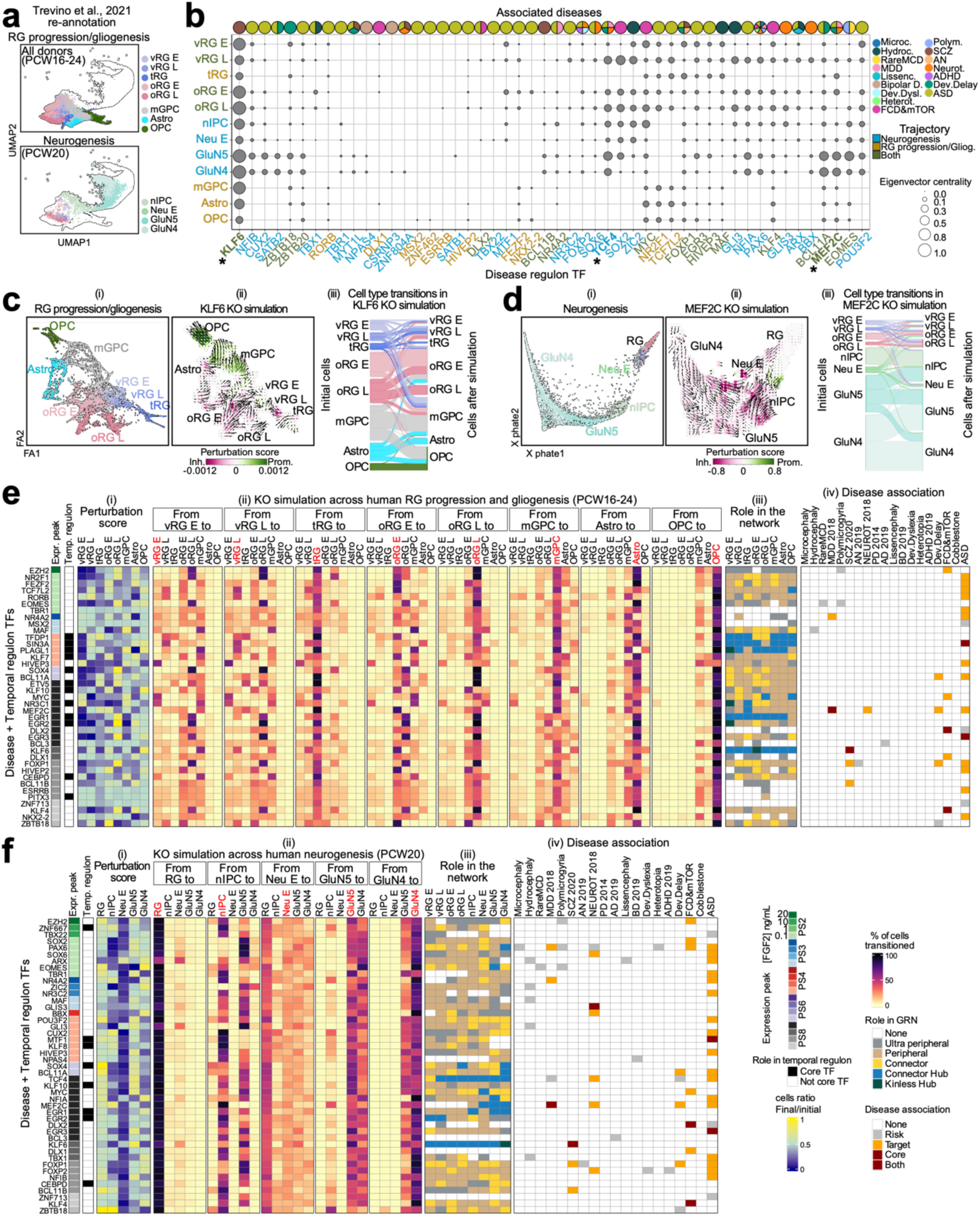
In silico perturbation of regulon genes. **a)** Trajectories from Trevino et al., 2021 analyzed in CellOracle: RG progression and gliogenesis for all donors (top); neurogenesis for each donor (bottom, only PCW20 is shown). **b)** Network centrality of disease-associated core TFs across RG progression, neurogenesis, and gliogenesis. The eigenvector centrality of a TF in the GRN of a cell type is shown by dots representing the influence of a gene in the network. Cell types (y axes) and TFs (x axes) are colored by the trajectory tested. The disease association of each TF is on the top bar. “*”: genes mentioned in the text. **c, d)** Partition-based graph abstraction (PAGA) map of RG maturation and gliogenesis (ci), and potential of heat diffusion for affinity-based transition embedding (PHATE) map of neurogenesis (di). KO simulation of (c) *KLF6* and (d) *MEF2C*. ii) Trajectory perturbation: arrows simulate cell flow after KO perturbation with color representing trajectory change, promoted (green) or depleted (red). iii) Cell transitions from original cell identities (left) and after KO simulation (right). **e, f)** KO simulation of TFs across (e) RG maturation and gliogenesis and (f) neurogenesis. TFs associated with disease and core TF of temporal regulons, selected from the test in S10 are shown. Expression peak across the in vitro hNSC progression is next to each TF. Temporal regulon column shows core TFs in the temporal regulons. i) Perturbation score indicating gain or depletion of a given cell type. ii) Cell type transitions after KO simulations. Grids represent the fraction of the original cell type (labeled red on the top) and their final identity. iii) Regulatory role of every TF in each cell type. iv) Gene-Disease association, specifying core TFs and target genes in the disease regulons.

Extending the KO simulation to all the TFs, we predicted the impact on individual cell-type fate determination across human RG progression/gliogenesis and/or neurogenesis (Fig. 4e and f, Supplementary Fig. 13 and 14). For instance, KO of KLF4, present at PS8 in both trajectories, depleted vRG cells favoring transitions to SVZ states (oRGs), reduced mGPC proportion promoting astrogenesis, and reduced nIPCs enhancing neurons, with less effects on differentiated cells. KO of NFIA and NFIB, both present at PS8 only in the neurogenic trajectory, increased RG cell proportion and depleted nIPCs and newborn neurons, respectively, unlike KO of ARX and HES1, both at PS2, which depleted RG cells incresing neurons, all cases suggesting an impact on NSC-to-neuron transition (Fig. 4f and Supplementary Fig. 14). We further applied CellOracle to a mouse developing cortex dataset ^63^ and each human donor, demonstrating similar effects of TF perturbation-induced cell transitions in human and mouse cells and across donors, strengthening our predictions (Supplementary Fig. 15). Some exceptions were observed, such as for FOXP1, possibly due to the methodology used for dataset generation, species differences, or disparities in developmental time points. These results suggest that TF lesions have cell state-restricted effects.

We further categorized the TFs by their importance within the network in each cell type, ranging from ultra-peripheral, i.e., isolated, to kinless hub, i.e., highly involved in the network (Fig. 4e and f, Supplementary Fig. 13 and 14). For instance, while KLF6 emerged as a hub in all the cell types implicated in RG progression/gliogenesis or neurogenesis, KLF4 played peripheral roles in the majority of the cells in both trajectories. EOMES (PS2) was peripheral across the glial lineage but became more connected, up to be a hub, in emerging neurons. These analyses provide a functional map of the TFs delineating their roles in each cell across NSC progression, neurogenesis, and gliogenesis. Altogether, these data emphasize the spatiotemporal context-specific effects of a lesion ^13^ and predict “risk temporal-domains” for each TF network perturbation across cortical development.

### Individual-specific critical phases in ASD patient-derived NSCs

To investigate the influence of intrinsic genetic factors on an individual’s early brain development, we newly generated NSCs from iPSCs derived from three idiopathic ASD patients (#375, #384, #434) and three controls (#290, #311, #317). Then, we differentiated NSCs into neurons applying the “neuronal differentiation protocol” and performed scRNA-seq at DIV 8 (Supplementary Fig. 20b; Supplementary Table 7). Unsupervised clustering, aided by curated cell-type markers derived from our monkey brain dataset ^37^, identified a predominant population of early neuroepithelial stem cells (NESC)/RG cells (RGEarly) along with other smaller clusters of late RG cells (RGLate), mesenchymal cells (Mes), neurons, and brain organizer progenitors with predominant anterior neural ridge/RPC-resembling cells (PC FGF17-like). We excluded cells with a high fraction of mitochondrial genes, or those coupled with marker genes that we could not confidently ascertain, resulting in a dataset of 44,311 high-quality cells for downstream analysis (Fig. 5a, Supplementary Fig. 16).

**Fig. 5.**
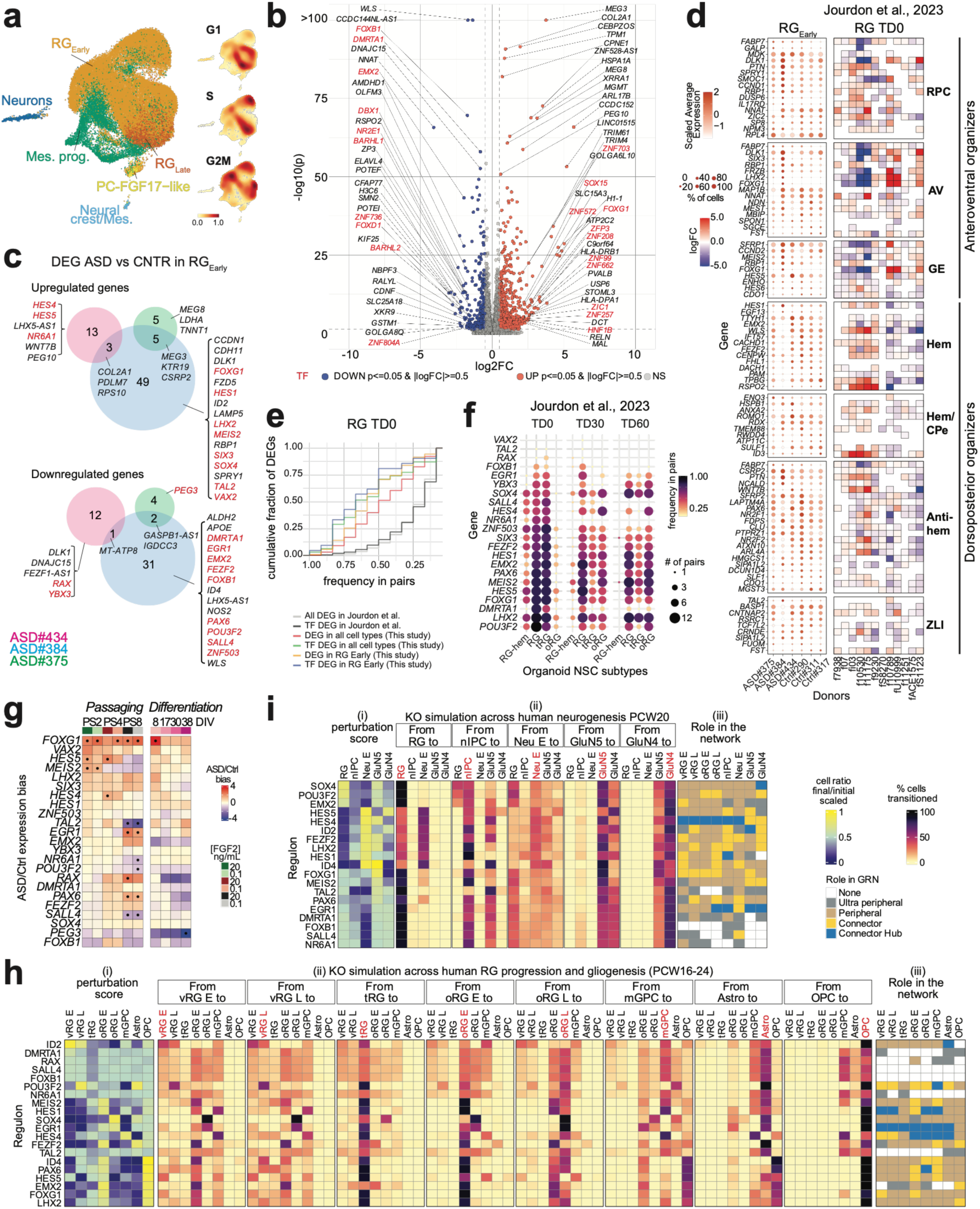
Analysis of GRNs in ASD patient-derived NSC lines. (**a**) Cell subtypes and density of the cell cycle phases, in control- and ASD-patient-derived NSCs. (**b**) DEGs between grouped ASD versus grouped control RG_Early_ cell pseudo-bulk. Fold change (FC) expression ratio between ASD and control cells (x axis) versus the significance of the differential expression (y axis). Top DEGs are labeled. (**c**) DEGs in RG_Early_ in individual ASD samples versus grouped controls. (**d**) Expression level (color gradient) and percentage of cells (dot size) expressing patterning center genes from Micali et al., 2023 in RG_Early_ of each line (left), and differential expression of the same genes across ASD-control organoid pairs in the RG cluster at TD0, from Jourdon et al., 2023. (**e**) Cumulative fraction of DEGs identified in our study (y-axis, grouped into different DEG subsets by color) found to be differentially expressed in varying frequencies among the ASD-control pairs in RG cluster at TD0 from Jourdon et al., 2023 ^34^ (x-axis). Distribution for all genes/TFs differentially expressed in Jourdon et al. is given as reference (black/grey lines). (**f**) TFs differentially expressed in RG_Early_ in individual ASD samples (from **c**) also found significantly perturbed (upregulated or downregulated) in at least 1 ASD-control pair in the DEG data from Jourdon et al., in different NSC subtypes, at 3 organoid stages (dot size shows number of ASD pairs significantly perturbed, dot color shows frequency among total number of pairs tested), sorted by recurrence in RG cluster at TD0. (**g**) Expression ratio of TFs found in **c** across sequential passaging and differentiation of ASD versus Control NSCs. (**h, i**) CellOracle KO perturbation of differentially expressed TFs from panel c identified across (h) RG progression/gliogenesis (20 TFs tested) and (i) neurogenesis (19 TFs tested in PCW20). Perturbation score (i), cell type transitions (ii), role of every TF in each cell type (iii) as in Fig. 4.

NSCs derived from ASD patients displayed higher proportions of cells in G1 phase, suggesting an enhanced commitment to neurons ^64^ (Supplementary Fig. 17). Next, we grouped RGEarly cells by cell cycle phase and donor, creating donor-balanced pseudobulk samples. Principal component analysis (PCA) of gene expression revealed high PC6 in ASD versus control samples. Then, projecting the passaged hNSC dataset ^40^ indicated the highest level in the neurogenic NSCs of PS4, aligning with the more distinguished ASD critical phase (Fig. 1e, Supplementary Fig. 18a and b). Next, we performed differential gene expression (DEG) analysis and found 1,259 DEGs between ASD and controls, including 131 TFs (absolute logFC >0.5 and adjusted p-value <0.05) (Fig. 5b; Supplementary Table 8). Notably, many brain patterning genes were deregulated (upregulated or downregulated) in ASD-derived NSCs compared to controls. Telencephalic DP genes, such as *WLS*, *DMARTA1*, *EMX2*, and *FEZF2*, were decreased, while the AV regulator *FOXG1* was upregulated, as previously reported ^33^, along with *ZIC1* and *RELN*.

To explore individual contributions to these gene expression differences, we performed DEG analysis on RGEarly cells from individual ASD lines versus grouped control lines. We identified 75 up-regulated (11 TFs) and 50 down-regulated (12 TFs) genes in ASD NSCs, exhibiting donor-specific expression variation (Fig. 5c; Supplementary Table 9). Notably, ASD#384 showed upregulation of AV patterning regulators (*FOXG1*, *MEIS2*, and *SIX3)*, and downregulation of DP regulators (*DMRTA1*, *EMX2*, and *WLS)*, while ASD#434 showed enhanced expression of *HES4/5* and *WNT7B* and decreased *DLK1*, all involved in patterning centers’ signaling network ^37^, suggesting an unbalanced telencephalic regionalization in these donors. Other cell subtypes in the culture, including Mes progenitors and RGLate cells also showed individual-specific gene expression variation (Supplementary Fig. 18d; Supplementary Table 9). Analysis of primate telencephalic organizer genes ^37^ across the ASD and control NSCs, further confirmed the expression patterns of these regulators across individual lines. Furthermore, transcriptomic comparison with a dataset from idiopathic ASD- and paired control-derived brain organoids from 13 families ^34^ validated the frequent deregulation of these organizer genes in the RG cluster at terminal differentiation day 0 (TD0) (Fig. 5d and Supplementary Fig. 19a). However, there were less pronounced differences in the expression of cortical regional markers (frontal, motor-somatosensory, occipital, and temporal) (Supplementary Fig. 18e). These findings suggest more compromised dorsoventral than frontotemporal axis patterning in ASD-affected telencephalons.

We observed a high recurrence of the DEGs identified in the RGEarly of our ASD lines within the DEGs detected across all the organoid NSC subtypes (RG-hem, RG, tRG, oRG) of the ASD-control pairs from Jourdon et al., 2023 (Fig. 5e; Supplementary Fig. 19a-c). This recurrence was more pronounced for the differentially expressed TFs. Notably, most of the 23 TFs differentially expressed in the RGEarly cells in our lines (Fig. 5c), including the NSC fate regulators *POU3F2*, *LHX2*, and *FEZF2*, were frequently deregulated also across the multiple donors, especially at TD0 (Fig. 5f), exhibiting variation in the direction of change across pairs (Supplementary Fig. 19d). Imprinted genes, including *DLK1*, *IGF2*, and the long non-coding RNA *MEG3/8*, were overrepresented among the DEGs in our ASD lines (p-value = 1.909e-07, odds ratio = 4.80; Fig. 5b; Supplementary Table 8), and were also frequently deregulated among these organoid pairs (Kstest P-value in RG at TD0= 4.950281e-05), consistent with findings in post mortem and ex vivo ASD patient samples ^65–68^, and mouse brain ^69,70^, or potentially due to iPSC reprogramming ^71^. This intersection strengthens our analysis, which was performed with a limited number of ASD probands, and suggests a frequent deregulation of major TFs involved in brain patterning and lineage commitment in the early NSCs of ASD-affected telencephalons.

Next, control and ASD NSCs were serially passaged with varying FGF2 doses or differentiated, and 60 samples were subjected to bulk RNA-seq (Supplementary Fig. 20a and b). Projection into in vitro ^40^ and in vivo ^37^ transcriptomic dimensions defining telencephalic regionalization confirmed the AV identity of ASD#384, and suggested a dorsal identity for Cntr#290, with no evident trajectory bias for the other lines (Supplementary Fig. 20c and d). Most of the 23 TFs deregulated in individual ASD lines showed consistent deregulation among the same donors and across passages and/or differentiation (Fig. 5g and Supplementary Fig. 21; Supplementary Table 10). For example, *FOXG1*, *MEIS2*, and *HES4/5* were upregulated, while *RAX* and *FOXB1* were downregulated at early passages and/or during differentiation in ASD NSCs. Other TFs, e.g., *MEIS2*, *TAL2*, and *PAX6*, shifted their expression from early to late passages or differentiation stages. Additionally, projection of PC6 from scRNA-seq in this bulk data showed high levels in ASD NSCs again in the critical PS4 (Supplementary Fig. 18c; <u>NeMO/PCA</u>). These findings indicate persistent alterations of the regulatory mechanisms acting in ASD NSCs as the cells traverse cortical maturation.

We recreated CellOracle networks in human fetal brain cells, to include some of the TFs filtered out in the previous analysis (Fig. 4e and f) and simulated the loss-of-function effect of each of these 23 TFs on cell fate determination during corticogenesis (Fig. 5h and i). Many TFs showed pleiotropic effects. For instance, *KO* of POU3F2 promoted the maturation of early RG cells and the differentiation of mGPC while hindering neuronal differentiation (Fig. 5h and i). Most perturbations putatively affected different phases of RG progression/gliogenesis, depleting RG cells at different states (e.g., MEIS2, SOX4, FEZF2, and EMX2), and altering the proportion of glial cell and oligodendrocyte precursors (OPC) (Fig. 5h). TF perturbations resulted in more cell-restricted effects across neurogenesis, affecting RG cells (HES5/4, FEZF2, and LHX2), and promoting neuron transition (ID4) (Fig. 5i). We further assessed the role of each TF in the networks. For example, HES4 emerged as a hub specifically in late RG cells during gliogenesis, but in all cell types during neurogenesis (Fig. 5h and i). These findings further underscore the significance of the context-specific effect of a TF depletion across NSC trajectories in complex cortical disorders such as ASD. In conclusion, these data highlight the potential of this in vitro modeling, and its integration with in vivo datasets, to gain insights into the vulnerable phases unique to each individual’s neurodevelopment and the phenotypic diversity within brain disorders.

## DISCUSSION

We have explored the pathobiology of cortical disorders in NSCs traversing the early phases of human brain development. Our recent findings indicate that risk genes associated with mental diseases are expressed during early patterning events and may play a role in RG cell identity specification, suggesting an even earlier fetal origin of these disorders than commonly observed ^37^. Additionally, we have previously established an in vitro system where hNSCs spontaneously recapitulate cortical fate transitions, including organizer states, neurogenesis and gliogenesis, offering a faithfully experimental model of the early phases of human brain development ^40^. Leveraging this in vitro system with in vivo data from mouse, macaque and human developing brain, we identified many risk genes involved in MCDs and NDDs which are expressed in organizing centers and in hNSCs as they progress through dorsal excitatory or ventral inhibitory neuronal lineages. Therefore, we defined their dynamic roles in the GRNs across the sequential state transitions of NSCs. Each disease gene set exhibits a distinct dynamic pattern during NSC progression and differentiation, characterized by specific “critical phases” that we defined as vulnerability periods, when most risk genes are highly active and disruptions impacting their activity may yield more pronounced effects. For example, microcephaly tends to have an earlier onset than focal cortical dysplasia, a finding consistent with other works ^23^.

The regulon analysis reveals TF cross-interactions that orchestrate multiple sets of risk genes, elucidating distinct pathogenic networks within a same disease, as well as how a same gene may converge on a broader network implicated in multiple disorders ^59^. Interestingly, our analysis “size” the role of a TF in a network, demonstrating that it might function as a hub in one regulon or as a peripheral target in another, reflecting its relevance in the critical phase and the restricted cellular etiology of a disease. Indeed, the impact of a mutation might depend on when and in which cell type it affects a gene’s function during brain development ^13,21,27,31^ as well as on the genomic context ^72^. Thus, our in silico gene perturbation analysis provides a valuable resource for interrogating and predicting the potential effects of TF loss of functions on discrete cell transitions, in sequential spatiotemporal contexts during human corticogenesis. The analysis reveals that different gene alterations can affect common lineage trajectories, potentially causing similar dysfunctions in distinct disorders. Conversely, the same perturbation at different developmental stages can yield distinct outcomes, leading to divergent symptoms in individuals carrying the same genetic variant. Therefore, our findings help in modeling and comprehending phenotypic heterogeneity within a disorder, pleiotropic effects of risk genes, and overlapping features and comorbidities of diverse diseases. Notably, we identified KLF TFs, known to be implicated in brain development and disorders ^73^, linking multiple regulons particularly in neurogenic and late NSCs in vitro. Consistently, these TFs exhibited the highest connectivity across neural cells in vivo, in line with a previous report ^74^, suggesting a role as master regulators in corticogenesis.

Using our in vitro model, we retrospectively explored the influence of the intrinsic genetic background on neurodevelopment in individuals with idiopathic ASD. To complement the limited number of donor-derived NSC lines, we integrated a larger dataset of ASD-control pair-generated organoids ^34^. We identified frequent deregulation of telencephalic organizer genes and neural fate regulators in early ASD NSCs, including *FOXG1*, *MEIS2*, *POU3F2*, and *FEZF2*, whose deleterious variants have been already identified in patients ^75–77^. Complementing the genomic studies, we now provide a time frame during brain development when perturbations of these risk genes can potentially lead to the disease. Building on previous works linking ASD pathology to NSC dysfunctions ^32–34^, we further suggest that altered patterning and spatiotemporal instruction of telencephalic NSCs by dysfunctional organizing centers may be among the earliest causes of altered neurodevelopmental trajectories in idiopathic ASD individuals. Further work is required to investigate the causality of these TF alterations and their link with clinical phenotypes. Additionally, we simulated the impact of these TF perturbations during human corticogenesis, revealing the phases when their function is potentially essential and their alteration critical. It would be interesting in future works to experimentally assess the impact of these perturbations across various genomic contexts and investigate phenotypic variability in a large number of donors ^72^.

The NeMO resource that we created enables the visualization of risk gene expression dynamics across multi-species’ cortical development, helping to identify molecular features of a disease and select appropriate study models. In conclusion, this work highlights a strategy to facilitate the assessment of very early phases of human brain development in vitro, and screen for individual-specific disease phenotypes. This approach opens new avenues to conceive more precise therapeutic interventions targeting NSC dysfunctions in the pathogenesis of cortical disorders.

## ACKNOWLEDGMENTS

This study was supported in part by NIDA Merit Award DA023999 (to P.R.); NIH grant R01HG010898-01 (to G.S. and N.S.); Instituto de Salud Carlos III Spain and European Social Fund grant MS20/00064 (to G.S.); grants PID2019-104700GA-I00 and PID2022-140137NB-I00 funded by /AEI/10.13039/501100011033 (to G.S.); grant 202230-30 from Fundació La Marató de TV3 (to G.S.); NIH grants HG012108, HG010898, HG012483, MH130991, U01MH116488, U01DA053628 (to N.S); NIH grant R01 MH109648 and Simons Foundation grant # 632742 (to F.M.V.); MacBrain Resource Center NIH Grant MH113257 (to A.D.). Data sharing and visualization via NeMO Analytics was supported by grants R24MH114815 and R01DC019370.

## AUTHOR CONTRIBUTION

N.M., G.S., S-K.K., and C.C. conceived the study. X.M., G.S. and C.C. performed the majority of the bioinformatic analyses. X.M., A.J. and S.M. performed analysis of the scRNA-seq data from iPSC lines and Fig. 5 and related figures. C.C. generated NEMO online resource. G.S., N.M., C.C. and S-K.K. supervised computational analyses. P.R., N.S., F.M.V. and A.D. provided resources. A.T.N.T and F.L. generated iPSC lines. N.M. and S-K.K. performed experiments with iPSCs. N.M. prepared fetal monkey brains for RNAscope. P.R., N.S. and F.M.V., provided additional supervision and guidance. N.M., G.S. and X.M. wrote the manuscript. N.M. and G.S. directed the work. All authors discussed and edited the manuscript.

## COMPETING INTERESTS

The authors declare no conflict of interest.

## DATA AVAILABILITY

The expression information for all the studies used in this report can be accessed at the following links. NeMO Analytics link for individual genes (<u>NeMO/genes</u>): nemoanalytics.org/p?l=Blanco2024&g=FOXG1 GWCoGAPS transcriptomic patterns (p1-24) from in vitro NSC data from Micali et al., 2020 (<u>NeMO/CoGAPS</u>): nemoanalytics.org/p?p=p&l=Blanco2024&c=Micali2020_HsNSCpassageFGF2.nmfP24&algo=nmf The summed expression of disease gene lists (<u>NeMO/diseasegenesets)</u>: nemoanalytics.org/p?p=p&l=Blanco2024&c=Blanco2024_CtxDiseaseGeneLists&algo=pca Principal Component analysis of DIV 8 scRNA-seq data in ASD and control lines (<u>NeMO/PCA</u>): nemoanalytics.org/p?p=p&l=Blanco2024&c=Blanco2024_Day08scASDvCON_RGearlyPseudoBulkPC s&algo=pca GWCoGAPS transcriptomic patterns (p1-30) from in vitro neuronal differentiation from Micali et al., 2020 (<u>NeMO/CoGAPSII</u>): https://nemoanalytics.org/p?p=p&l=Blanco2024&c=Micali2020_NeuronDiff_30&algo=nmf

## SUPPLEMENTARY FIGURES

**Supplementary Fig. 1. Related to Fig. 1.**
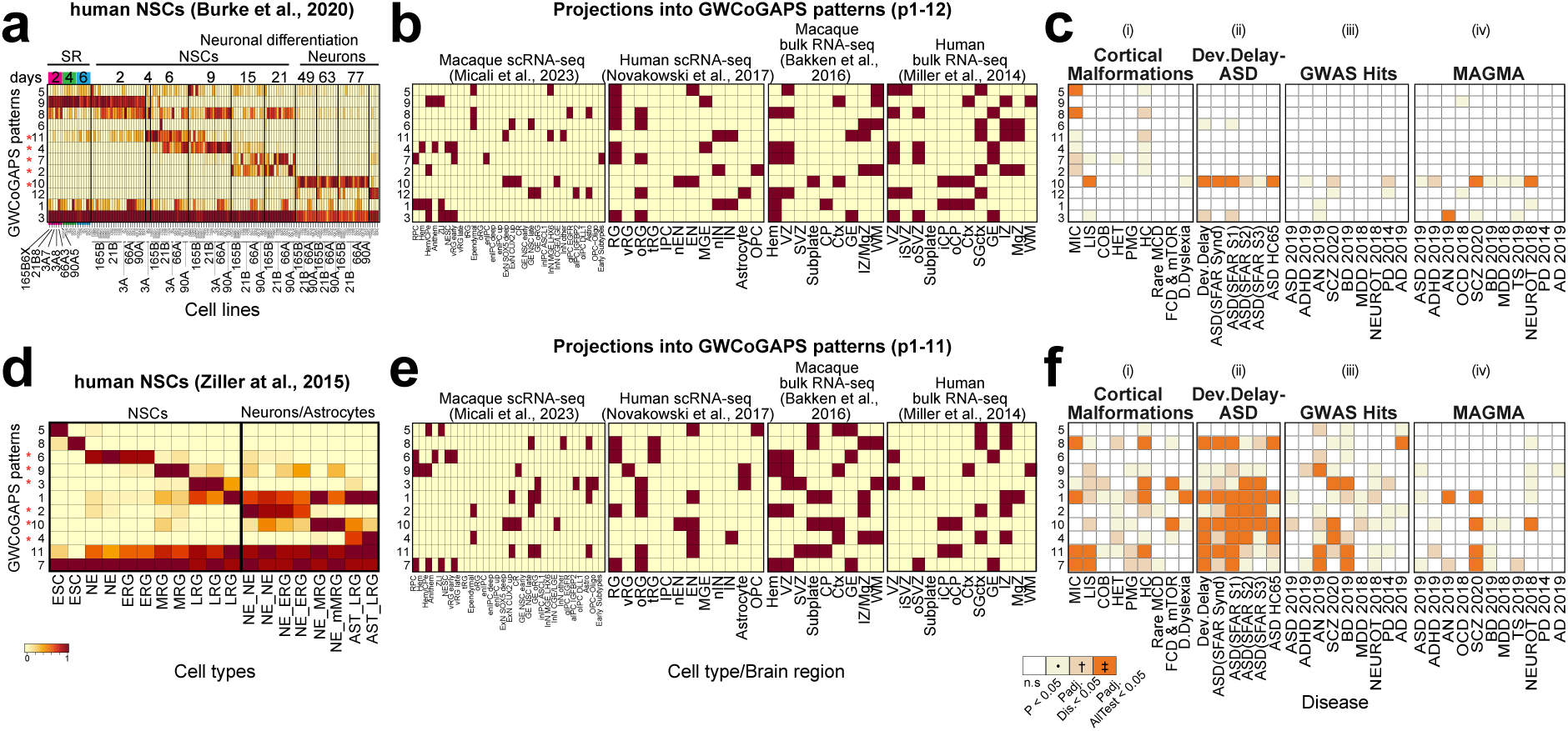
Cortical disorder-associated risk genes expressed in human neurons in vitro. **a and d**) GWCoGAPS analysis defined transcriptomic patterns across (a) neuronal differentiation of human NSC-derived from multiple iPSC lines ^44^ (p1-12), and (d) across sequentially passaged human NSCs derived from H9 ESCs ^51^ (p1-11). Column labels indicate steps of differentiation from self-renewal (SR) of the iPSCs to NSCs and terminally differentiated neurons (a), and from NSCs to terminally differentiated neurons and astrocytes (d). (*) indicates the patterns defining the strongest changes across time and distinguishing discrete cortical cells in (b and e). **b and e**) The telencephalic identity of the in vitro cells at different phases of differentiation was confirmed by projection of scRNA-seq data from the developing macaque ^37^ and human ^48^ telencephalon, and bulk RNA-seq data from microdissected developing human ^49^ and macaque ^50^ cortex into the GWCoGAPS patterns from (a) and (d). Thresholding within each in vivo dataset was used to show tissues and cells with highest expression of each GWCoGAPS pattern, i.e. dark cells indicate high levels of in vitro transcriptomic patterns in the in vivo data. D: dorsal; V: ventral; ESC: embryonic stem cells; NE: neuroepithelial stem cells; ERG: early radial glia; MRG: midradial glia; LRG: late radial glia; NE-NE: neurons derived from NE; NE-ERG: neurons derived from ERG; NE-MRG: neurons derived from MRG; NE-LRG: neurons derived from LRG; AST-LRG: astrocytes derived from LRG. RG: radial glia; oRG: outer RG; vRG: ventricular RG; tRG:truncated RG; IPC: intermediate precursor cell; nEN: new excitatory neurons; EN: excitatory neurons; MGE: medial ganglionic eminence; GE: ganglionic eminence; nIN: new inhibitory neurons; IN: inhibitory neurons; Astro: astrocytes; OPC: oligodendrocyte progenitor cells; VZ: ventricular zone; SVZ: subventricular zone, i: inner, o: outer; CP: cortical plate; WM: white matter; IZ: intermediate zone; MGZ: marginal zone; SGctx: subgranular cortex; Ctx: cortex. **c and f**) Enrichment analysis of the disease gene sets in GWCoGAPS patterns from panels a (c) and d (f). n.s.: not significant; P: uncorrected P-values at p<0.05 (yellow); Padj. Dis: significance correcting by each disease independently (light orange); Padj. AllTest: significance after multiple-testing correction using the whole dataset (orange).

**Supplementary Fig. 2. Related to Fig. 1.**
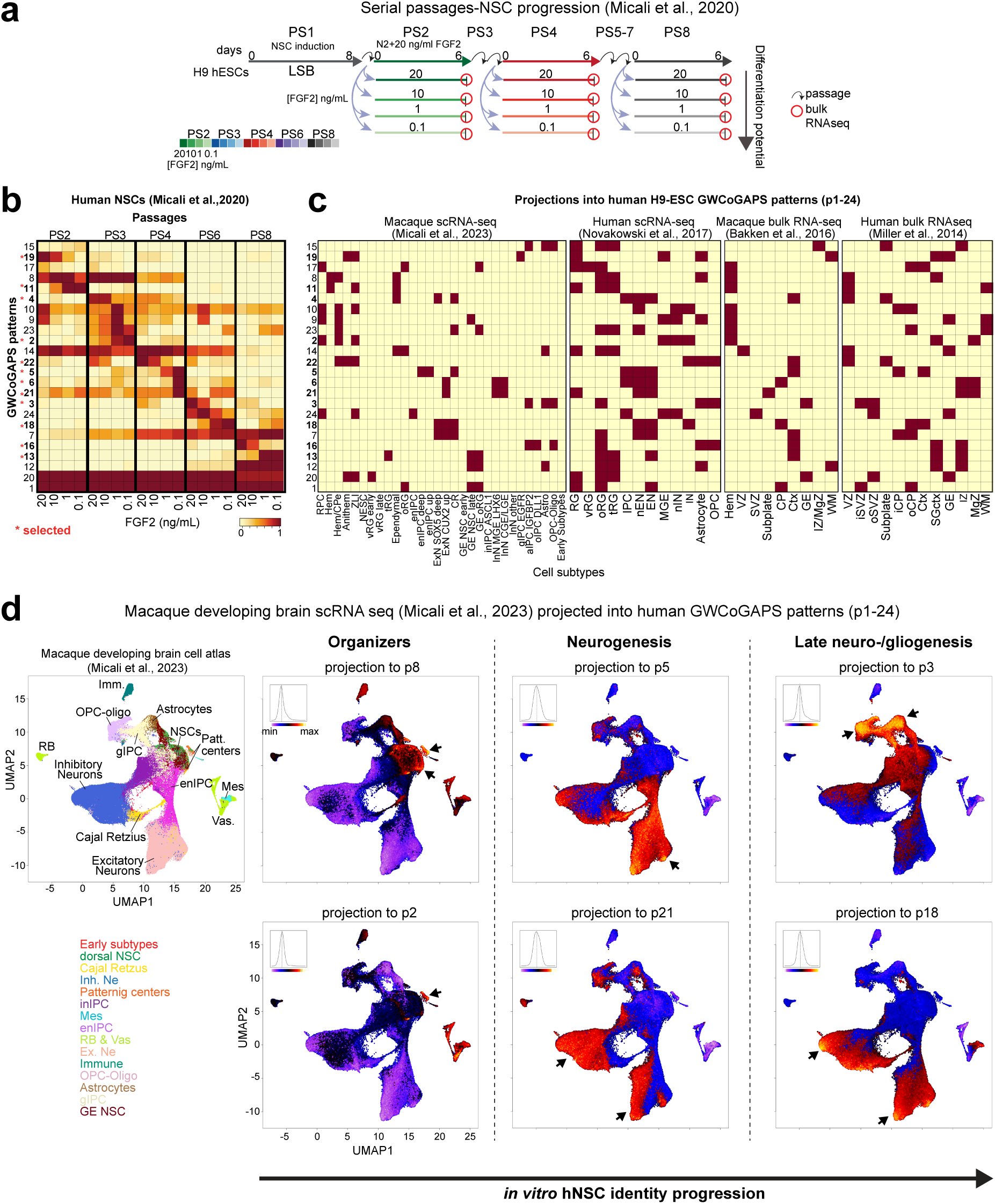
Sequentially passaged hNSCs recapitulate cell states and transcriptomic dynamics *of* in vivo corticogenesis. **a)** Scheme of H9 human embryonic stem cell (hESC) differentiation and progression of NSCs across sequential passages (PS) in presence of different FGF2 doses from Micali et al., 2020 ^40^, here defined as NSC progression protocol. N2 + LSB (LDN193189 + SB431542) medium was applied at PS1 for 8 days, then hNSCs were serially passaged in N2 + 20 ng/mL FGF2 medium every 6 days up to PS8. In parallel, cultures of NSCs at PS2, 3, 4, 6, and 8 were subjected to FGF2 modulation (20, 10, 1, 0.1 ng/mL) for 6 days during their terminal passage before RNA collection. **b**) Heatmap depicting GWCoGAPS patterns (p1-24) describing progression of hNSCs across passages (PS2-8) and FGF2 dose from Micali et al., 2020 ^40^. (*) indicates the patterns defining the strongest changes across passage and FGF2 and distinguishing discrete cortical cells in (c). These selected GWCoGAPS patterns are shown in Fig. 1. **c**) Projection of scRNA-seq data from the developing macaque ^37^ and human ^48^ telencephalon, and bulk RNA-seq data from microdissected developing human ^49^ and macaque ^50^ cortex into GWCoGAPS patterns from (b). Thresholding within each in vivo dataset was used to show tissues and cells with highest expression of each GWCoGAPS pattern, i.e. dark cells indicate high levels of in vitro transcriptomic patterns in the in vivo data. Abbreviations as in S1. **d**) Projection of scRNA-seq data from developing macaque brain ^37^ into specific GWCoGAPS patterns from ^40^. Early passage patterns (p8 and p2) show high expression in in vivo organizers; mid- and late passage patterns (p5 and p21) have high expression in in vivo neurogenesis, and late passage patterns (p3 and p18) show high expression in both late neurogenesis and gliogenesis in vivo (<u>NeMO/CoGAPS</u>).

**Supplementary Fig. 3. Related to Fig. 1.**
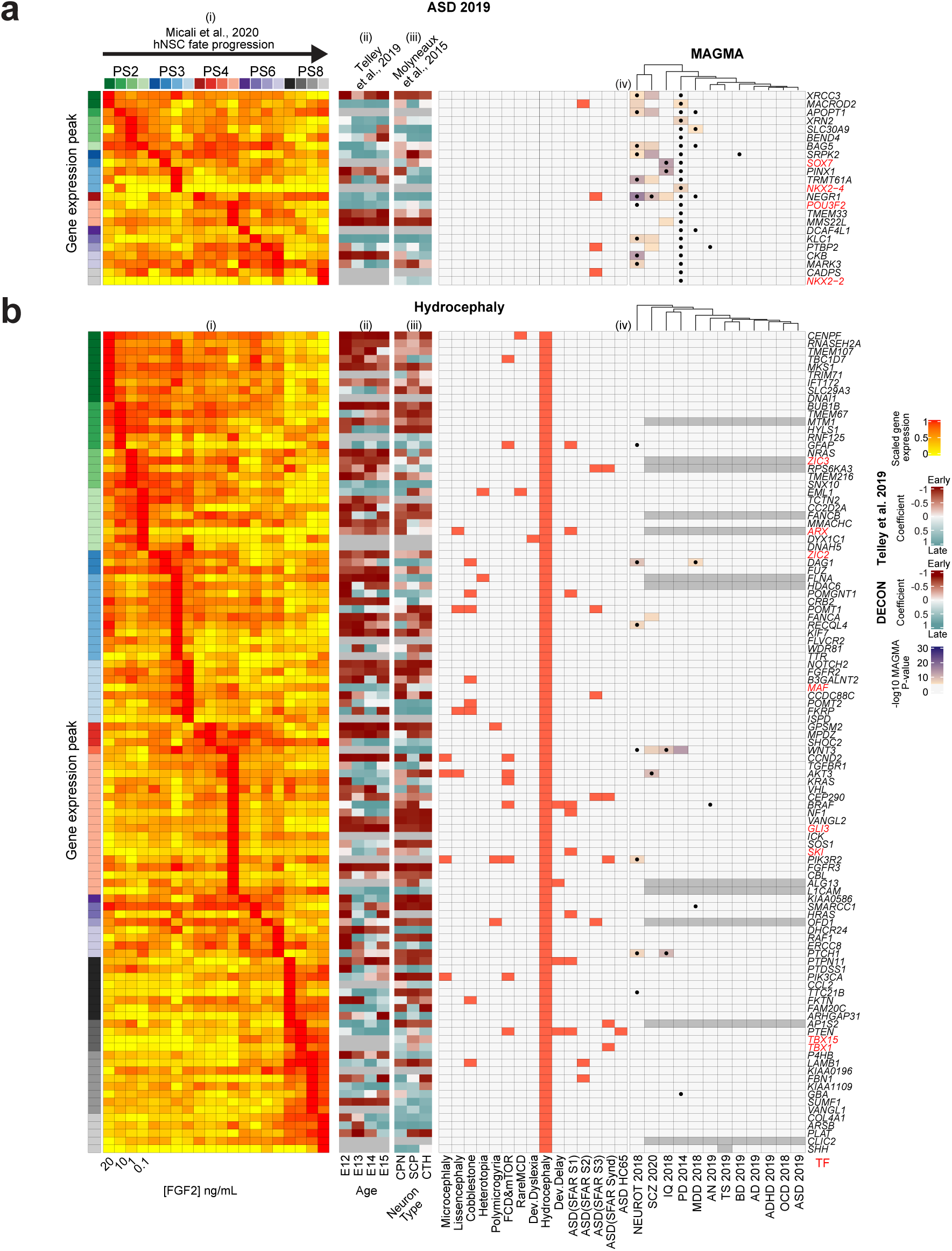
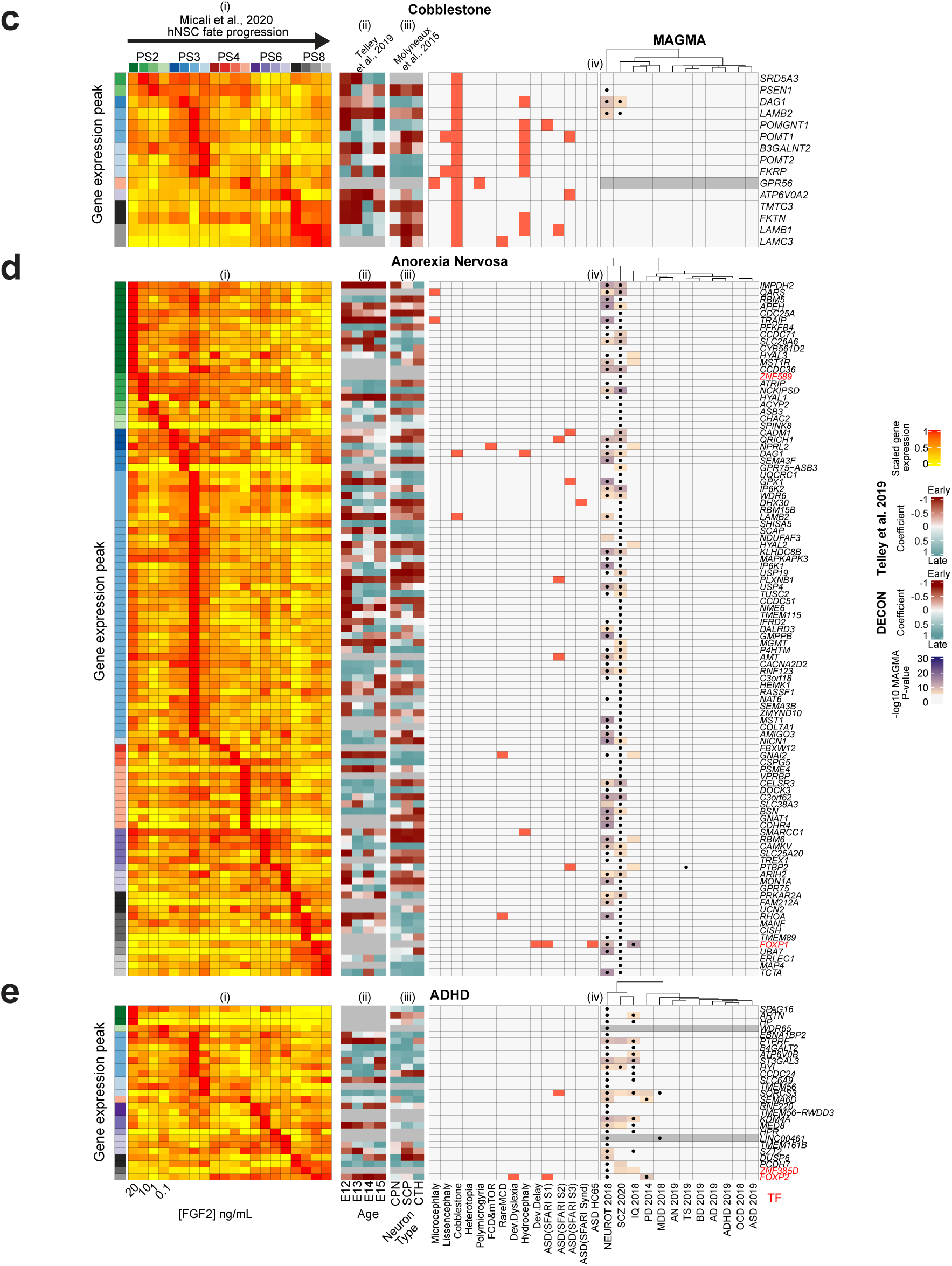
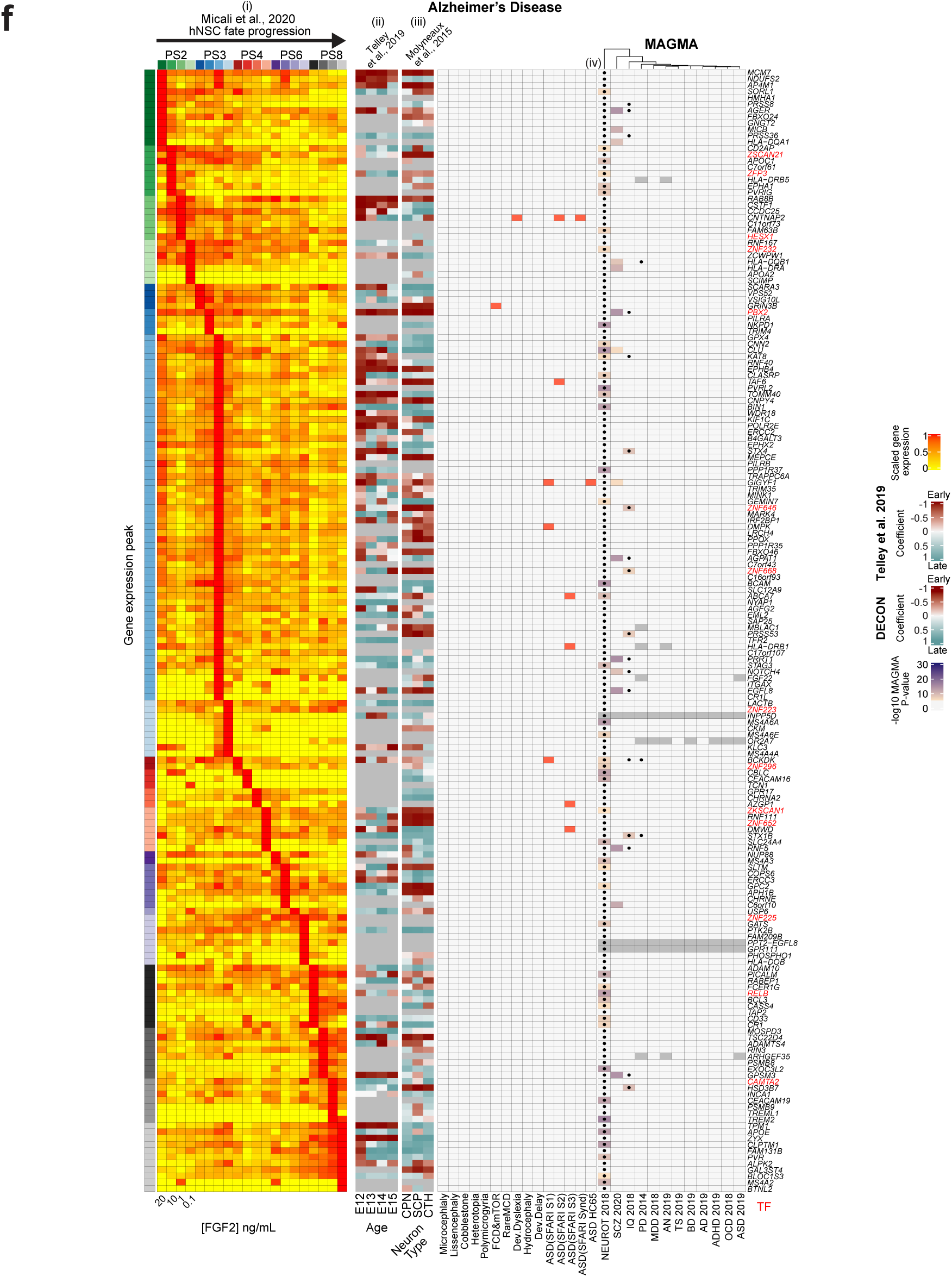
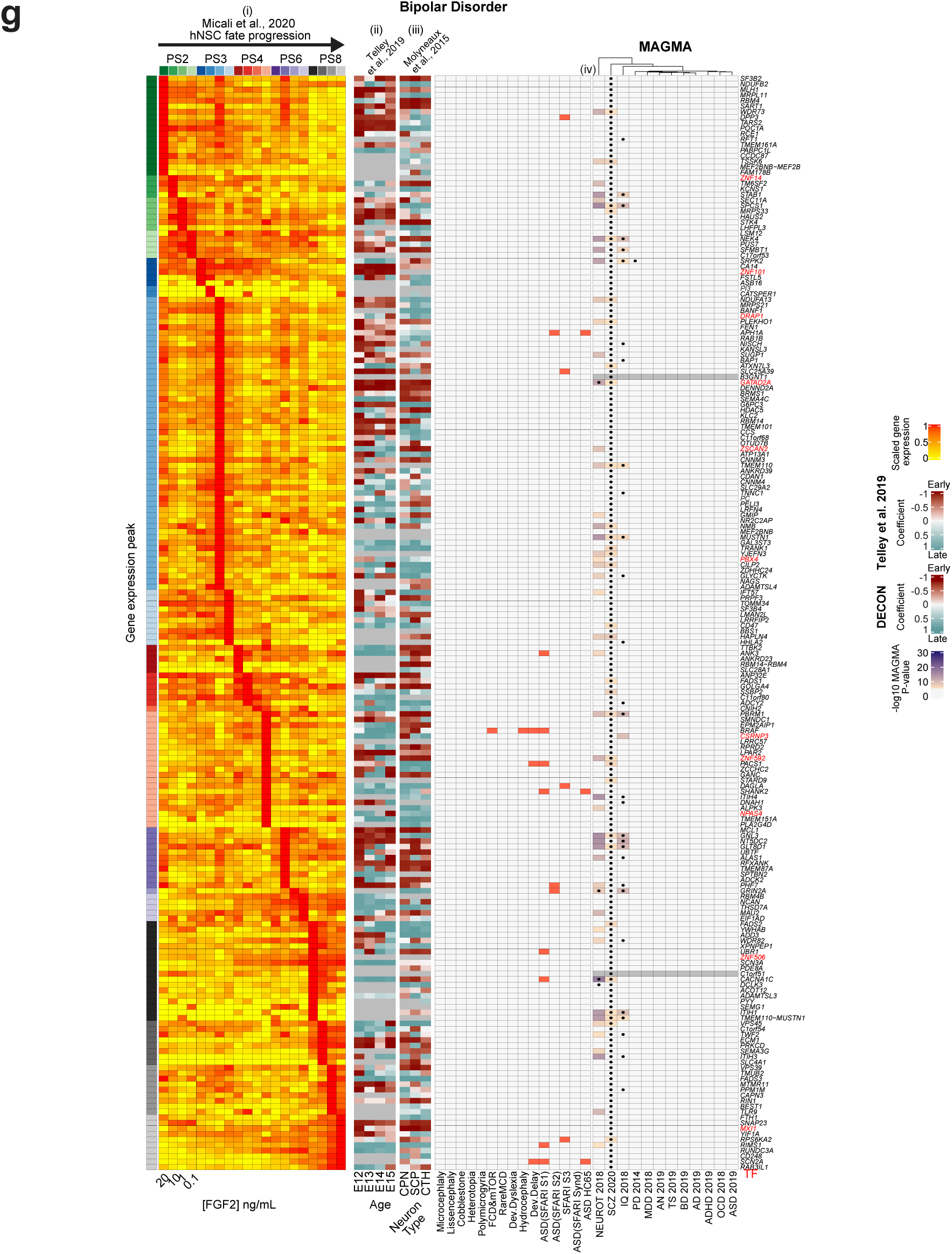
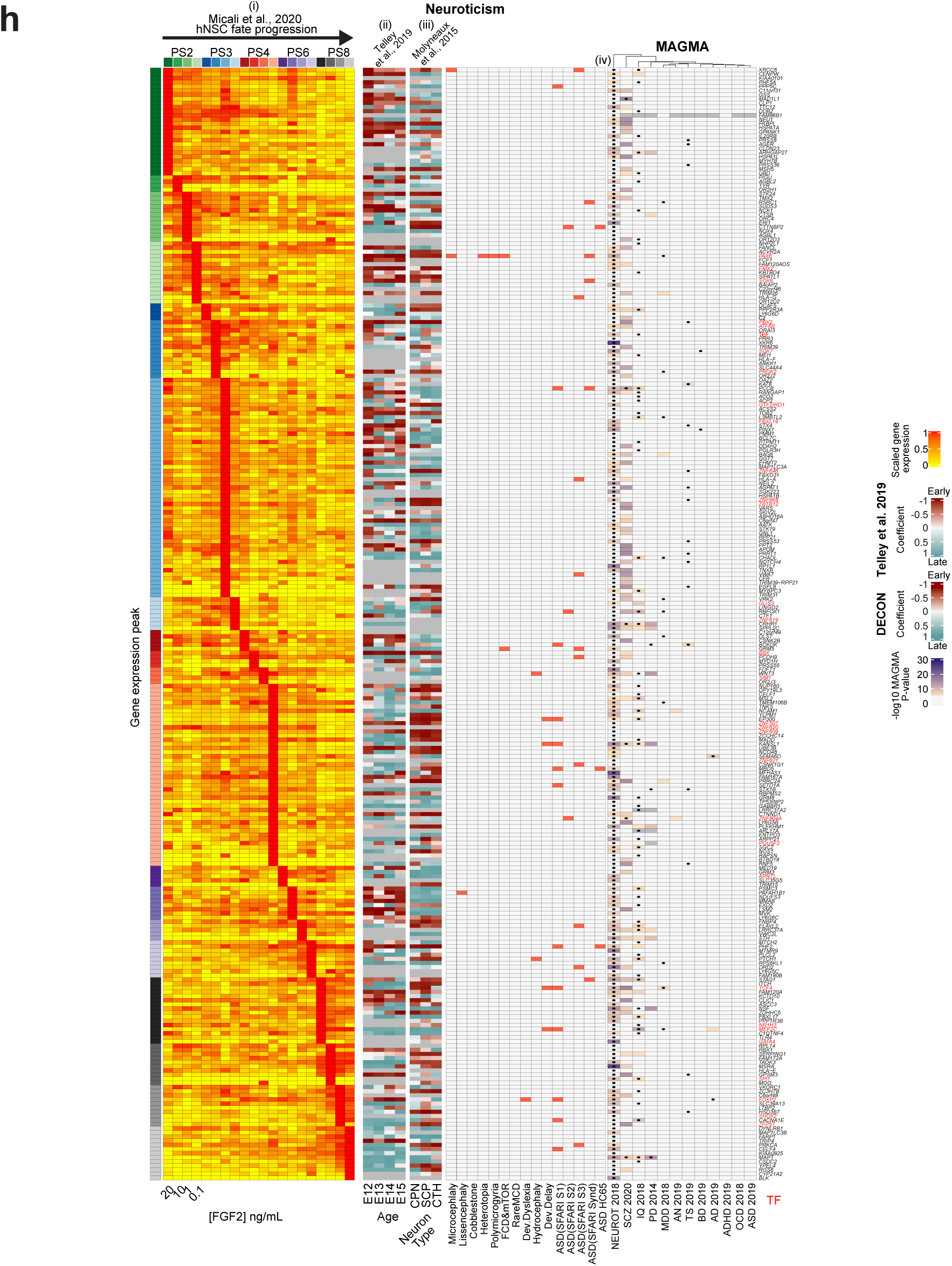
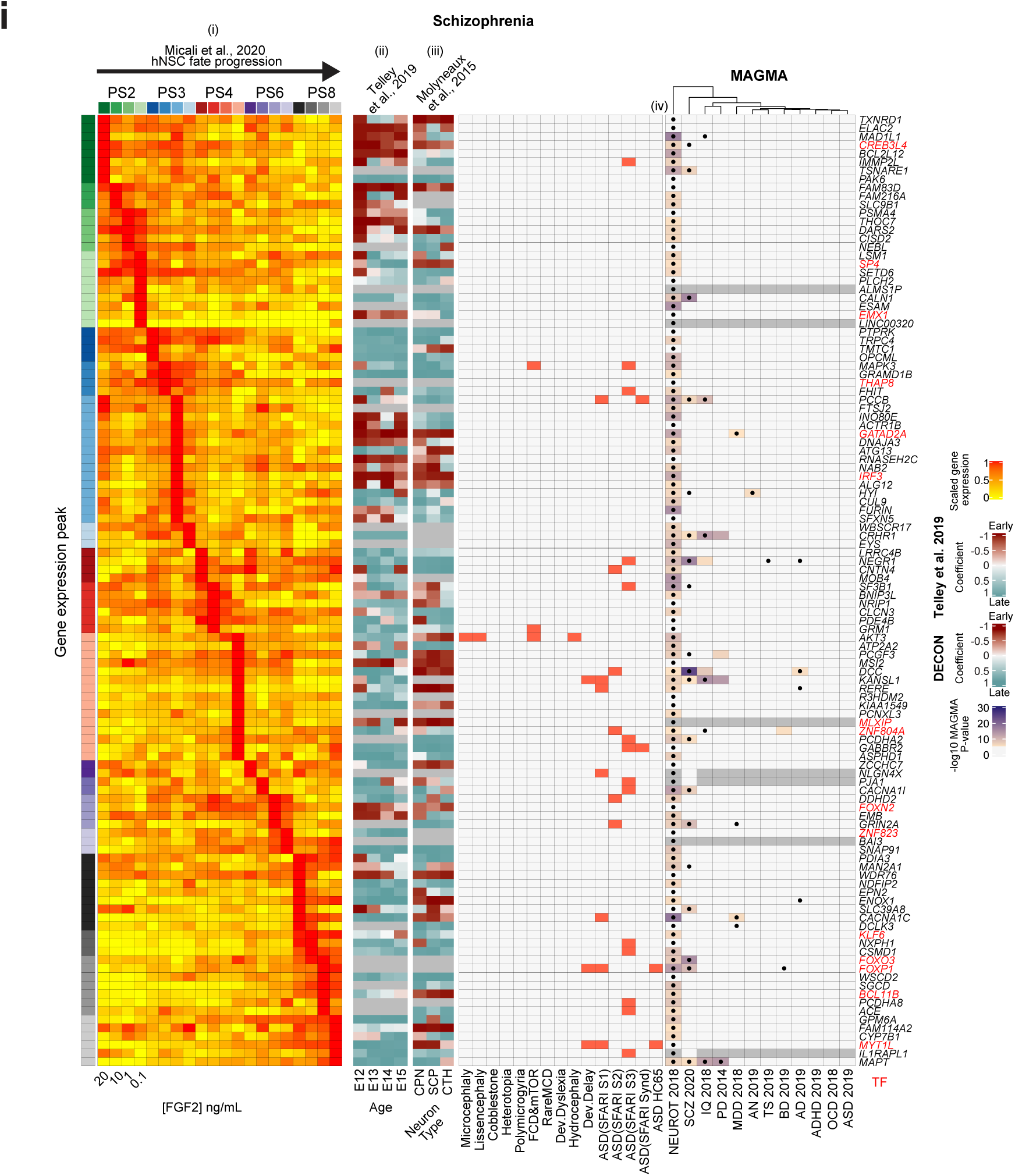
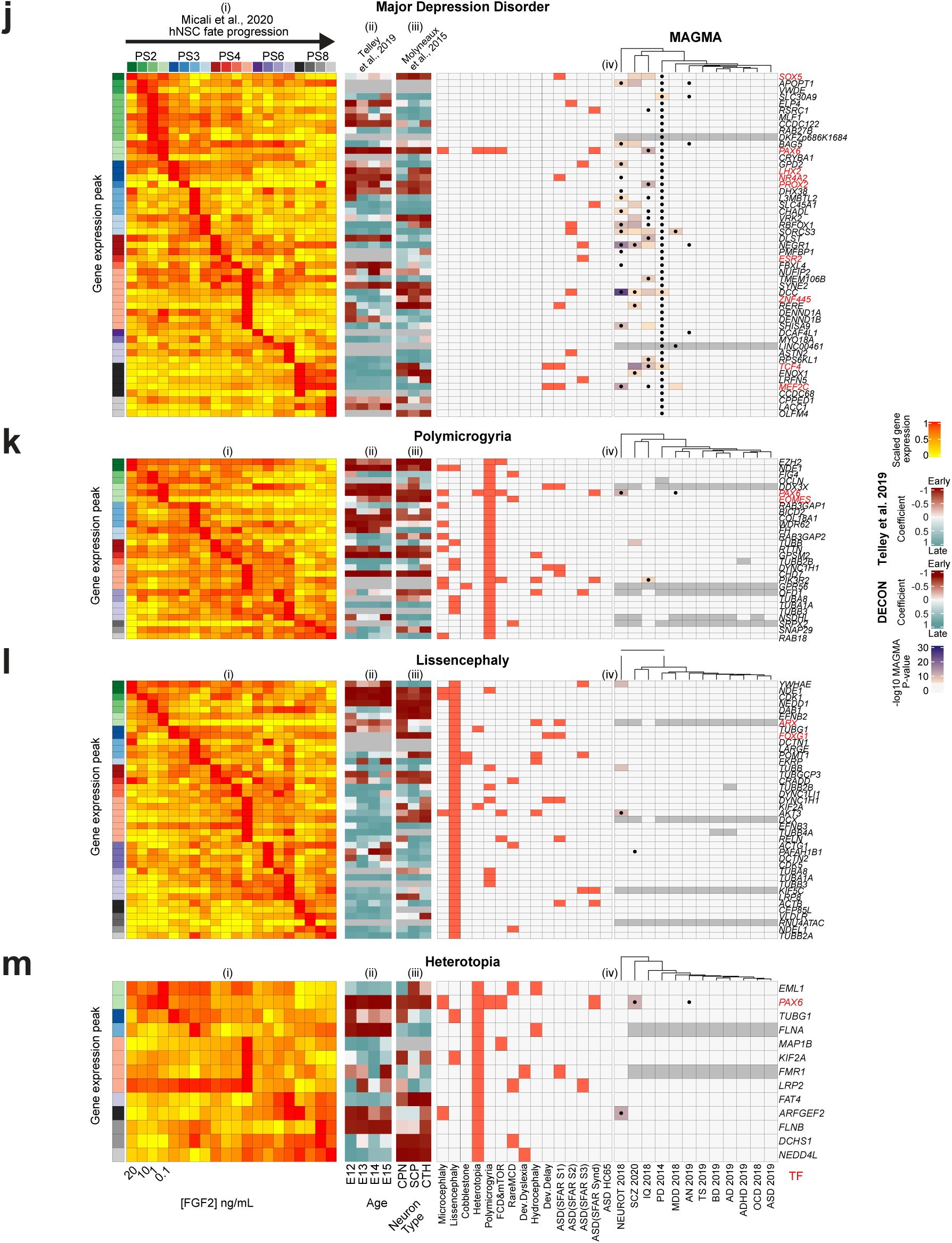
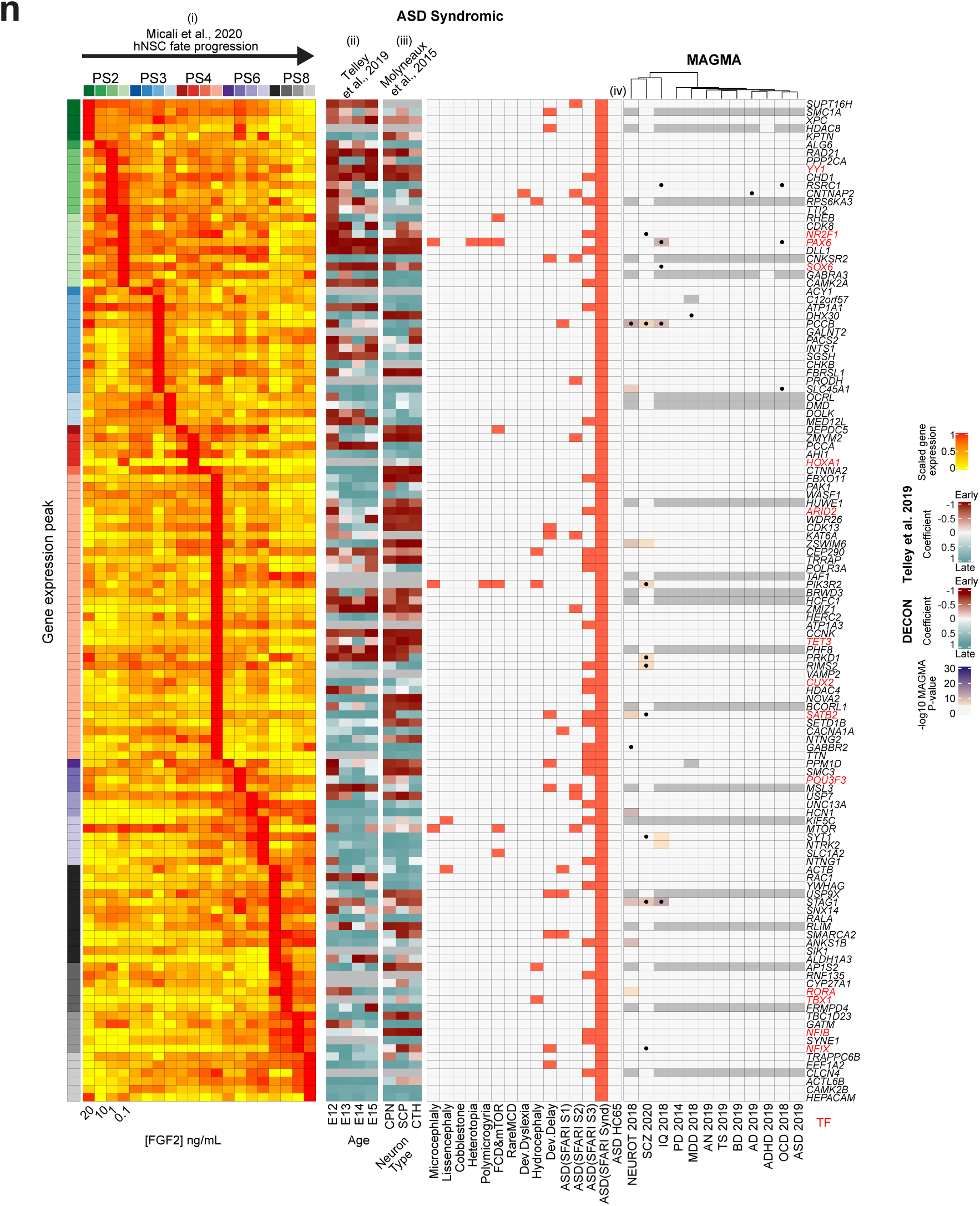
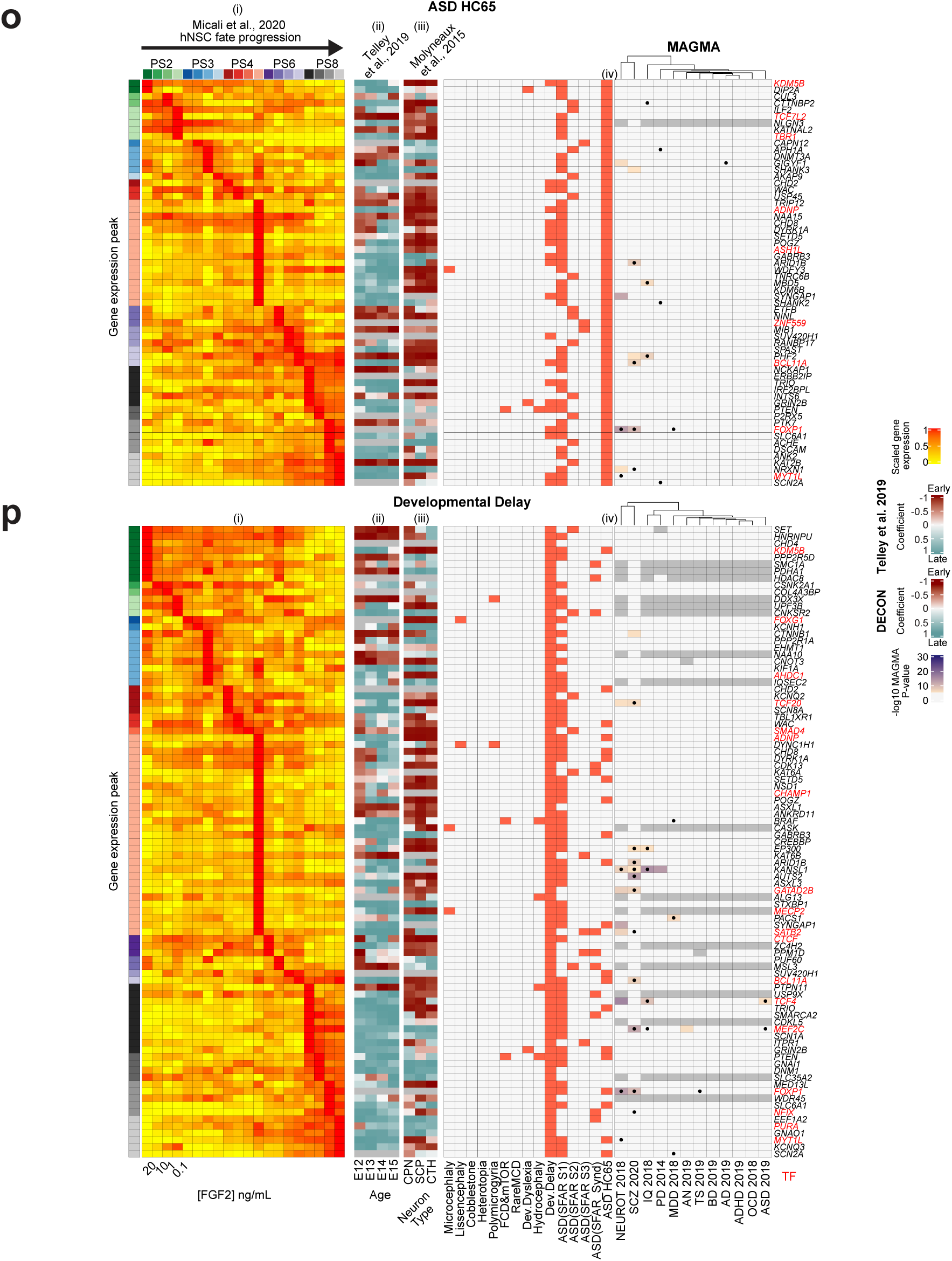
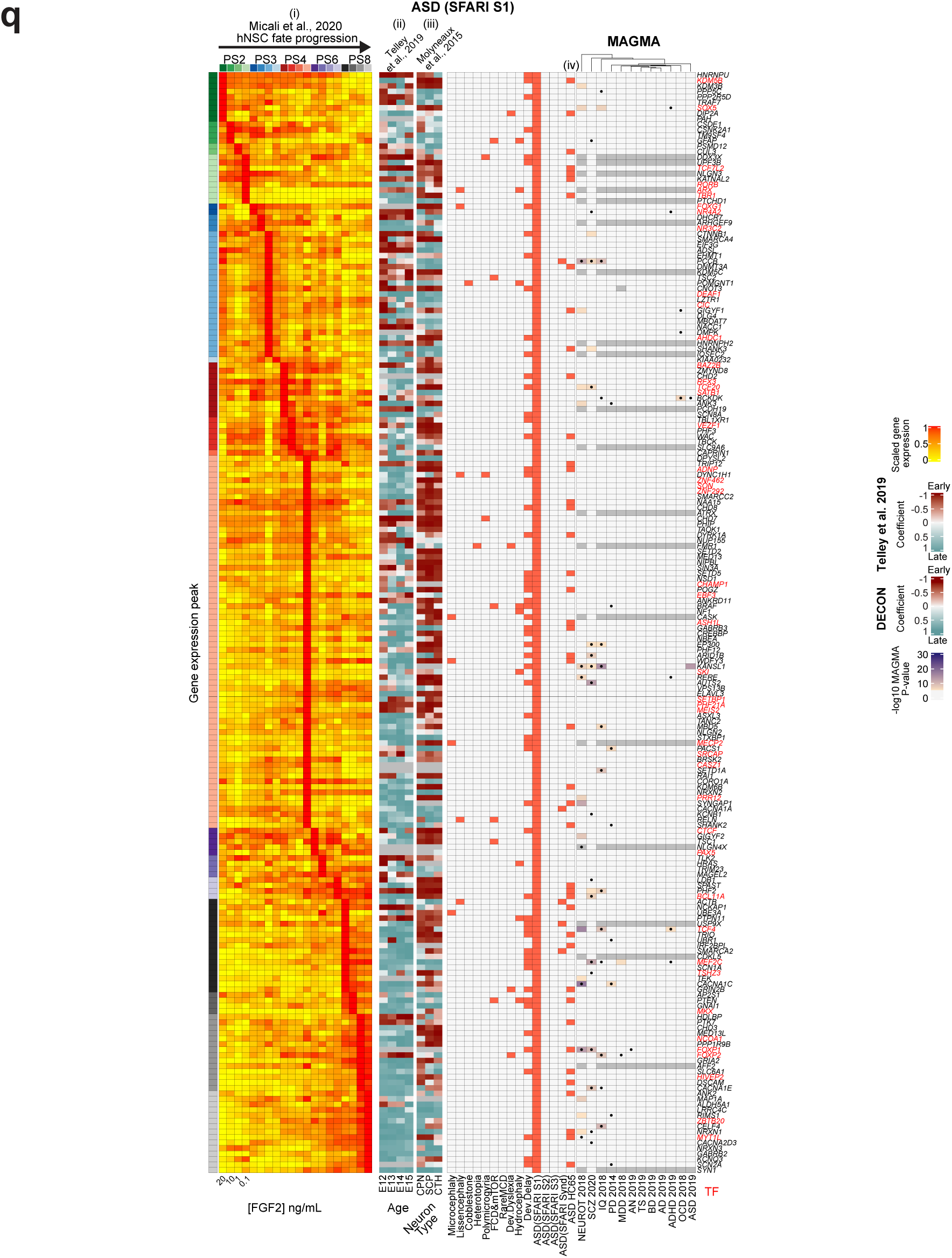
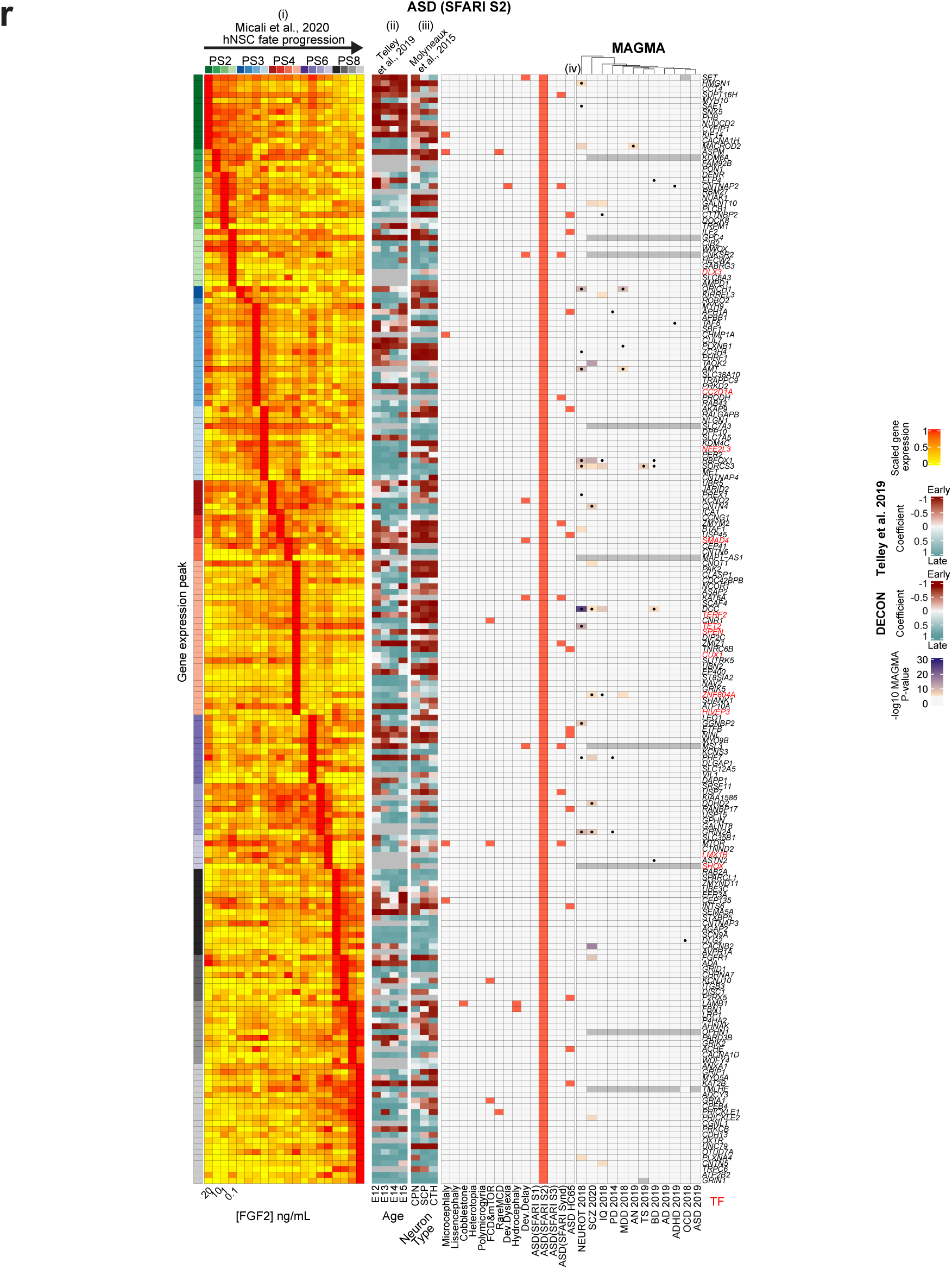
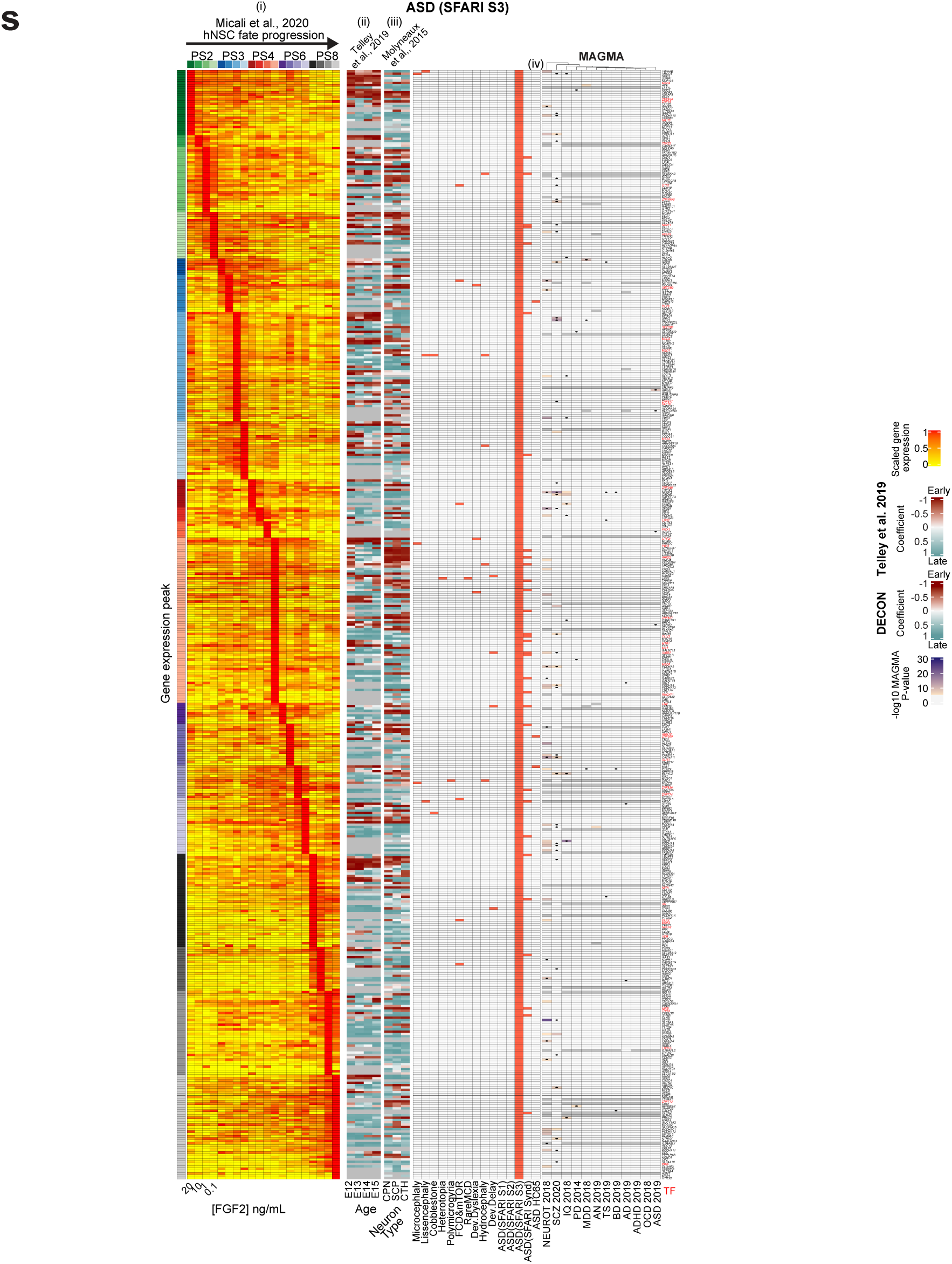
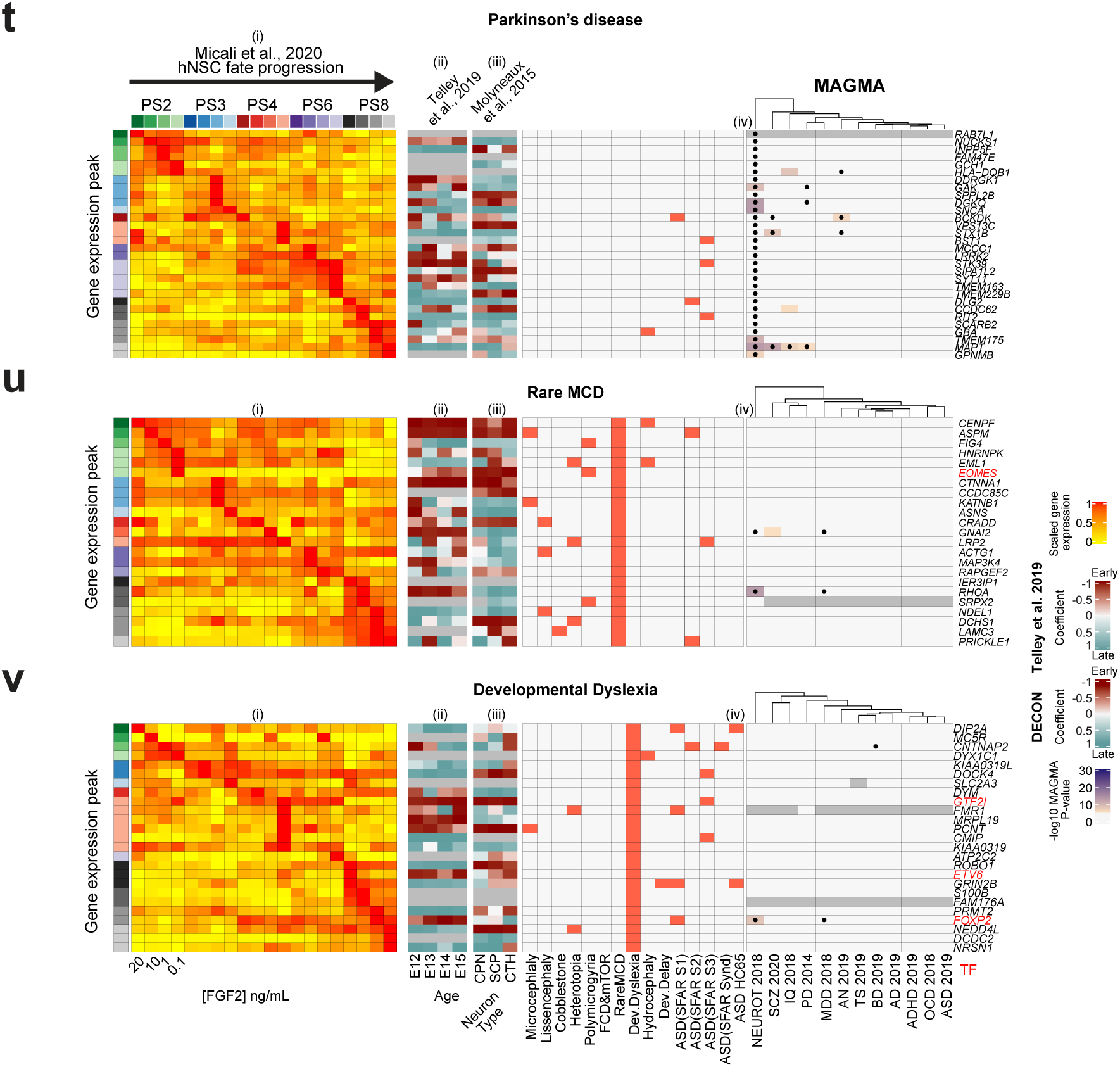
Expression dynamics of disease-associated risk genes across cortical neurogenesis. **a-v)** Multi-panel heatmaps displaying: (i) expression levels for each risk gene in the FGF2-regulated hNSC progression across passages ^40^. Genes are ordered by the temporal peak of expression. The distribution is represented by the left column colored by passage and FGF2 concentration. (ii) Temporal gene expression change across neuronal differentiation of age specific RG cells from developing mouse cortex ^56^. (iii) Temporal expression change across the maturation of the neurons from DeCoN dataset ^57^. (iv) Disease-gene association (left panel), and log10 P value of the MAGMA gene-level test of association with each GWAS dataset (right panel). Black dots indicate a top hit gene in the corresponding GWAS publication, based on genome-wide significant loci.

**Supplementary Fig. 4. Related to Fig. 1.**
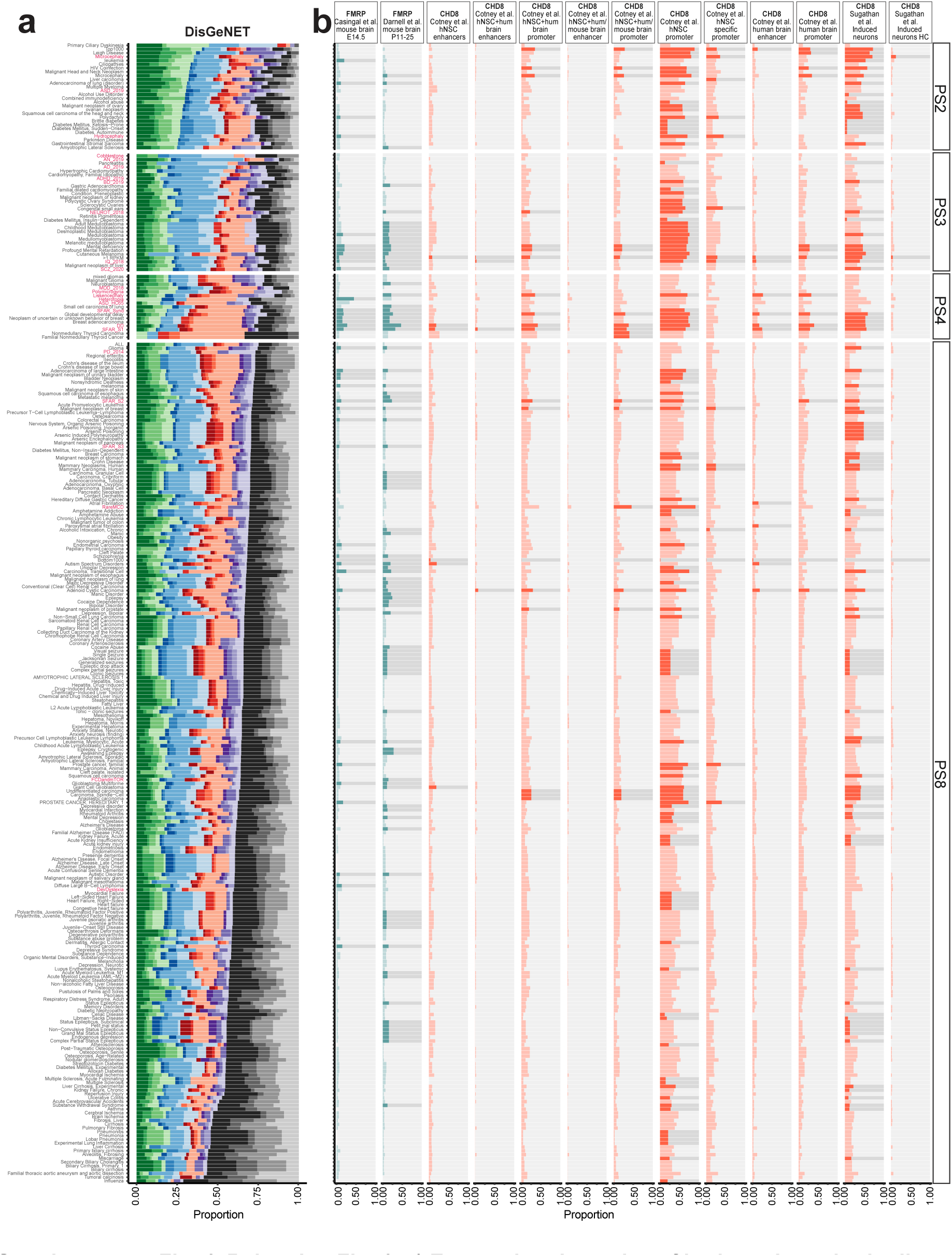
**a)** Expression dynamics of brain and non-brain disease-associated genes and FMRP and CHD8 targets across the progression of the hNSCs. Temporal distribution of the human disease-associated genes from DisGeNET across the FGF2-regulated progression of hNSCs ordered as: PS2 and PS3 high to low, PS4 and PS8 low to high. The majority of the cortical disorder-related gene sets resulted highly enriched at PS4 low FGF2, indicating that this pattern represents a neuronal specific signature which is not shared by non-brain diseases. Glioblastoma risk genes result enriched in the late RG cells of PS8, consistent with previous works ^37,78^. Moreover, certain cortical disorders and brain cancers might share similar NSC transcriptomic features. **b)** Barplot summarizing the gene proportion for each disease influenced by FMRP and CHD8. We tested the enrichment of FMRP and CHD8 targets derived from An et al., 2018, ^15^, Cotney et al., 2015 ^79^, Darnell et al., 2011 ^80^, Sugathan et al., 2014 ^81^, Casingal et al., 2020 ^82^, including embryonic and adult datasets, in our NDD gene sets and DisGeNET human diseases, across the progression of the in vitro hNSCs (Supplementary Table 2). FMRP targets derived from adult brain showed the highest overlap with all ASD gene sets, except for those GWAS-derived, and Dev.Delay-associated genes. Other diseases including FCD, SCZ, Neuroticism and IQ also showed a significant enrichment in specific sets of targets. LIS risk genes were enriched only in adult FMRP targets, while HET- or HC-associated genes were only enriched in embryonic FMRP targets. CHD8 targets also showed significant associations with non-GWAS ASD gene sets. CHD8 targets identified in NSCs showed significant overlap with cortical disorders caused by genes of earlier expression, for example MIC and several types of cancers. Genes associated with late onset diseases, such as AD and PD, were not enriched in any set of CHD8 and FMRP targets. These data indicate a transcriptional intersection of FMRP and CHD8 with their targets from the early phases of the NSC progression up to neurons. Besides replicating the expected association of ASD with CHD8 and FMRP targets, other cortical disorders also presented significant numbers of targets, with strong dependency on the tissue where they have been identified and developmental time. The findings suggest that the effects of the alteration of FMRP and CHD8 function extend beyond ASD and may implicate genes of earlier expression during telencephalic development involved in MCDs and other NDDs.

**Supplementary Fig. 5. Related to Fig. 2.**
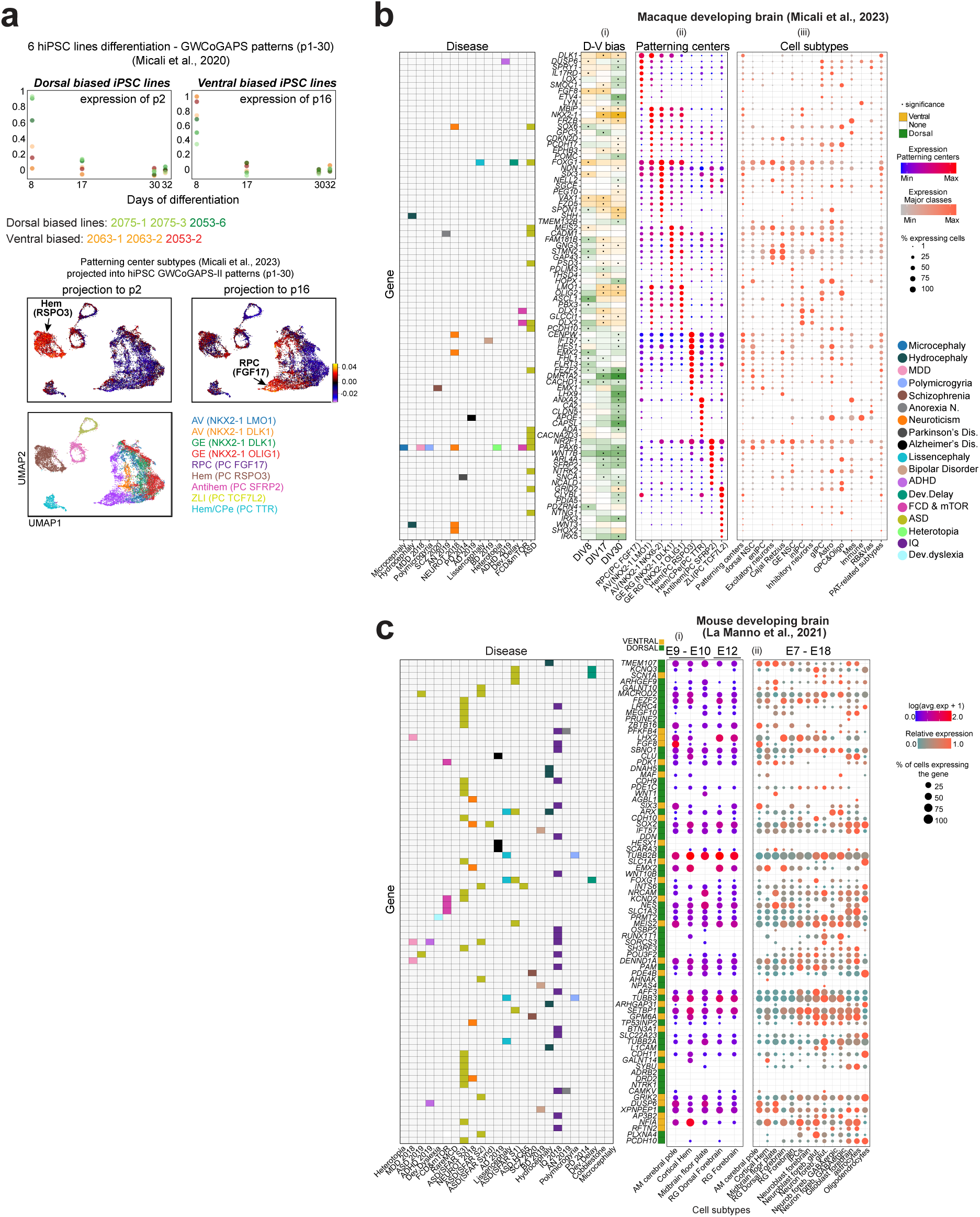
Expression of risk genes in telencephalic organizers. **a)** (Top panel) We have previously described 6 hiPSC lines (2063-1, -2; 2053-2, -6; 2075-1, -3) differentiated into forebrain NSCs. RNAseq data were generated across differentiation and decomposed with GWCoGAPS, which identified 30 patterns of gene expression (p1-30) ^40^. In Micali et al., 2020, GWCoGAPS and projection of bulk human developing cortex RNAseq data ^49^, distinguished 2 groups of cells: 2053-6 and 2075-1, -3, and 2063-1, -2 and 2053-2, involving dorsal or ventral telencephalic genes across their differentiation trajectory, respectively (Supplementary Fig. 20ci). GWCoGAPS patterns p2 and p16 showed differential expression in these 2 groups of lines at DIV8, as shown in top panel here. Patterns p2 was highly expressed in 2053-6 and 2075-1, -3 lines while patterns p16 was highly expressed in 2063-1, -2 and 2053-2 lines. The data in Micali et. al, 2020 indicate that cortical hem genes were highly weighted in p2, while anteroventral organizer genes were more represented in p16. Here, we further confirmed the differentiation bias of these 2 groups of cell lines by projection of scRNA-seq data from our recently reported developing macaque brain ^37^ into the GWCoGAPS patterns (p1-30). (Bottom panel) Projection of patterning center (PC) scRNA-seq data from developing macaque brain into the dorsal telencephalic pattern p2 and the ventral pattern p16 shows enrichment of p2 in monkey cortical hem cluster, and enrichment of p16 in rostral patterning center (RPC) cluster (<u>NeMO/CoGAPSII</u>). UMAP of all the PC clusters (bottom left panel). **b**) (i) Dorsoventral expression bias of PC marker genes from macaque brain single-cell data ^37^ in NSC lines at DIV 8-30, regardless of disease association and bias significance; gene expression of the same PC markers in (ii) clusters annotated as PCs [RPC (PC FGF17), AV (PC NKX2-1 LMO1), AV (PC NKX2-1 NKX6-1), GE (RG NKX2-1 DLK1), GE (NKX2-1 OLIG1), hem (PC RSPO3), hem/CPe, (PC TTR), antihem (PC SFRP2), and ZLI (PC TCF7L2)], and in (iii) other cell subtypes from the same macaque dataset. Only filtered PC marker genes are displayed, using no more than 15 genes per cell cluster, based on the lowest p-value from the original data ^37^ (see methods). Gene-disease association on the left. **c**) Dot plots representing risk gene expression in the cell clusters annotated as (i) patterning centers (anteromedial pole, cortical hem and floor plate) and forebrain RGs, and (ii) other neural cell types at different maturation phases from mouse fetal brain single-cell data ^58^. Only disease genes with significant dorsoventral expression differences in the 6 hNSC lines at DIV 8 are displayed. Significance in dorsoventral bias is indicated in the left colored column. Gene-disease association on the left. A-M: Antero-medial cerebral pole.

**Supplementary Fig. 6. Related to Fig. 3.**
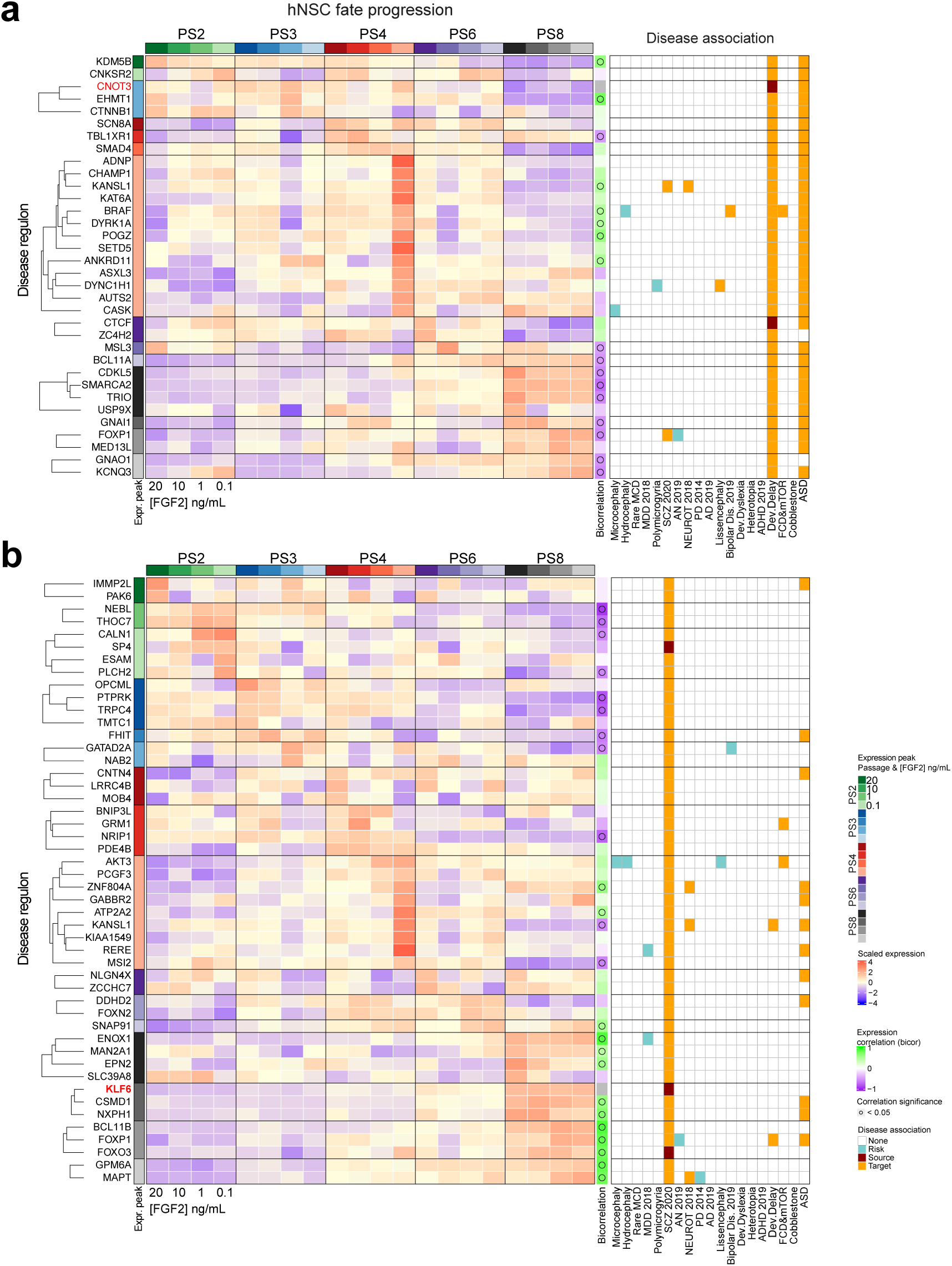
Expression dynamics of disease-regulon members across hNSC progression. (**a and b**) Expression of core TFs (in red) and target genes of 2 representative regulons, one early (CNOT3) and one late (KLF6), across the progressing hNSCs. The “Peak Sample” column represents the peak of gene expression. The “Bicorrelation” column shows the correlation between the expression of any target gene and the core TF. The grid on the right indicates gene-disease association, specifying core TFs and target genes in the disease regulons. Each core TF may have a differential expression level and functionality along cortical development. Their target genes can be expressed at multiple phases across the NSC progression diverging from the core TF expression peak. This uncoupling might reflect multiple TF-effector mechanisms (e.g. co-factors’ stoichiometry and their combinations, post-translational modifications and subcellular localizations, etc.), a negative regulatory effect, or unspecific associations between TFs and their motifs due to motif similarity.

**Supplementary Fig. 7. Related to Fig. 3.**
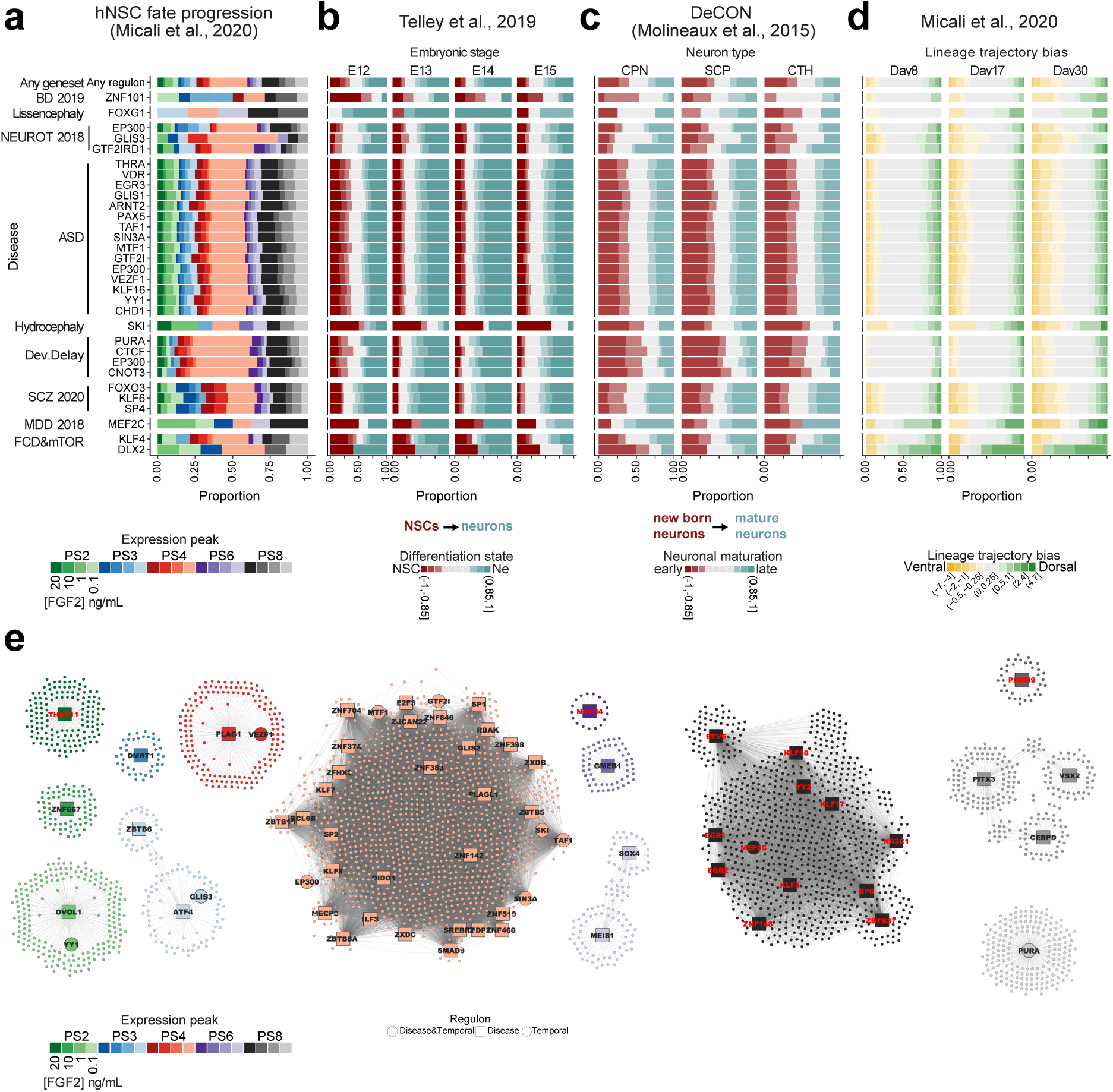
Disease-regulon genes expressed in vivo and in vitro. Proportion of genes for each disease regulon **a**) sorted by passages and FGF2 conditions according to their maximum expression as explained in Fig. 3a; **b**) classified into different bins of expression fold change across NSC differentiation from ^56^; **c**) classified into different bins of expression fold change across the maturation of the neurons from ^57^; **d**) classified into different bins of expression fold change in dorsoventral trajectory bias from ^40^. **e**) Temporal regulons across passages. Small nodes represent target genes, and bigger, labelled nodes represent core TFs of temporal regulons (squares), disease regulons (circles) or both (octagons).

**Supplementary Fig. 8. Related to Fig. 3.**
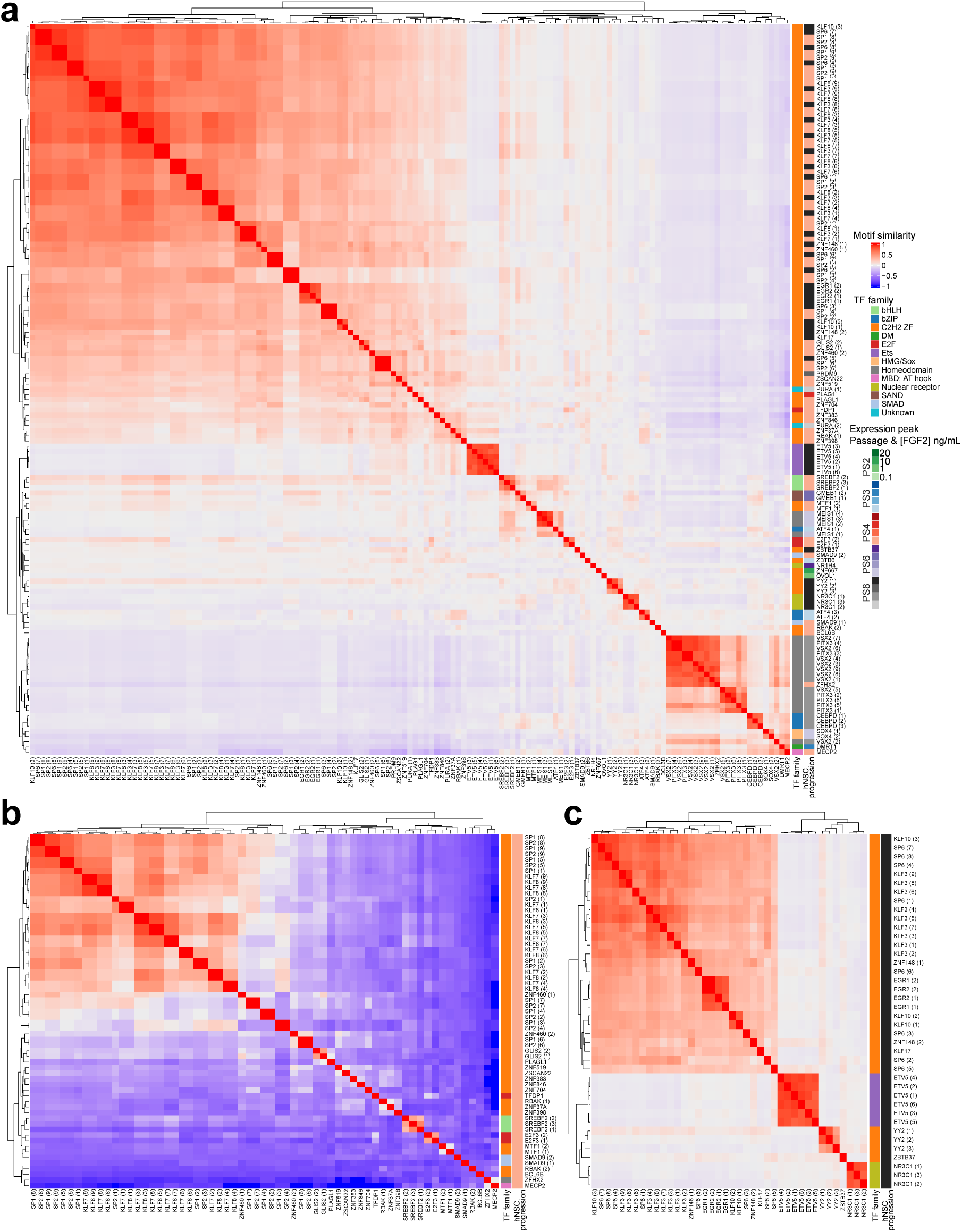
Motif similarity of core TFs in temporal regulons. (**a-c**) Similarity of all motifs associated with core TFs of regulons found in the progression of hNSCs. a) All regulons. b) Regulons from Passage 4, 0.1 ng/mL FGF2. c) Regulons from Passage 8, 20 ng/mL FGF2. Genes associated with multiple motifs are identified in parenthesis. The passage and FGF2 concentration of maximum expression is indicated in the right column “hNSC progression”. Motif similarity and TF-motif association data are from Lambert et al., 2018 ^83^.

**Supplementary Fig. 9. Related to Fig. 4.**
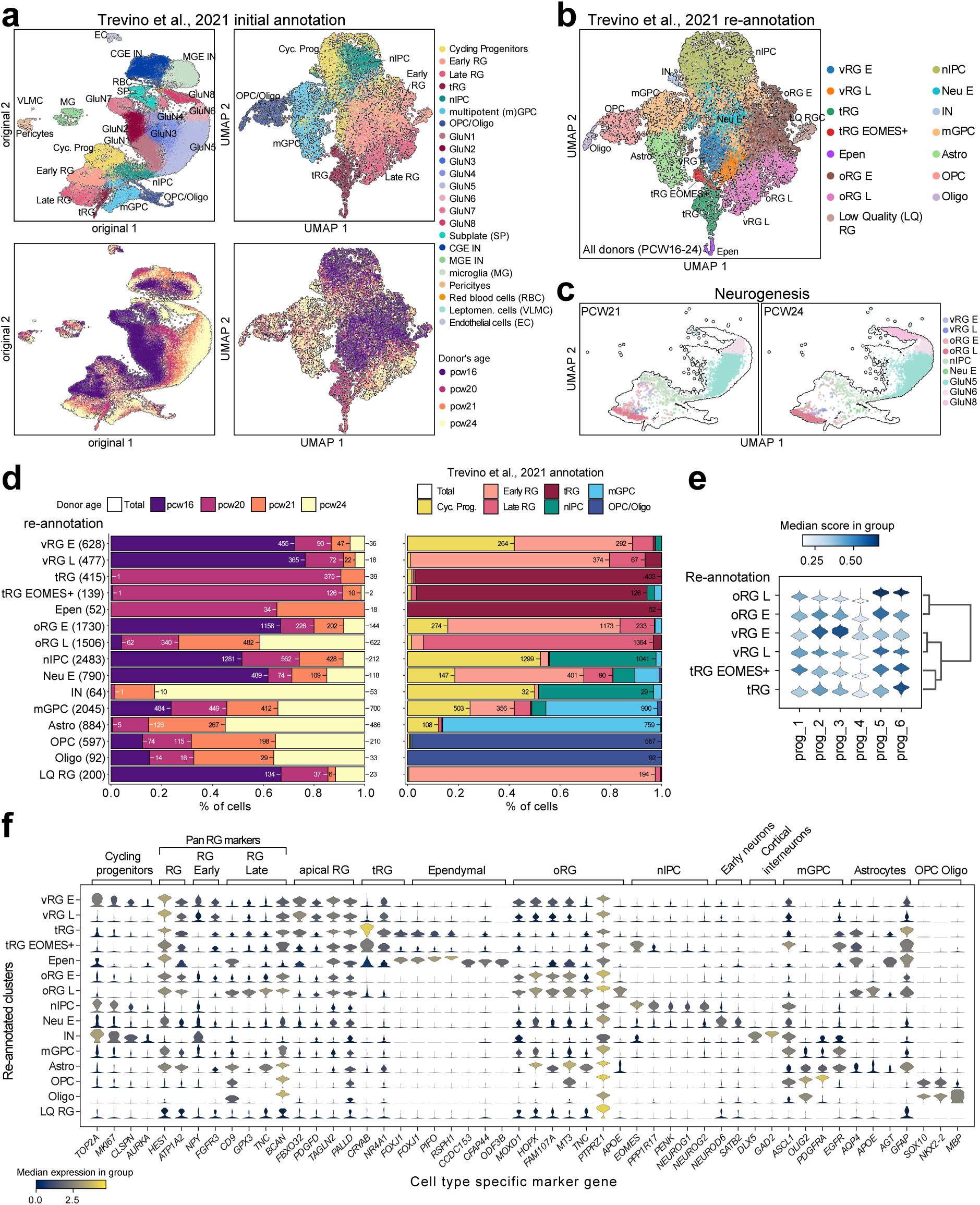
Re-annotation of single cells from human fetal brain data. **(a)** UMAP representation of scRNA-seq data from developing human brain ^61^. All cells in the study are represented (left). A subset of progenitor cells of interest for this study was reannotated (right). Colors show the original cell annotation (top) and the age of the donors (bottom). **b)** UMAP colored by the new annotation of the subset of cells in this study. **c)** Neurogenesis trajectories in Trevino et al., 2021 analyzed in CellOracle. PCW20, PCW21 and PCW24 donors were independently considered. PCW16 donor was not included in the neurogenic lineage since we could not obtain a clear trajectory from NSCs towards differentiated cells in the cell diffusion maps. For the maturation of RGs and gliogenesis, all donors were included. **d)** Size, donor and distribution of the new cell identities. The total number of cells is shown next to the cell type nomenclature. The barplots represent cell distribution in the donors (left) and the distribution of cell types from the original annotation (right) present in the new cell groups. **e)** Expression scores of genes associated with the progression of RG cells. Marker genes associated with the progression of RG cells from Telley et al., 2019 ^56^ were used to establish the early/late subclasses of reannotated RG cells in Trevino et al., 2021. The gene module “prog_1” includes genes with both early and late expression; gene modules “prog_2” to “prog_6” classify genes according to their early-to-late expression bias. Genes in “prog_2” and “prog_3” have an early expression pattern, while “prog_5” and “prog_6” include genes with a late expression pattern. We observed early and late gene signatures enriched in specific vRG and oRG clusters which we use to label early and late vRG and early and late oRG clusters. **f)** Gene expression of cell type markers in re-annotated cell subtypes. Violin plots representing expression normalized by total counts per cell, in log-scale. The color represents the median gene expression in that group of cells.

**Supplementary Fig. 10. Related to Fig. 4.**
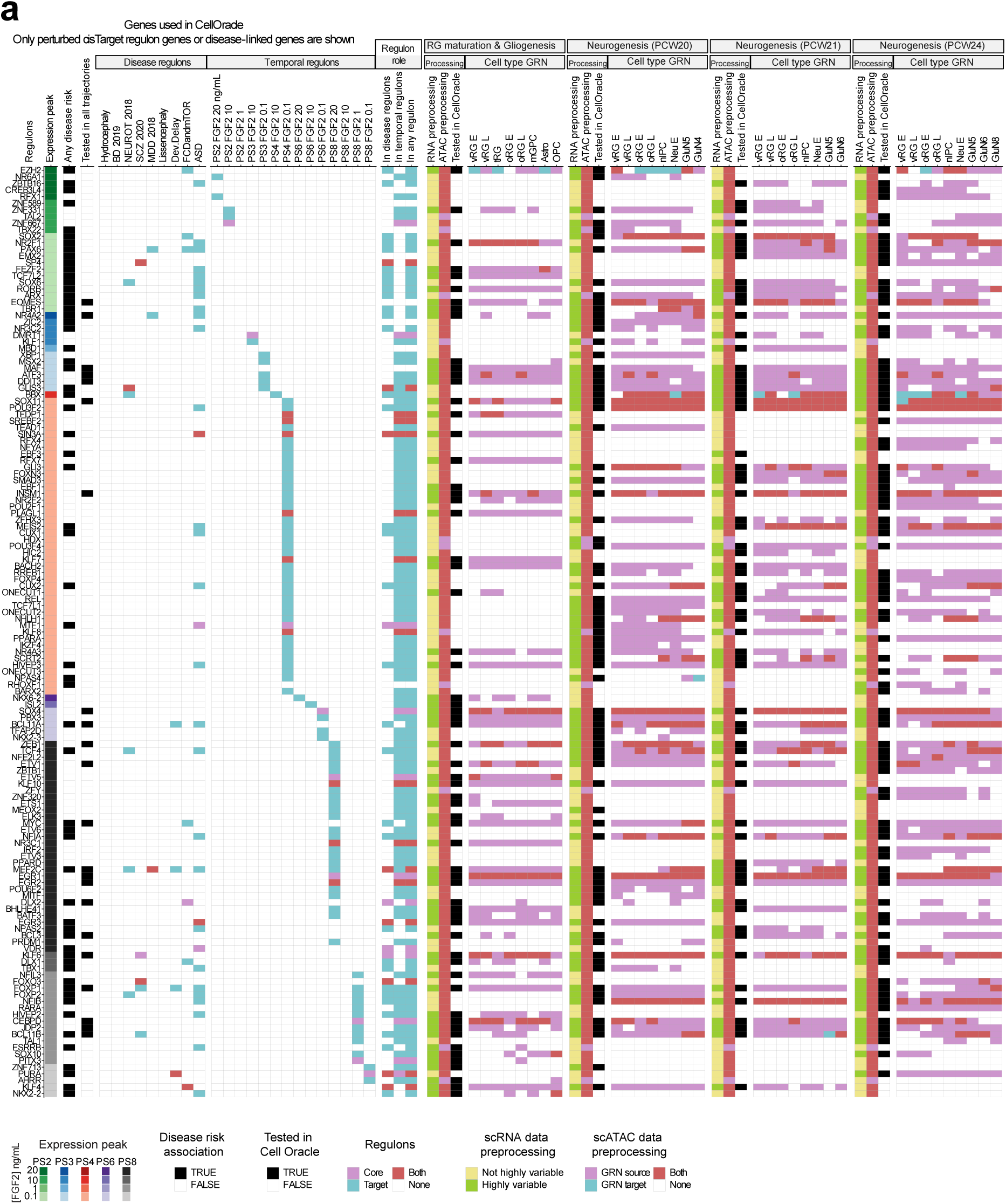
Overview of the genes tested in CellOracle associated with any disease or regulon. **a)** From the left to the right column: phase of peak expression across hNSC progression at different passage and FGF2 condition from Micali et al., 2020 i; association to any disease; test of a gene in CellOracle in both neurogenesis and gliogenesis. The columns under disease-regulons and temporal-regulons show if a gene has been identified as core TF or target gene in the regulons from Fig. 3, derived from disease-associations and from the hNSC progression, respectively. These are summarized in the ‘Regulon role’ columns. For each subset, it is also shown if a gene passed preprocessing in scRNA-seq data and scATAC-Seq data, if the knock-out (KO) simulation succeeded, and the role (source gene or target gene) in each cell type’s GRN (subsetted to the top 2000 regulatory connections in the network). ScRNA-seq preprocessing included removing low-expression genes and considering only the top 3000 highly variable genes (see methods). ScATAC-seq preprocessing included the generation of a base GRN using Cicero ^84^ to find coaccessible chromatin regions and a motif-based filtering to recover potentially regulatory links between these regions. For every trajectory, the gene expression data is modelled using the GRN derived from the corresponding scATAC-seq data. Expression is modeled individually in each cell type, resulting in a cell type-specific model of gene expression that connects TFs and target genes.

**Supplementary Fig. 11. Related to Fig. 4.**
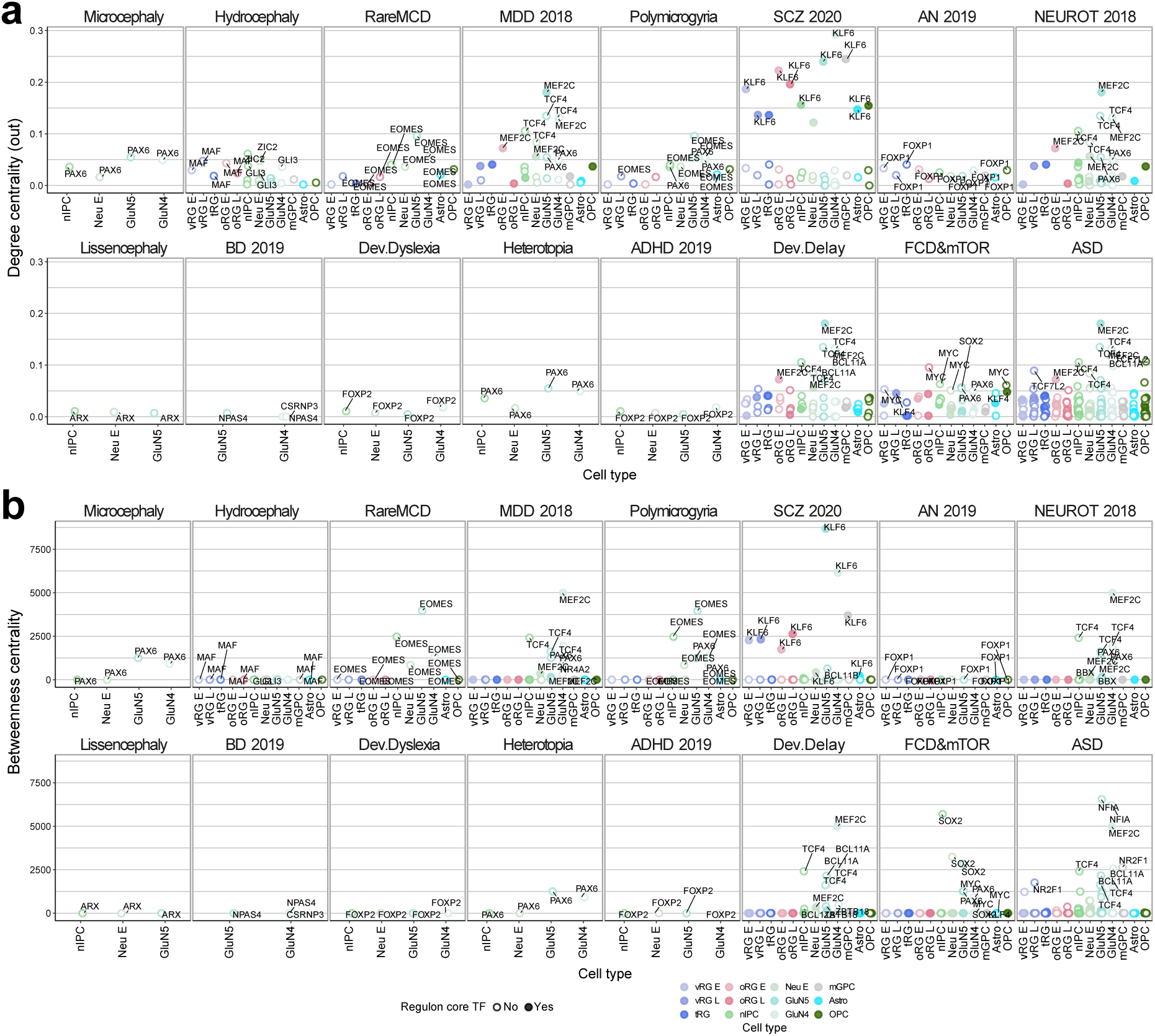
Network scores for the disease-associated TFs. The columns split the genes based on the associated diseases. The colored dots on the x axis represent the cell types with a score value for the genes on the y axis: **a**) degree centrality (out connections) represents the number of genes regulated by a given TF relative to the network size; **b**) betweenness centrality represents the importance of a given TF for connecting any two genes in the network. For genes and cell types measured in both trajectories (RG maturation and gliogenesis, and neurogenesis), data from RG maturation and gliogenesis are shown.

**Supplementary Fig. 12. Related to Fig. 4.**
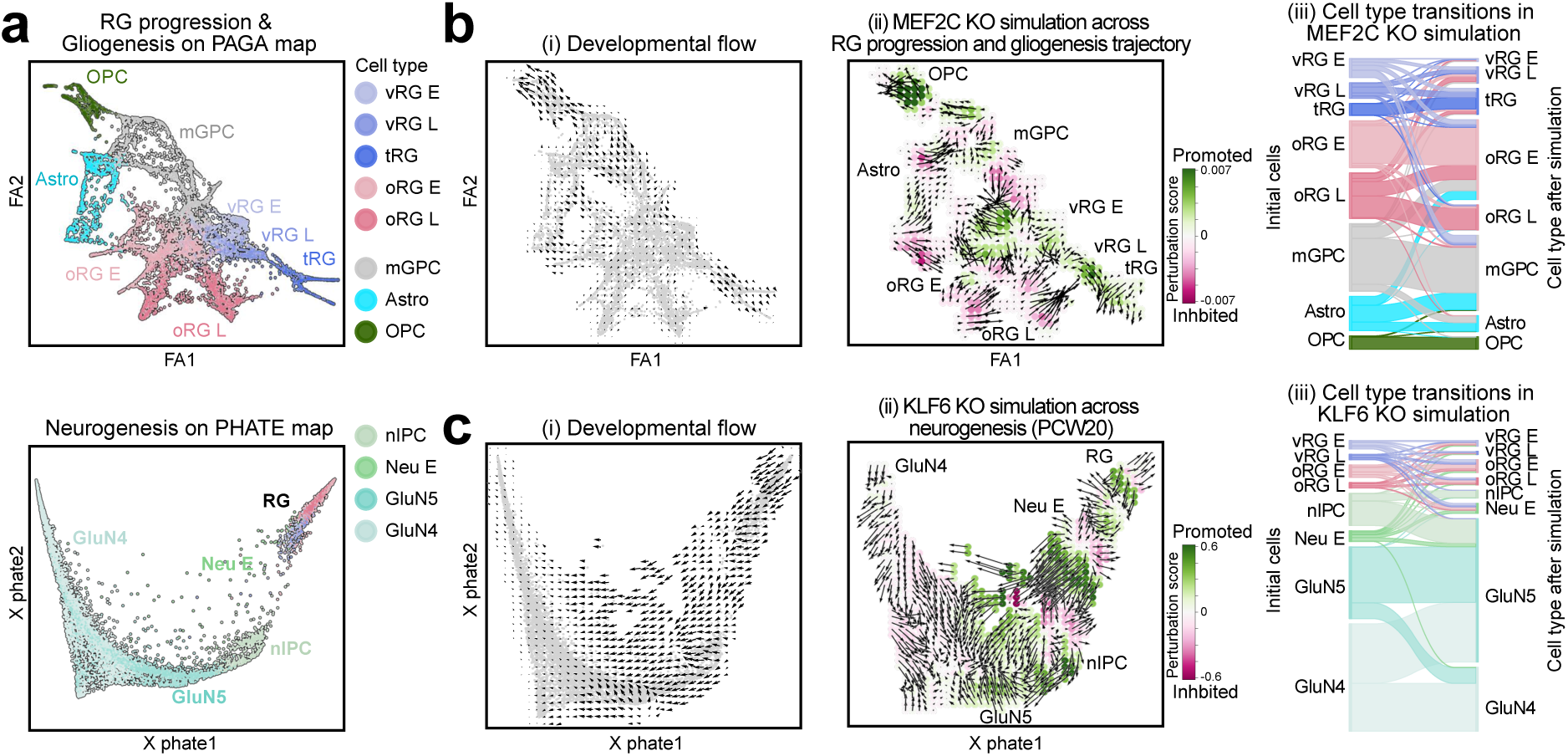
CellOracle KO simulation of core TFs across RG maturation/gliogenesis and neurogenesis. **a)** PAGA map of RG maturation and gliogenesis (top), and PHATE map of neurogenesis (bottom). **b and c)** (i) Developmental flow of (b) RG maturation and gliogenesis and (c) neurogenesis. Arrows represent the direction of the progression in the map. ii) KO simulation for *MEF2C* and *KLF6* in RG maturation/gliogenesis and neurogenesis trajectory perturbation, respectively. Arrows show the perturbation of the cell flow that the KO simulation produces, and color represents the flow direction change upon perturbation (green means same direction, i.e., promoted trajectory, and red means opposite direction, i.e., depleted trajectory). iii) Sankey plot representing the cell transitions observed in the perturbation. Original cell identities (left axis) and after KO simulation (right axis) are shown.

**Supplementary Fig. 13. Related to Fig. 4.**
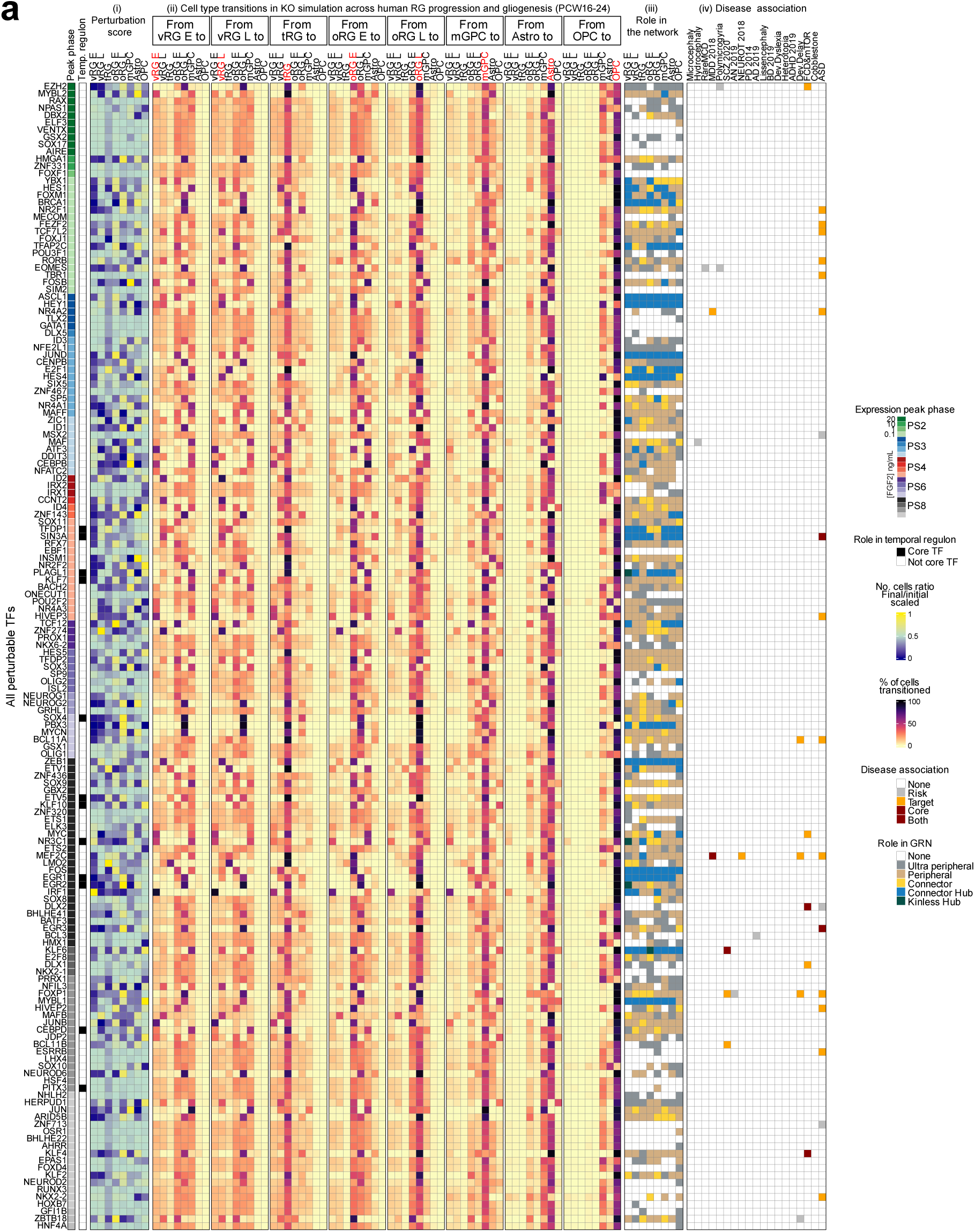
Comprehensive TF KO simulation across RG maturation and gliogenesis (PCW16-24). **a)** The peak expression phase across the in vitro hNSC progression from Micali et al., 2020 is shown for all perturbable TFs on the left column. i) Perturbation score indicating gaining or depletion of a given cell type, measured as the ratio of cells with an identity before and after the perturbation. Values are normalized per cell type. ii) Cell type transitions upon KO simulation. The first bar of the top axis represents the original cell type, and the second bar represents the cell identity resulting from the simulation. The grids represent the fraction of the original cell type (labelled in red) and their final identity. iii) Regulatory role of every gene in each cell type. TFs connecting different GRN modules are considered hubs, and they are ordered according to their connectivity to local modules: from ‘Ultra peripheral’ to ‘Kinless’ for non-hubs, and from ‘Provincial’ to ‘Kinless’ for hub TFs. iv) Disease association of every gene. TF association to disease can be risk-only, target of a disease-regulon and core of a disease-regulon.

**Supplementary Fig. 14. Related to Fig. 4.**
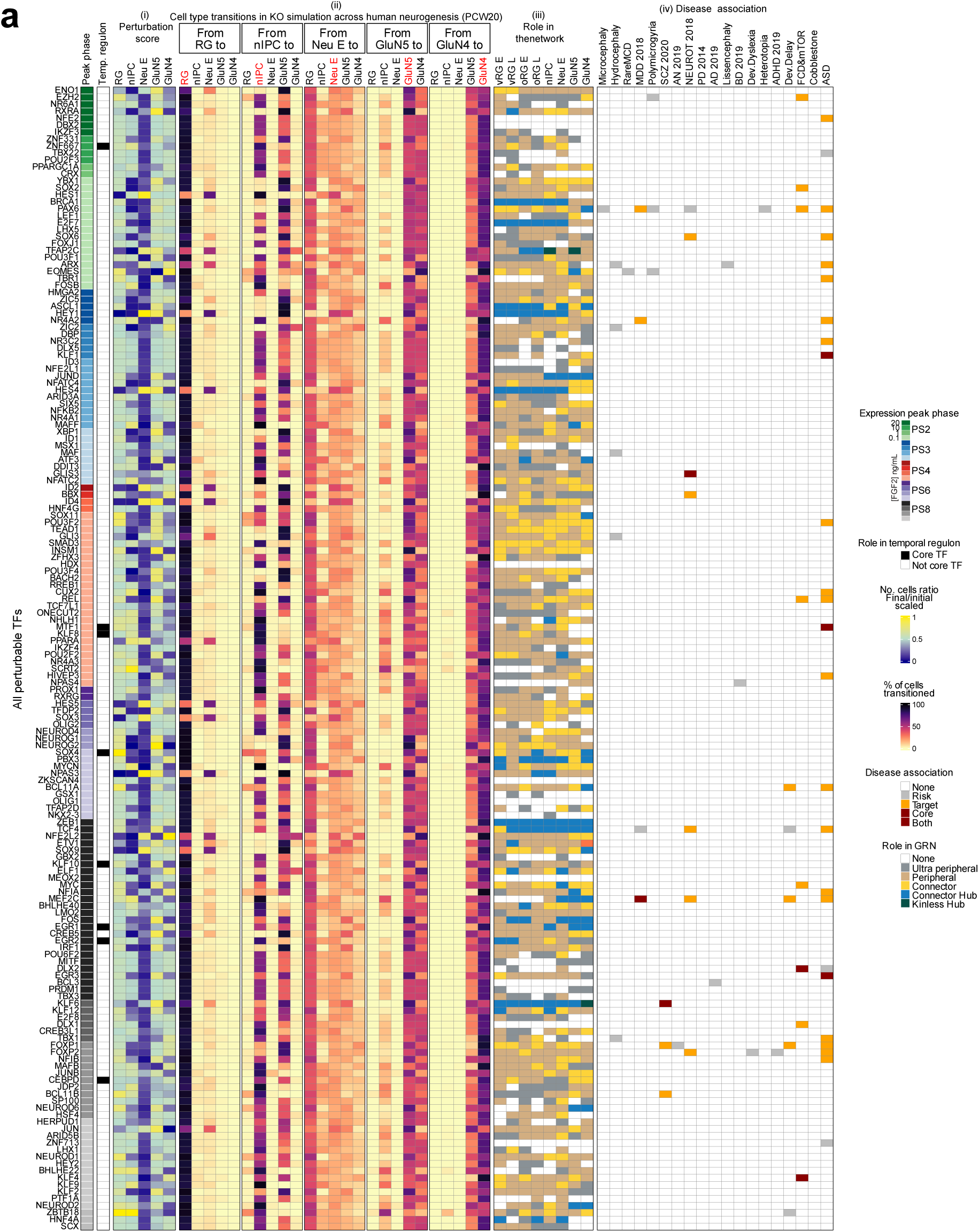
Comprehensive TF KO simulation across neurogenesis (PCW20). **a)** The peak expression phase across the in vitro hNSC progression from Micali et al., 2020 is shown for all perturbable TFs on the left column. i) Perturbation score indicating gaining or depletion of a given cell type, measured as the ratio of cells with an identity before and after the perturbation. Values are normalized per cell type. ii) Cell type transitions upon KO simulation. The first bar of the top axis represents the original cell type, and the second bar represents the cell identity resulting from the simulation. The grids represent the fraction of the original cell type (labelled in red) and their final identity. iii) Regulatory role of every gene in each cell type. TFs connecting different GRN modules are considered hubs, and they are ordered according to their connectivity to local modules: from ‘Ultra peripheral’ to ‘Kinless’ for non-hubs, and from ‘Provincial’ to ‘Kinless’ for hub TFs. iv) Disease association of every gene. TF association to disease can be risk-only, target of a disease-regulon and core of a disease-regulon.

**Supplementary Fig. 15. Related to Fig. 4.**
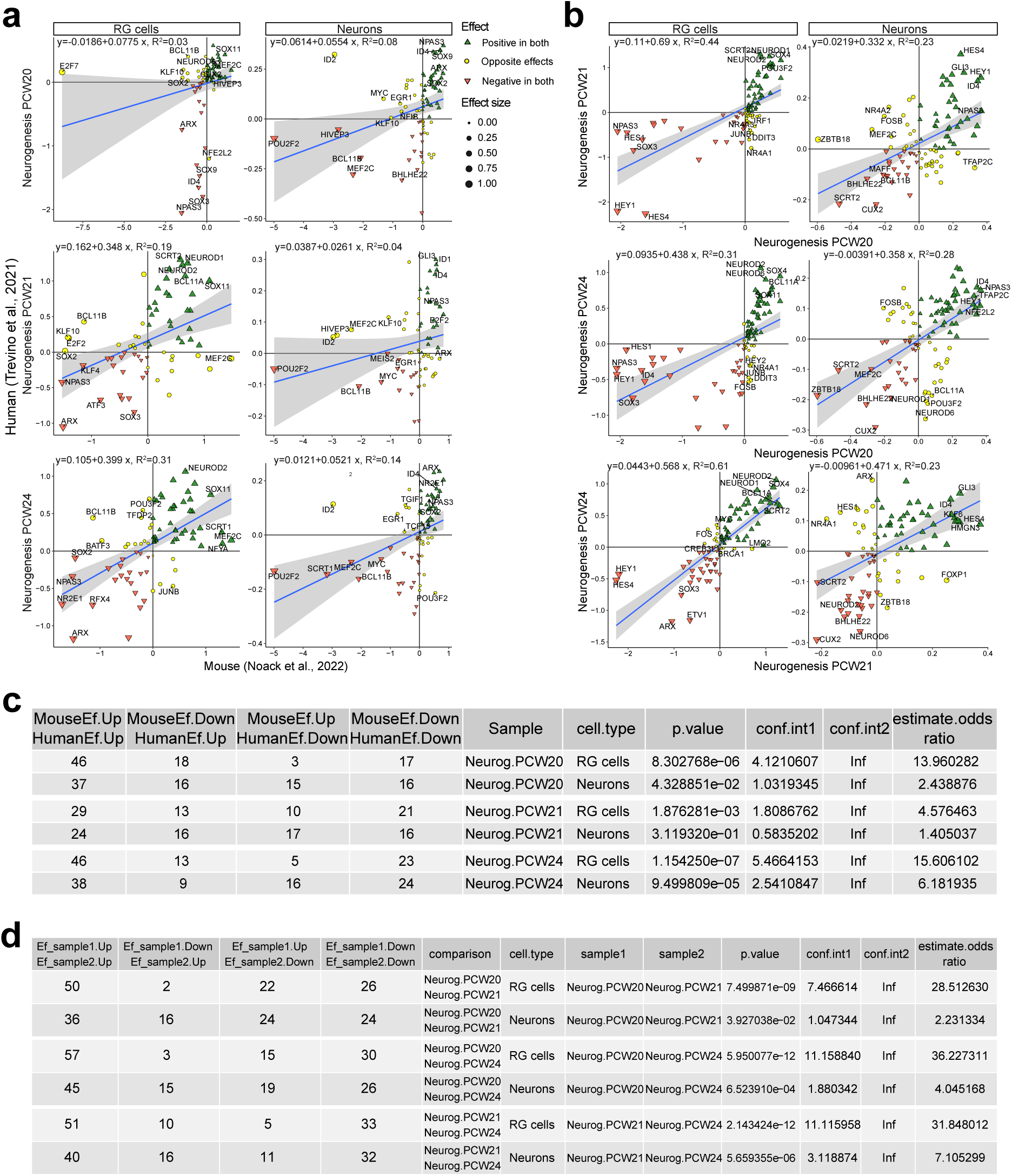
KO simulation in neurogenesis across species. (**a**) Comparison of KO simulation between mouse (x axis) and human (y axis) neurogenesis leveraging data from Noack et al, 2021 ^63^ and donors PCW20, PCW21 and PCW24 from Trevino et al, 2021 ^61^, respectively. The effect of the gene KO simulation on each cell identity group (RG cells and neurons) is shown as the ratio of final over initial number of cells, in log2 scale. Only genes knocked-out in both datasets are shown. A regression model is overlaid on the plot. The genes with a positive or negative effect in both datasets, or opposite effects after KO simulation, are indicated. **b**) Comparison of KO effects in human donors. Same as in a, one-to-one comparisons in neurogenesis of three human donors. **c and d)** Fisher tests of genes showing the same direction of effect. For each comparison, the table shows the number of genes with a positive or negative effect in both datasets, and opposite effects. The odd ratio of genes with coincident effects, its confidence interval and p-value are also indicated.

**Supplementary Fig. 16. Related to Fig. 5.**
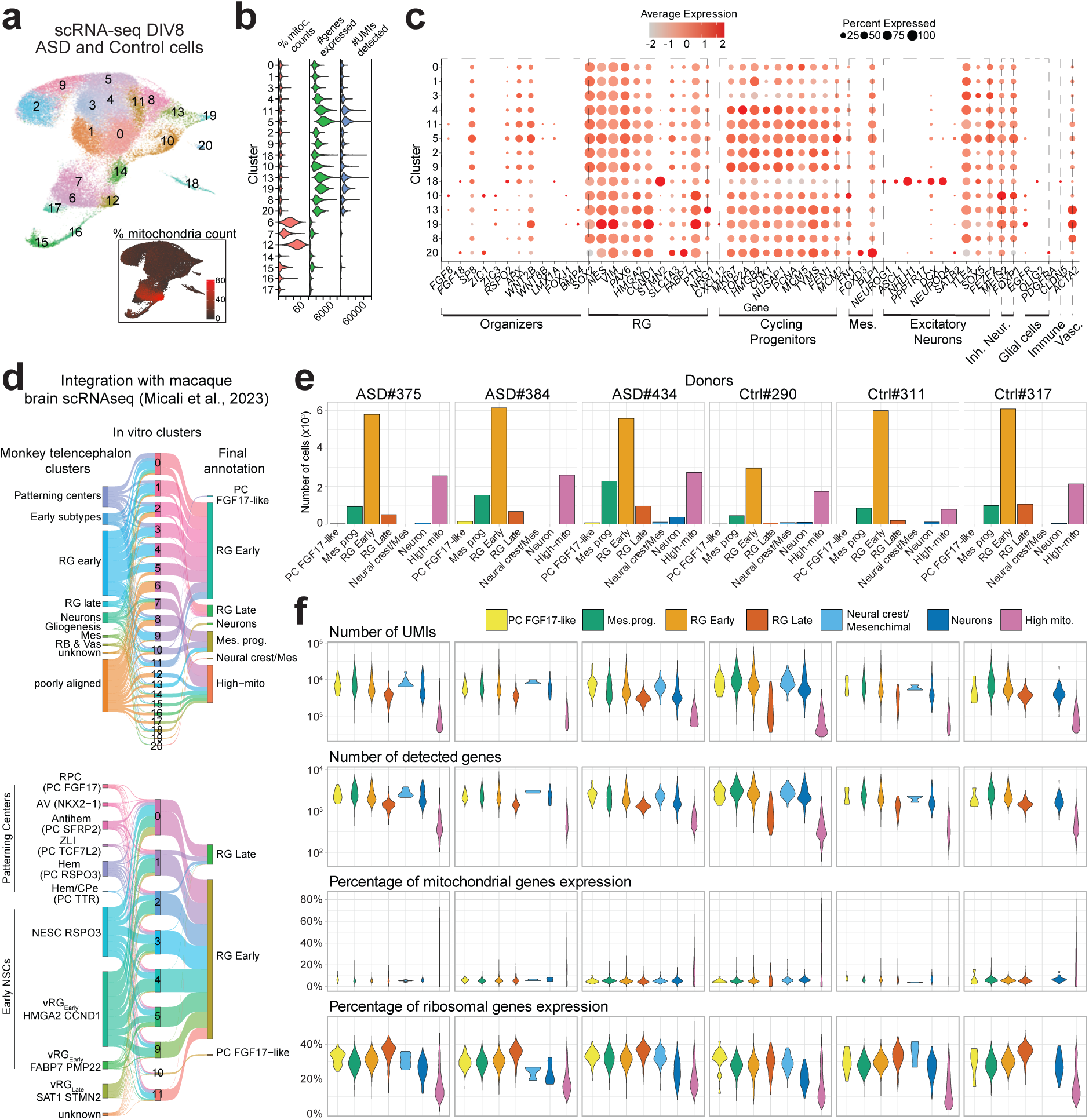
Quality overview of control- and ASD-derived in vitro NSC scRNA-seq data. **a)** UMAP showing clusters of cell types identified. **b)** Quality check of the clusters. Violin plots showing the percentage of mitochondrial counts, the number of genes detected and the number of Unique Molecular Identifiers (UMIs) per cluster. **c**) expression level (color gradient) and percentage of cells (dot size) expressing subtype markers for each cluster identified. **d**) Sankey plots showing the integration of the DIV8 scRNA-seq with macaque developing brain dataset ^37^. Left, middle and right column showing macaque annotation, raw seurat clusters and final annotation in the DIV8 dataset, respectively. The top panel includes the most representative cells of the macaque dataset and all cells in the DIV8 dataset. The bottom panel includes patterning centers and the ventricular radial glial cell (vRG) subtypes of the macaque dataset, and PC FGF17-like and RG clusters in the DIV8 dataset. **e**) Bar plot showing number of cells for each cluster in each donor. **f**) Number of UMIs, total detected genes, percentage of mitochondrial and ribosomal genes expression in each cluster for each donor.

**Supplementary Fig. 17. Related to Fig. 5.**
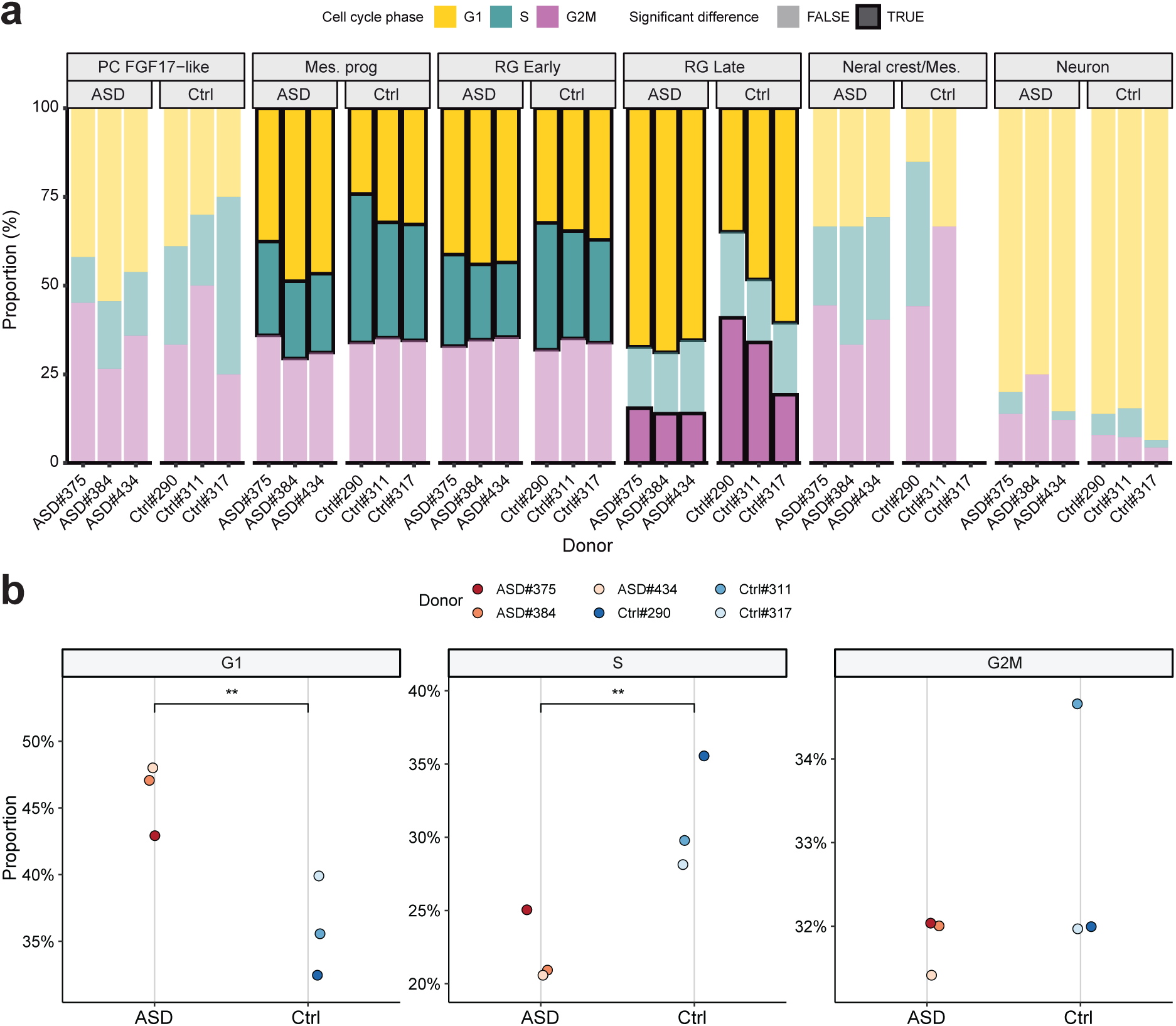
Cell cycle phase characterization of control- and ASD-derived cell lines. **a)** Fraction of cell cycle phases in every cell type in each donor. Significant differences between ASD and control cells are highlighted with black outlines. **b**) Per-donor proportion of total cells in the different cell cycle phases. Significant differences between groups are marked with “**” (adjusted p-value < 0.05).

**Supplementary Fig. 18. Related to Fig. 5.**
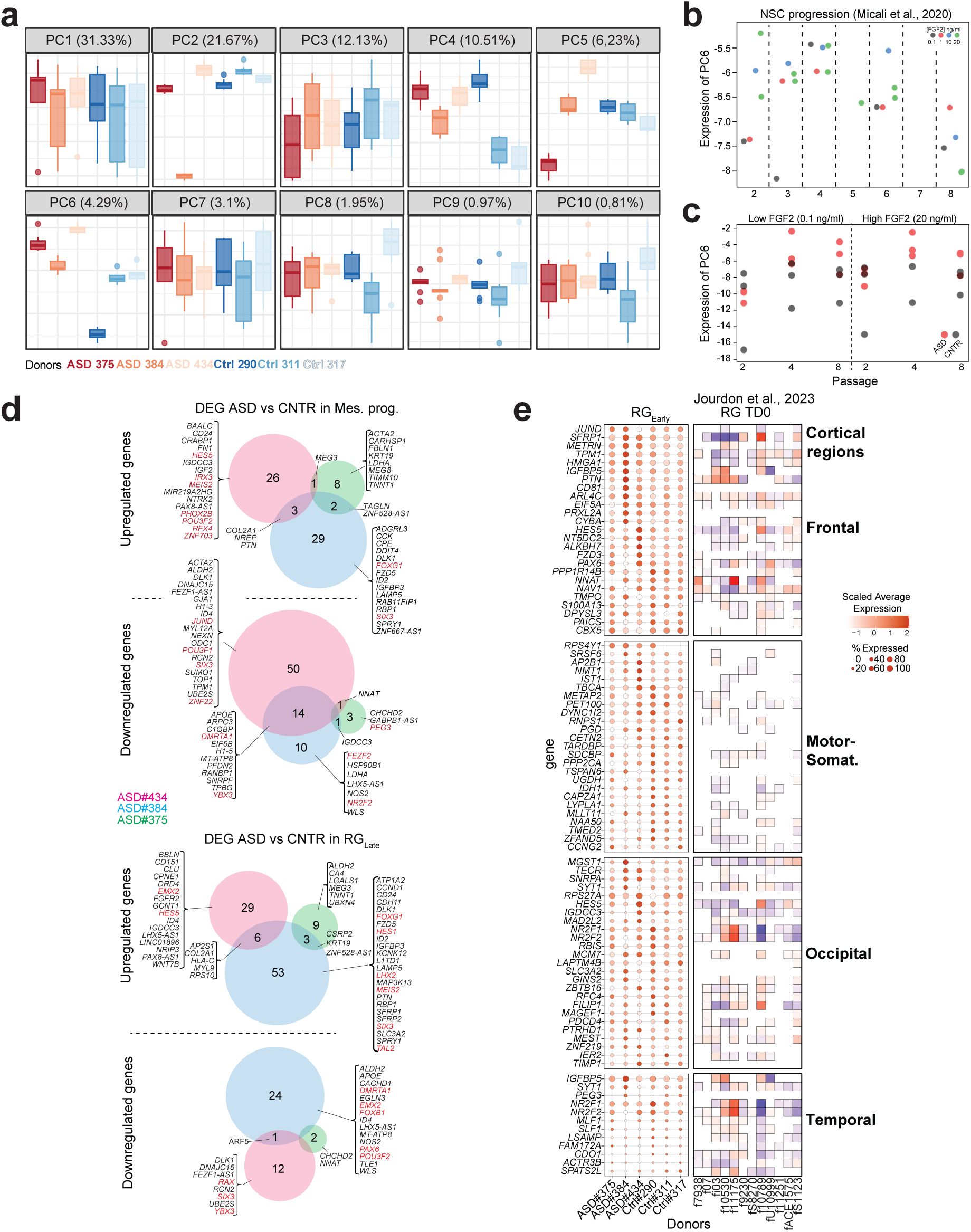
Differential gene expression analysis of control- and ASD-derived cell lines. **a)** Principal component analysis of RG_Early_ pseudobulk samples from scRNA-seq data generated from control and ASD hiPSCs at DIV 8 of neural induction. PC6 segregates control and ASD lines. **b**) Projection of Micali et al., 2020 bulk RNA-seq from sequentially passaged human NSCs into PC6. PC6 shows highest levels in PS4 NSCs (<u>NeMO/PCA</u>). **c**) Projection of bulk RNA-seq data from sequentially passaged control and ASD cell lines in vitro (see Fig. 5g and Supplementary Fig. 20) into PC6. PC6, which segregates control and ASD lines in the scRNA-seq, also shows elevated levels in these same ASD lines in the bulk passaging data (<u>NeMO/PCA</u>). **d**) Venn diagrams of DEGs in mesenchymal (Mes.) progenitor cells and RGLate in individual ASD samples versus grouped controls, evaluated in scRNA-seq data and excluding sex chromosome genes. In Mes. progenitors, 69 genes (9 TFs) up-regulated and 79 genes (9 TFs) down-regulated were found in ASD. In RGLate, 100 genes (8 TFs) up-regulated and 39 genes (8 TFs) down-regulated were found in ASD (Supplementary Table 9). **e**) Expression level (color gradient) and percentage of cells (dot size) expressing cortical region genes in RGEarly in each NSC line (left); and corresponding differential expression across ASD-control pairs in cortical organoid RG cluster at TD0, from Jourdon et al., 2023 ^34^. The genes were selected from those associated with cortical regions in progenitor cell types in Micali et al., 2023. 25 genes with the highest expression in our data that were expressed maximum in two cortical regions were considered.

**Supplementary Fig. 19. Related to Fig. 5.**
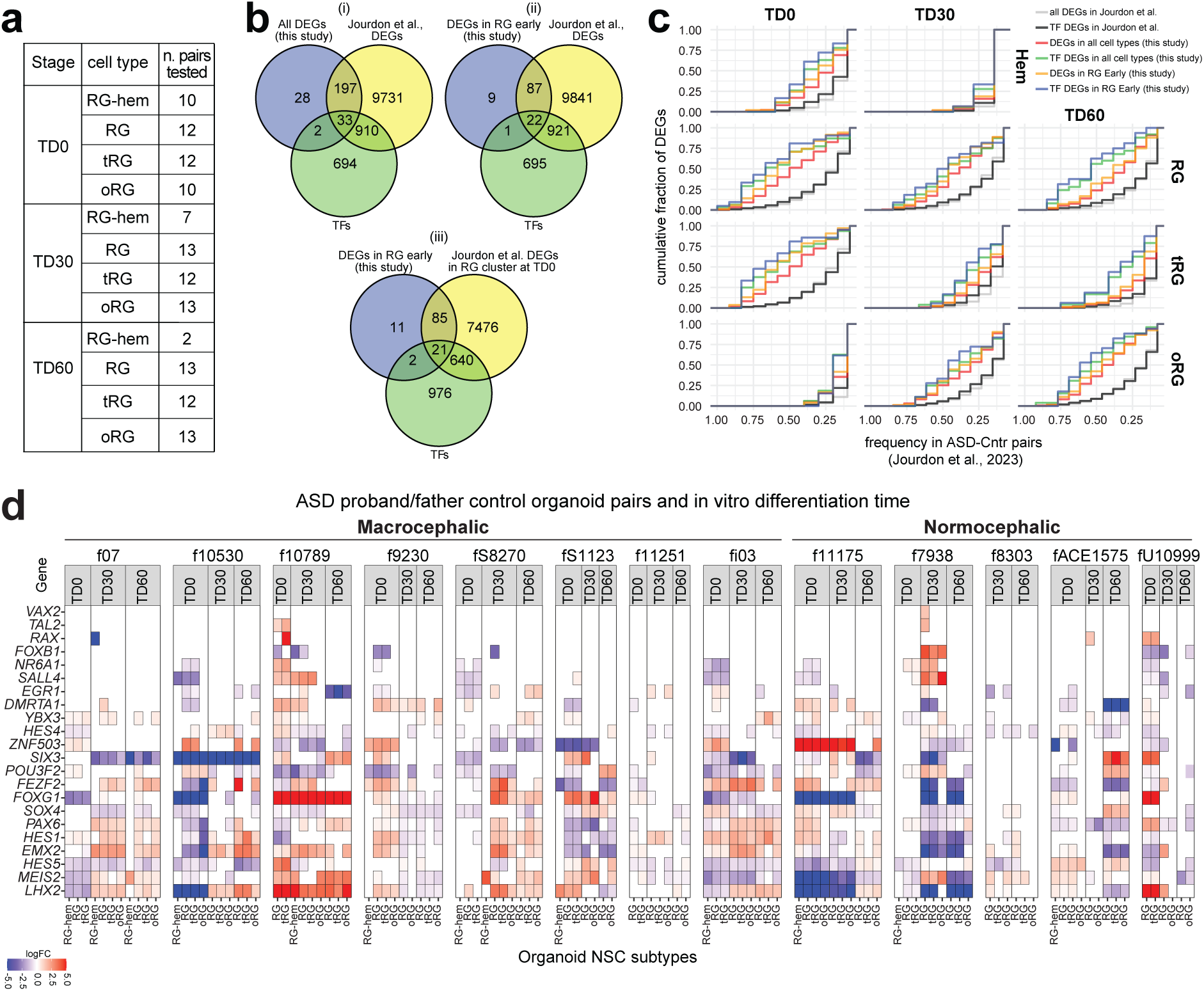
Intersection with differential gene expression in multiple ASD-control pair derived organoid lines. **a)** Number of ASD-control cortical organoid pairs, per NSC subtype (RG-hem, RG, tRG, oRG) and differentiation stage (terminal differentiation, TD0, TD30 and TD60), tested for differential expression in Jourdon et al., 2023 ^34^. In Jourdon et al., 2023, cortical organoids were generated from iPSC lines derived from ASD probands and control fathers from 13 families (or “pairs”); scRNA-seq data were collected at 3 time points (TD0, TD30 and TD60, with TD0 corresponding to the initiation of neurogenesis) and differential expression test was performed independently in each cell type (glmGamPoi, adjusted p-value < 0.01, absoluted log2FC > 0.25). **b)** Venn diagrams showing (i) intersection between all pairwise DEGs in organoid NSC subtypes (RG-hem, RG, tRG, oRG) from Jourdon et al. dataset and ASD versus Control line DEGs in any cluster of DIV8 scRNA-seq data identified in this study (Supplementary Table 9); (ii) intersection between all pairwise DEGs in organoid NSC subtypes (RG-hem, RG, tRG, oRG) from Jourdon et al. dataset and DEGs from ASD versus Control lines in RG_Early_ cluster from scRNA-seq at DIV8 identified in this study; (iii) intersection only for DEGs in RG_Early_ from this study and from organoid RG cluster of TD0 from Jourdon et al. In each diagram, TFs from both sets were identified using a list of all human TFs (in green) ^83^. **c)** Overlap of DEGs from this study (Fig. 5c) and DEGs identified in any of the ASD-control pairs from Jourdon et al., 2023 ^34^. The overlap is depicted as the cumulative fraction of DEGs identified in our study (y-axis, grouped into different DEG subsets by color) that are also found to be differentially expressed in varying frequencies among the ASD-control pairs in Jourdon et al. (x-axis). Distribution of all genes/TFs DEG in Jourdon et al. is given as reference (black/grey lines). **d)** Heatmap of pairwise DEGs between ASD proband (8 macrocephalic and 5 normocephalic)- and father control-derived organoids from Jourdon et al. for the TFs differentially expressed in RG_Early_ of the individual ASD lines identified in Fig. 5c. Note that the direction of change, although often concordant across the organoid NSC subtypes at the same stage, varied between pairs, i.e., a same gene may exhibit upregulation or downregulation across pairs. This result suggests that while perturbation of a gene is frequent across ASD lines compared to controls (as shown in S19C), the direction of change may be different among pairs.

**Supplementary Fig. 20. Related to Fig. 5.**
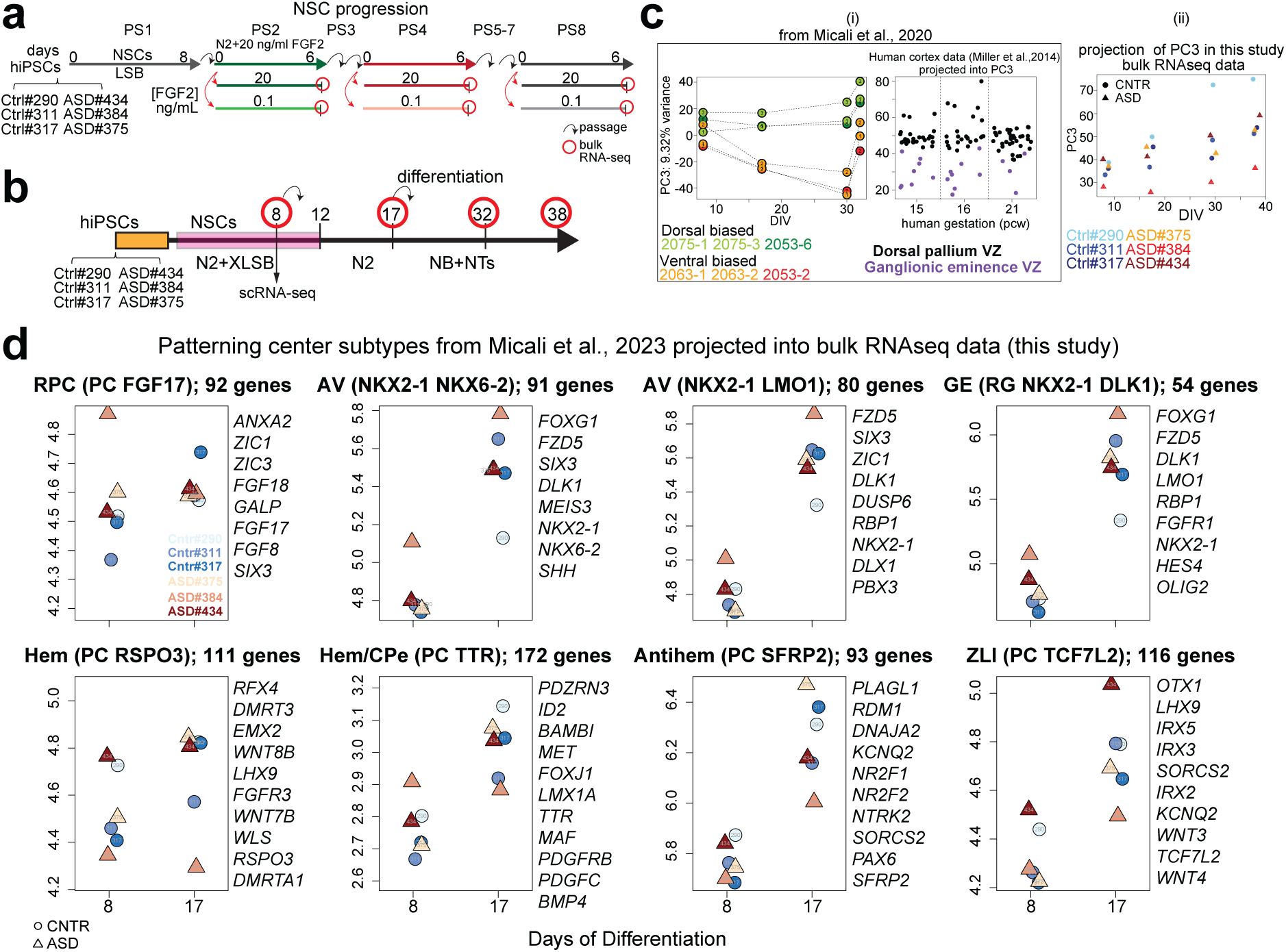
Characterization of the telencephalic identity and trajectory bias of control- and ASD-derived in vitro NSCs. **a)** NSC progression protocol for six newly-generated hiPSC lines from 3 control (#290, #311, #317)- and 3 ASD (#375, #384, #434)-affected donors showing progression of NSCs across sequential passages (PS) and FGF2 doses, similar to the scheme described in Supplementary Fig. 2. N2 + LSB was applied at PS1 for 8 days, then hNSCs were serially passaged in N2 + 20 ng/mL FGF2 every 6 days up to PS8. In parallel, cultures of NSCs at PS2, 4, and 8 were subjected to FGF2 modulation (20 or 0.1 ng/mL) for 6 days in their terminal passage, before bulk RNA collection and RNA-seq. **b)** Neuronal differentiation protocol. The same 6 iPSC lines were passaged in mTesR + Rock inhibitor, then the day after were switched to N2-B27 + XLSB medium for 12 days to induce NSCs. Cells were terminally differentiated into neurons using Neurobasal (NB) + neurothrophins (NTs) until DIV 38 as described in Micali et al., 2020 ^40^. On day 8 and 17, NSCs were passaged. Single cell suspensions for the scRNA-seq were collected at DIV8 from parallel cultures. Days for bulk RNA collection are indicated. **c)** (i) PC3 of 6 human NSC lines (2075-1 and -3; 2063-1 and -2; 2053-6 and -2) from 3 donors showing dorsal (2075-1 and -3, 2053-6, green) or ventral (2063-1 and -2; 2053-2 orange) neuronal lineage bias, based on the projection of human developing pallium and ganglionic eminence bulk transcriptomic data ^49^. These data, previously reported in Micali et al., 2020, show that hiPSC-derived NSC lines have dorsal or ventral telencephalic lineage bias even with the same neuronal differentiation protocol. Lines 2075-1 and -3 exhibit dorsal pallium bias in their differentiation trajectory; 2063-1 and -2 exhibit ventral telencephalic bias; lines 2053-6 and -2, which are 2 replicates of the same donor, show divergent lineage trajectories, i.e. dorsal (2053-6) and ventral (2053-2). (ii) To examine telencephalic differentiation bias, the bulk RNA-seq data generated from 3 control and 3 ASD-derived NSC lines, differentiated using the neuronal differentiation protocol (panel b), were projected into the Micali et al., 2020 PC3, distinguishing dorsal versus ventral NSC fates. The analysis confirmed a ventral identity for ASD#384, a dorsal identity for Cntr#290, and no clear trajectory bias for the other lines. **d)** Projection of macaque patterning centers signatures from Micali et al., 2023 ^37^ into the bulk RNA-seq of the control- and ASD-derived NSC lines at DIV8 and 17. The data highlight AV telencephalic features in #384 and slight posterior telencephalic organizer features in ASD#434 which express ZLI genes. RPC: rostral patterning center; AV: anteroventral; GE: ganglionic eminence; CPe: Choroid Plexus epitelium; ZLI: zona limitans intrathalamica.

**Supplementary Fig. 21. Related to Fig. 5.**
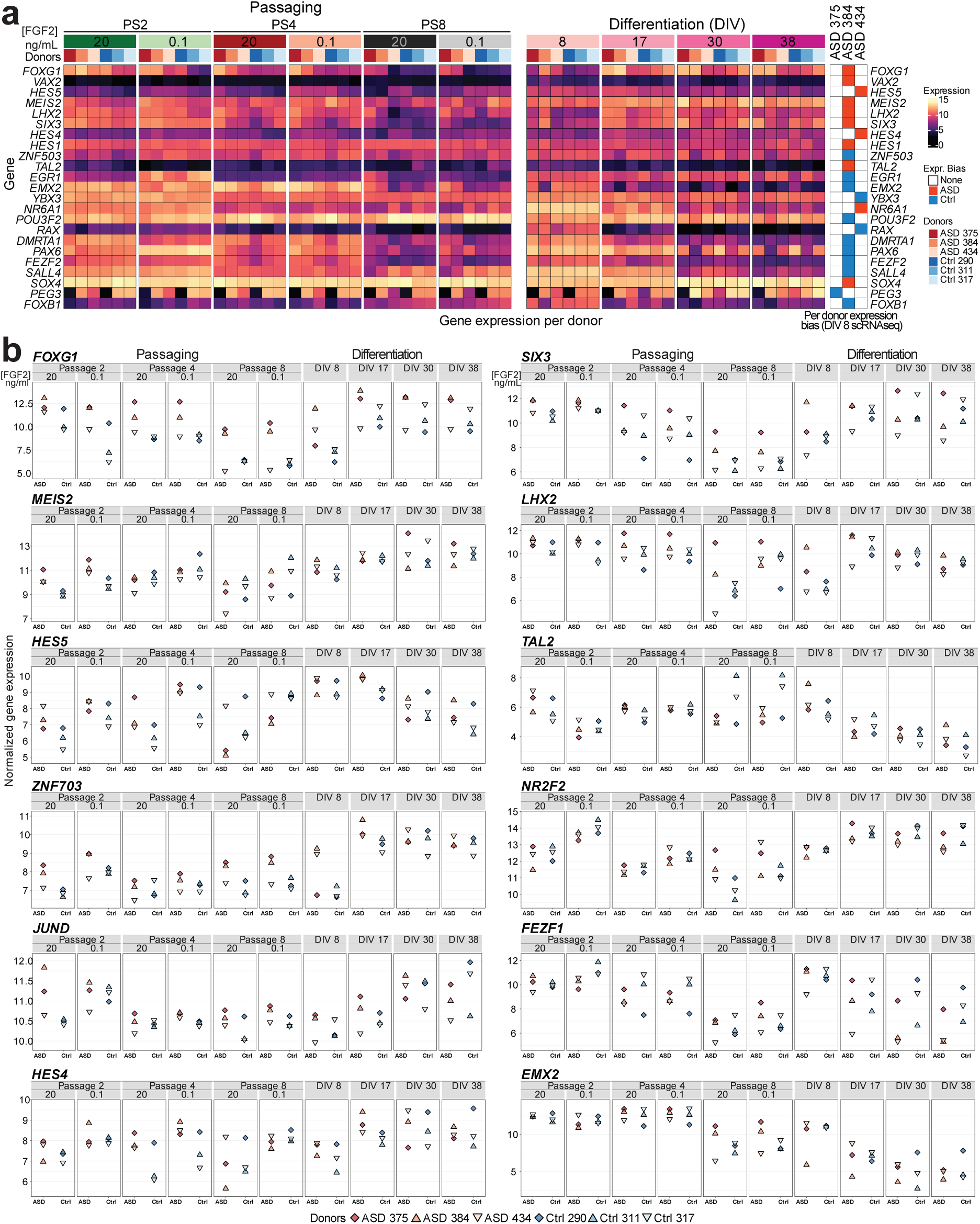
TF expression across progression and differentiation of control- and ASD-derived NSCs. **a**) Expression of TFs differentially expressed in RG_Early_ in individual ASD samples versus grouped controls from Fig. 5c, across passages and differentiation stages. The expression bias of these TFs in ASD/Control RG_Early_ from scRNA-seq analysis is on the right. **b)** Normalized gene expression (see Methods) for selected genes from panel (a), and selected DEGs (*ZNF703*, *JUND*, *NR2F2*, *FEZF1*) from Mes progenitors and RG_Late_ clusters, shown in Supplementary Fig. 18d, across passages and differentiation.

## METHODS

### Compilation of brain disease risk gene lists

We collected lists of genes associated with MCDs, including microcephaly (MIC), lissencephaly (LIS), cobblestone (COB), heterotopia (HET), polymicrogyria (POLYM), hydrocephaly (HC), focal cortical dysplasia (FCD), rare MCDs, mTORophaties (mTOR), and developmental dyslexia (Dev. Dyslexia) from literature ^20,36,38,85–94^. From GWAS datasets, we collected gene lists associated with NDDs including schizophrenia (SCZ) ^95,96^, attention deficit hyperactivity disorder (ADHD) ^97^, major depressive disorder (MDD) ^98^, bipolar disorder (BD) ^99^, neuroticism (NEUROT) ^100^, autism spectrum disorder (ASD)^76^, anorexia nervosa (AN) ^101^, the neurodegenerative Alzheimer’s (AD) ^102^ and Parkinson’s diseases (PD) ^103^, and intelligence quotient (IQ) ^104^. Genes related to developmental delay (DD) from the Deciphering Developmental Disorders Study consortium ^105^ were collected as well. Additional lists of genes associated with ASD were collected from SFARI dataset (https://gene.sfari.org), distinguished in category S (syndromic), 1 (high confidence), 2 (strong candidate) and 3 (suggestive evidence), while high-confidence ASD genes (ASD HC65) were obtained from Sanders et al., 2015 ^106^. Lists of ASD-related genes from GWAS, SFARI and ASD HC65 were combined into a single list (ASD) for gene regulatory network analyses. This compilation resulted in a collection of 2,842 genes associated with a total of 25 cortical malformations and neuropsychiatric traits provided in Supplementary Table 1.

Finally, we included genes identified through Multi-marker Analysis of GenoMic Annotation (MAGMA) for conditions such as ADHD, AD, AN, ASD, BD, IQ, MDD, NEUROT, PD, SCZ, Tourette syndrome (TS) ^107^ and obsessive-compulsive disorder (OCD) ^108^ (Supplementary Table 1). We used MAGMA to perform a gene-set enrichment analysis of GWAS signal around genes implicated in each disease among the top 5% of genes in each GWCoGAPS pattern.

Heatmaps were created using the heatmap() function in the R statistical language, as well as the levelplot() function in the “lattice” R package. Patterns from GWCoGAPS decomposition ^46^ of bulk RNA-seq data from human PSC-derived NSCs from our previous work ^40^ range from a minimum of 0 to a maximum value of 1.

### Derivation of NSCs from human PSCs

Culture and differentiation of the cells were performed in our previous work ^40^. Briefly, forebrain NSCs were generated from human embryonic stem cells (hESC) line H9 and then serially passaged in 20 ng/mL FGF2 from passage (PS)2 to PS8. Human NSCs from each passage were exposed to 0.1, 1, 10 and 20 ng/mL FGF2 for 6 days to assess their differentiation trajectory, before RNA extraction and bulk RNA-seq analysis. The 6 human iPSC lines (2063-1, 2; 2053-2, 6, 2075-1, 3) were differentiated into forebrain NSCs. hNSCs were passaged on DIV 8, then neuronal differentiation started on DIV 12. On DIV 17, hNSCs were passaged over astrocytes or without astrocytes and terminally differentiated into neurons until DIV 32. In this work, we focused on hNSCs differentiated without astrocytes and the DIV 8, 17 and 30 RNA-seq samples.

### GWCoGAPS non-negative matrix factorization decomposition of bulk RNA-seq data

The Bayesian non-negative matrix factorization (NMF) algorithm GWCoGAPS ^45,46^ decomposes the data matrix of experimental observations, “D” into 2 matrices “P” and “A”, hence D ∼ P*A. Here, “D” is the log2 RNA sequencing CPM matrix, with genes as rows and samples as columns. “P” is the pattern matrix with patterns as rows and samples as columns in which each pattern contains values for each sample, i.e., the strength of that pattern in each sample. “A” is the amplitude matrix with genes as rows and patterns as columns, with values indicating the strength of involvement of a given gene in each pattern. In this way, the “A” values for the individual patterns provide a “recipe” for reconstructing the full pattern of gene expression for each gene. GWCoGAPS was run in this work on the bulk RNA-seq data from Burke et al., 2020 ^44^ and Ziller et al., 2015 ^51^. For Burke et al., 2020 we defined 12 patterns, for Ziller et al., 2015 we defined 11 patterns by altering the number of patterns (Npats) variable. The GWCoGAPS analysis for the other datsets from Micali et al., 2020 considered in this current work are reported in the original article ^40^.

### Projection and enrichment of disease gene sets in the GWCoGAPS patterns

Gene expression data can be visualized or “projected” into a low-dimensional space defined by another data set, allowing the exploration of transcriptional modules of the first data set as they change in the new data. To relate the identity of human NSCs in vitro to cortical development in vivo, scRNA-seq data from the developing macaque ^37^ and human telencephalon ^48^ and bulk RNA-seq data from microdissected developing human ^49^ and macaque ^50^ cortex were projected into the GWCoGAPS patterns using the projectR() function in the projectR package in R ^47^. This yielded measures of the strength of each GWCoGAPS pattern in each sample, which were then averaged across common anatomical and cellular sample annotations. Using the “limma” package within Bioconductor in the R statistical language ^109^, we employed the geneSetTest() function, which uses a Wicoxon rank-sum test, to assess the enrichment of disease-associated gene sets in each GWCoGAPS pattern. The heatmaps in Fig. 1a display the negative log10 (*p*-value) of these enrichments.

### Quantification of expression changes during neuronal differentiation and maturation

We obtained expression counts from E12-15 RG cells differentiating into neurons from the single-cell mouse cortex dataset reported in Telley et al., 2019 ^56^, and from maturating cortical projection neuron subtypes reported in the DeCoN dataset ^57^. Coefficients of temporal expression changes across time were calculated using the lm() function in R, on the scaled log2 CPM (counts per million) values. Ortholog genes between mouse and human were obtained from BioMart.

### Quantification of expression changes between dorsal and ventral lineage biased hiPSC lines

We obtained expression counts from six hiPSC-derived NSC lines reported in Micali et al., 2020 ^40^ and calculated fold changes between lines forming dorsoposterior telencephalic organizer states and glutamatergic lineages (lines 2075-1, 20175-3 and 2053-6) or anteroventral states and GABAergic neuronal lineages (lines 2063-1, 2063-2 and 2053-2), using DESeq2’s Wald test ^110^.

We obtained marker genes from the clusters annotated as patterning centers (PCs), composed by RPC (PC FGF17), AV (PC NKX2-1 LMO1), AV (PC NKX2-1 NKX6-1), GE (RG NKX2-1 DLK1), GE (NKX2-1 OLIG1), hem (PC RSPO3), hem/CPe, (PC TTR), antihem (PC SFRP2), and ZLI (PC TCF7L2), from the developing macaque telencephalon dataset ^37^. We filtered out the marker genes with an adjusted p-value > 0.05 or absolute log2FC < 0.4, as well as genes with <10% expressing cells of a specific PC cluster or >50% expressing cells in the other PC clusters. We also required the fraction of expressing cells in a specific PC cluster to be at least 1.5 times higher than the other PC clusters. We represented the dorsoventral bias of the filtered marker genes at DIV8, 17 and 30, their disease association and their expression in PCs and other cell types from the macaque dataset. We showed only disease-risk genes with a significant dorsoventral bias in vitro in Fig. 2b, and marker genes with the lowest p-value (no more than 15 genes per PC cluster) in Supplementary Fig. 5b. Similarly, we represented the expression of disease-associated genes with a significant bias at DIV8 in cell clusters annotated as PCs (anteromedial pole, cortical hem and floor plate) and forebrain RG cells, and other neural cell types at different maturation phase from mouse fetal brain single-cell data ^58^ (Supplementary Fig. 5c).

### Overlap between FMRP and CHD8 targets and disease gene sets

CHD8 and FMRP targets are shown in Supplementary Table 2. CHD8 of targets were obtained from Sugathan et al., 2014 ^81^, and Cotney et al., 2015 ^79^. FMRP targets were obtained from Casingal et al., 2020 ^82^, and Darnell et al., 2011 ^80^. We tested enrichment of each set of FMRP and CHD8 targets in each disease gene set from this study and from DisGeNET database ^55^, accessed in May 2021, considering diseases with at least 40 genes, by means of Fisheŕs exact test.

### Gene regulatory network analysis of bulk RNA-seq data using RcisTarget

We leveraged RcisTarget ^111^ to build gene regulatory networks based on motif enrichment analysis within gene lists. We generated sets of genes using two criteria: 1) grouping genes by their associated diseases, which yielded disease regulons, and 2) grouping genes by their expression peak across in vitro hNSC progression from Micali et al., 2020, yielding temporal regulons. In the lists of disease genes, one gene can be associated with more than one disease. In the case of the expression-peak list, a gene can only be assign to one group. RcisTarget searches for enrichment of TF binding motifs using a precalculated database of binding motifs in gene flanking sequences (database version: v10 – human). RcisTarget was set up to identify motifs in the flanking sequences (up to 10kb around the gene transcription-start site, TSS) and we used a threshold of normalized enrichment score of 3. From the output of RcisTarget, we selected only TFs labeled as “high confidence”. Finally, we determined a network qualified as a regulon if the core TF was also included in the input list used for computing the network. Genes not expressed in the in vitro system were excluded from the analysis.

For each disease list that includes at least one regulon, we generated 1000 random lists, each containing the same number of genes as the original list, that were tested using RcisTarget. We counted the occurrences of every TF in the core TFs of the random results. We then calculated the p-value of the core TFs observed in the original lists, represented as the proportion of random lists in which the TF was identified as core TF.

Since genes in the same disease list are not necessarily co-expressed, we tested the correlation between the expression of a core TF and its target genes in the hNSC dataset from Micali et al., 2020. We used the bicorAndPvalue function, from the WGCNA R package ^112^, on the expression values of 20 samples (PS2, PS3, PS4, PS6, PS8; and 20, 10, 1, 0.1 ng/mL FGF2 for each passage). We counted the number of genes in each regulon that were significantly correlated (positively and negatively) with the core TF. The number of targets significantly correlated with a regulon was validated with another permutation strategy. We created random lists of target genes matching the number of target genes in the regulons, maintaining the proportion of genes peaking at each passage and FGF2 concentration. For each list of random target genes, we computed the expression correlations and the number of correlated targets, as described above. We used the distribution of random targets significantly correlated with the core TF to determine the p-value of our results.

The regulatory connections found in disease regulons require both TF and target gene to be associated with the same disease. To account for regulatory connections between core TFs associated with different diseases, we ran RcisTarget a second time for the diseases in which regulons were found. For each disease, we included core TFs from other regulons which were not associated with the disease, one at a time. This allowed us to uncover regulatory connections between core TFs of different diseases not identified in the initial analysis.

We computed the enrichment of disease-risk genes within temporal regulons using the fisher.test function from R. When any category in the contingency table had 0 counts, we applied Haldane correction (adding 0.5 to all categories). From this analysis, we gathered the fraction of genes associated with every disease in the regulon, the odds ratio of the enrichment and p-value.

Finally, to explore the similarity of motifs from core TFs in the temporal regulons, we retrieved databases of TFs, motifs, and motif similarity distances ^83^. Using these data, we gathered all motifs associated with our core TFs and clustered their similarities, using the default embedded hierarchical clustering function in the ComplexHeatmap R library.

### Reannotation of RG cells in Trevino et al., 2021 scRNA-seq dataset

ScRNA-seq data of the prenatal human cortex from four donors, PCW16, 20, 21, and 24 from Trevino et al., 2021 ^61^ were obtained from the Gene Expression Omnibus (GEO) dataset GSE162170. We used scanpy ^113^ to preprocess the expression data. We retained cells with less than 10% mitochondrial counts and genes with at least one count. This resulted in a dataset of 53,231 cells and 27,886 expressed genes. To classify cell-cycle phases in each cell, we scored the expression of cell-cycle phase-associated genes ^114^ and classified cells based on these genes. Next, we identified highly variable genes within each donor and selected the top 5,000 genes. We normalized the data to a value of 10000 counts per cell, log-transformed it, and scaled it. To model gene expression and integrate data across samples, we employed scVI ^115^, a deep generative model for scRNA-seq data analysis. We used the sequencing batch as a batch variable and included mitochondrial and ribosomal count fractions as covariates, as well as cell cycle scores. The latent space size was set to 10, and we obtained corresponding embeddings for all cells in the preprocessed data.

To further define early cell types present in the dataset, we selected progenitors (cycling, multipotent glial precursor cells (mGPCs), oligodendrocyte precursor cells (OPCs) and neuronal intermediate precursor cells (nIPCs)) and early, late, and truncated RG cells. We used 15 nearest neighbors in the embedding space, created an UMAP representation, and identified cell clusters using the Leiden algorithm. Using a selection of known marker genes from Micali et al., 2023 and Trevino et al., 2021, we assigned cell type identities based on clusters’ gene expression and excluded a low-count RG cluster (LQ RG), ependymal cells and interneurons. We could identify and label ventricular (v), truncated (t), and outer (o) RG cells, as well as early neurons, nIPCs, mGPCs, OPCs and astrocytes. Finally, in order to distinguish early from late states in both vRG and oRG clusters, we subclustered RG cells based on the marker genes of the mouse RG temporal progression from Telley et al., 2019, which are distinguished as early (gene modules “prog_2” and “prog_3”), mid (“prog_4”) and late (“prog_5” and “prog_6”). We chose this mouse dataset because mice possess a small proportion of oRG cells, therefore the early to late gene modules could reflect RG maturation rather than transcriptional changes from vRG to oRG cells. We found that this early-to-late gene signature was shared between vRG and oRG cells, and we used it to classify early and late vRG (vRG E and vRG L) and oRG (oRG E and oRG L) cells.

### In silico knock-out simulations in scRNA-seq data using CellOracle

We leveraged CellOracle ^62^ to simulate the impact of the perturbation of TFs on the transcriptome of each cell type along the progression of NSCs, neurogenesis and gliogenesis, using the reannotated data from Trevino et al., 2021. We considered vRG E and vRG L, tRG, oRG E and oRG L, as well as mGPCs, astrocytes and OPCs of all donors (PCW16, 20, 21, and 24) to obtain a representative dataset of the maturation of RG cells and gliogenic differentiation. To analyze the neurogenic trajectories, we pulled RG cell clusters, except tRG cells, which are late progenitors and might have reduced neurogenic potential^116^, and considered nIPCs, early neurons and glutamatergic neurons, from each donor separately. We obtained three data subsets of neurogenesis at PCW 20, 21, and 24, and one subset of RG maturation and gliogenesis comprising all ages. We conducted CellOracle perturbation analysis in each of the data subsets: RG maturation/gliogenesis, and neurogenesis PCW20, PCW21 and PCW24. PCW16 donor was not included in the neurogenic lineage since we could not obtain a clear trajectory from NSCs towards differentiated cells in the cell diffusion maps.

First, we preprocessed the data as suggested in the CellOracle pipeline. In brief, genes with 0 counts were removed, count data was normalized per cell and only the 3000 top highly variable genes were used. Then, data were normalized again, log-transformed and scaled. We embedded cells in UMAP, PHATE ^117^ and ForceAtlas2 ^118^ 2D-maps based on the scVI 10-dimensional embeddings computed previously and chose the dimensionality reduction that best fits with each trajectory. In the case of the neurogenic subsets from individual donors, PHATE maps were produced (knn: 100, phate decay: 15, t: “auto”). In the RG maturation/gliogenesis subset we first computed a diffusion map (knn: 10, number of components: 20) to then generate a ForceAtlas2 map from it (knn: 50). K nearest neighbors (knn) was selected as the pseudotime model to fit in CellOracle.

Second, sample-matched scATAC-seq data from the same study were leveraged to reconstruct a base gene regulatory network (GRN) specific to each subset of the dataset. Monocle3 ^119^ was used to preprocess the data using latent semantic indexing (LSI) as the normalization method. Peaks and peak-to-peak co-accessibility were obtained by Cicero ^84^. Peaks overlapping a TSS were annotated to the corresponding gene and only those peaks with a co-accessibility greater than 0.8 were retained. Peaks were scanned for motifs using CellOracle’s scan function, which uses the gimmemotifs ^120^ motif scanner (false positive rate: 0.02, default motif database: gimme.vertebrate.v5.0 using binding and inferred motifs, and cumulative binding score cutoff: 8) to generate an annotated peak-motif binary matrix. This subset-specific base GRN was fit to each cell type using the CellOracle functions: cell-type specific links were retrieved by fitting the GRN to the cell-type specific expression matrix using a bagging ridge regression model (bagging_number: 20, alpha: 10). The links in the resulting networks were filtered by p-value (p-value < 0.001) and from those, only the top 2000 links were retained based on their mean coefficients. After this filtering, the model was fit once more to adjust the coefficients of the preserved links.

Lastly, before CellOracle’s simulation can be performed, a simulation grid needs to be fit in the cell pseudotime map to estimate a developmental flow from the data. We manually selected the following parameters in the CellOracle pipeline: ridge alpha: 10, mass smooth factor: 0.8, grid points: 40, scale of the flow: 40. To avoid empty grid points, the knn and “min_p_mass” filter values were adjusted in each subset given the differences in the number of cells, shape and distribution of the cells in the corresponding maps (200 and 1.7E-3 in RG maturation/gliogenesis, 72 and 180 in neurogenesis PCW20, 35 and 500 in neurogenesis PCW21, and 35 and 1000 in neurogenesis PCW24). These parameters were then used to perform the simulation step as well, combined with the desired gene and expression value to simulate. We analyzed the effect of completely knocking-out the expression of a gene, i.e., simulating an expression value of 0, for each transcription factor available in the expression data and GRN. The results represent the cell-type transitions observed in CellOracle’s simulation (run for 500 steps, replicated 5 times). Gene network scores and roles were computed using built-in CellOracle functions.

### Comparison of CellOracle results

To assess the reproducibility of our CellOracle analysis in neurogenesis between donors, we compared the results obtained in the neurogenesis subsets from the separate donors. We considered three main cell states: RG cells, IPCs, and neurons. After CellOracle’s simulations, we counted the number of IPCs that transitioned into RG cells (earlier state) or into neurons (later state), in each donor. We did one-to-one comparisons between donors of the fraction of IPCs that transitioned into RG cells or neurons using a linear regression model. We also tested the number of gene knockouts with a coincident effect (“to progenitor” or “to mature neuron”) across donors using a fisher.test.

To understand if the findings in human were transferable to other species as well, we leveraged available scRNA-seq and scATAC-seq data of the mouse developing cortex from Noack et al., 2022 ^63^. Preprocessing and CellOracle analysis were performed on this dataset, following the steps described previously (embedding: UMAP from original publication, knn: 180, min_p_mass: 11). The results were compared with the PCW 20, 21, and 24 donors in human neurogenesis, using the same approach described above.

### ASD scRNA-seq data preprocessing and cell type annotation

Cell Ranger (version 6.0.1) was used to align the scRNA-seq reads to the human GRCh38 assembly (p13) and GENCODE genome annotation (release 41), followed by UMI and barcode quantification. The resulting gene-by-cell count matrices were used in scrublet to predict doublet scores ^121^. As the number of input cells per library was less than 10,000, only a few cells were predicted as doublets, which were removed for downstream analysis. To have consistent clustering across control and ASD samples, we integrated the data using Seurat CCA methods ^122^. After obtaining the integrated data matrix across control and ASD, we scaled the data and regressed out the cell cycle scores predicted by the Seurat CellCycleScoring function, followed by principal component analysis and UMAP analysis.

The cell annotation was based on canonical marker expression and integration with other existing datasets. Leveraging the number of genes and UMIs detected in each cluster as well as their percent of mitochondria reads distribution, we spotted the clusters (cluster 6, 7, 12, 14, 15, 16, 17) showing high mitochondria reads and/or low genes and UMIs. These were filtered out for the remaining analyses, resulting in 44,311 high-quality cells. Through manual inspection of curated markers, we identified cluster 18 expressing neuronal markers (e.g., *NEUROG1, NHLH1, DCX, NEUROD4*) and cluster 20 expressing neural crest/mesenchymal markers (e.g., *FOXD3, PLP1*) ^37^. We found that cluster 10 cells formed two separated populations on the UMAP, suggesting heterogeneity within this cluster. Notably, one population expressed high levels of the markers of the anteriorventral telencephalic organizers (e.g., *FGF8, ZIC1, ZIC3*). To further confirm their identity, we integrated the data with our monkey telencephalic organizer dataset ^37^ and found that the *FGF8^+^/ZIC1^+^/ZIC3^+^* population was closer to the macaque anterior neural ridge/rostral patterning center cluster (RPC, PC FGF17) on the UMAP. Therefore, we termed this *FGF8^+^/ZIC1^+^/ZIC3^+^* cell population within cluster 10 as “PC FGF17-like” cells, while the remaining cluster 10 cells were annotated as explained below. Through the integration with the macaque telencephalon scRNA-seq dataset, we confirmed the identity of the other cell clusters expressing progenitor markers such as *SOX2, NES and VIM.* Among these clusters, we found that the *SOX2*^+^/*PAX6*-low clusters (8, 13, 19, and the *FGF8^-^/ZIC1^-^/ZIC3^-^*population within cluster 10) were aligned to the monkey mesenchymal and vascular-related cell types, and accordingly were annotated as “Mes. prog.”. The majority of the cells in the clusters with high *PAX6* expression (0, 1, 2, 3, 4, 5, 9, 11), aligned to macaque early NSCs, including patterning centers progenitors, neuroepithelial stem cells (NESCs) and early vRG; the rest of the cells in these clusters were more transcriptomically similar to the macaque late vRG (Supplementary Fig. 16d). As the clusters are not one-to-one match to the macaque NSC clusters, we leveraged integrated principal components dimensions and utilized neighbor voting to predict the cell identity. Specifically, the macaque dataset was downsampled to have a balanced number of cells per cell type, followed by random sampling of 90% cells for neighbor voting prediction. The sampling process was repeated for 100 times, and the predicted label for each cell with more than 50% occurrences was retained, otherwise predicting as “unknown”. A similar label transfer procedure was used for the integration with the whole macaque scRNA-seq dataset (top panel of Supplementary Fig. 16d), except that we further calculated Local Inverse Simpson’s Index (LISI) scores and identified the poorly annotated cells (average LISI score < 1.025). The results were visualized in sankey plots using the ggsankey package. Through this analysis, we were able to predict the cells within the *PAX6*-high clusters (0, 1, 2, 3, 4, 5, 9, 11) as “RG_Early_” or “RG_Late_”. After the assignment of cell identities, we performed integration and UMAP embedding on the filtered data.

### Differential abundance test of cell cycle phases in ASD scRNA-seq samples

We used *propeller* ^123^ to test for differences in the cell cycle phase distribution between ASD and control donors. We computed the fraction of cells in G1, S and G2M for each donor. The “logit” transformation method available in *propeller* was used and the differences between ASD and control groups were tested. Note we observed similar results using the “asin” transformation. Differences with an associated false discovery rate (FDR) < 0.05 were considered significant. Additionally, we split the data into cell types and repeated the cell cycle phase proportion test between ASD and control samples in each cell type.

### Differential expression analysis between ASD and control groups in pseudo-bulk RG_Early_ from scRNA-seq data

We performed differential expression analysis between ASD and control grouped samples on pseudo-bulk samples generated from the scRNA-seq data of RG_Early_ cluster, the most abundant cell type in the data and in all donors. The approach we followed to create pseudo-bulk samples was modified from He et al., 2020 ^124^. Briefly, we first divided RG_Early_ by cell cycle phase and donor. All cells were embedded into a common neighbor graph using Seurat’s FindNeighbors function. Then, in each donor and cell cycle phase combination, we selected 3 cells at random, named *capitals*. This was the number of desired pseudo-bulk samples per group to obtain. The distances in the neighbor graph to these capitals were measured for all cells in the group and they were assigned to the closest capital. We filled the capitals until the desired number of cells per sample was reached, i.e., 314 cells. This number corresponds to the minimum number of cells per donor and cell cycle phase combination divided by the number of samples we wanted to create. Finally, for each capital, the raw counts from all cells assigned to it were aggregated into a single pseudo-bulk sample. We visualized the variation present in these data using PCA. We performed differential expression testing between the ASD and control groups using DESeq2^110^. Specifically, we employed a Wald test and we included condition (ASD or control), cell cycle phase and donor as variables in the expression model, as well as the interaction between phase and condition. Genes with an adjusted p-value < 0.05 and an absolute log2FC > 0.5 were considered differentially expressed. To represent the results, we constructed a volcano plot displaying log2FC and adjusted p-value, labeling significant genes with adjusted p-value < 10^-12.5 or in the top 30 genes by absolute log2FC. Additionally, we computed the enrichment of imprinted genes among the significant DEGs using all genes tested, by means of a one-sided Fisher’s exact test. Imprinted human genes were obtained from the National Center for Biotechnology Information database geneimprint.com, and we considered imprinted genes those labeled as “Imprinted” in the database.

### Differential expression analysis between individual ASD donors and grouped control samples in scRNA-seq data

We compared gene expression in individual ASD samples with grouped control samples. Differential expression analysis was done in each cell type present in the data using Seurat’s FindMarkers function, obtaining up- and down-regulated genes in each ASD line. To keep the same number of cells between ASD and control groups in each test, we sampled the minimum number of cells from the bigger group. Differential gene expression results were filtered according to the following criteria: sex-chromosome genes were filtered out given the sex-class imbalance in the data, adjusted p-value < 0.05, fraction of cells expressed >= 10% in the up-regulated group, with a difference of at least 5% with the opposite group, absolute log2FC >= 0.4. We required that the expression in an ASD sample tested was lower than the minimum expression found in any of the three control samples for down-regulated genes. Conversely, we required that the expression in an ASD sample tested was higher than the maximum expression found in any of the three control samples for up-regulated genes. Using the filtered results, we checked donor-specific and overlapping up- and down-regulated genes between ASD cases and grouped controls. The overlaps in up- and down-regulated genes between donors were visualized using the venneuler R package. Filtered differentially expressed genes from this individual-donor analysis were reanalyzed in our CellOracle system using neurogenesis and gliogenesis data from Trevino et al., 2021. Briefly, we reran the CellOracle pipeline by enforcing the selected genes to not be excluded during the quality control steps, as long as they were expressed. This allowed us to consider differentially expressed genes in ASD that were not included in the previous CellOracle analysis showed in Fig. 4 and related figures. The remaining computations were performed as described before.

### Differential gene expression analysis between ASD and control NSCs across passaging and differentiation

Differential gene expression between ASD and control NSCs was tested in the bulk RNA-seq data from passaging and differentiation experiments. Raw counts were provided to DESeq2 and a Wald test was performed. In the differentiation experiment, DIV, condition (ASD or control) and donor were considered as variables in the model. In the passaging experiment, passage (PS2, PS4 and PS8), FGF2 concentration (20 and 0.1 ng/mL), condition (ASD or control) and donor, as well as interaction factors between passage, FGF2 concentration and condition, were used.

For visualization, we normalized expression values using DESeq2’s variance-stabilizing transformation (vst) function. We selected TFs found to be differentially expressed in RG_Early_ cells of individual donors (see above), and we used the ComplexHeatmap library to plot the normalized gene expression of all control and ASD lines across the passaging and differentiation bulk RNA-seq datasets. We also represented the expression ratio as log2FC at every point in the experiment (passage and FGF2 concentration in the passaging experiment; DIV in the differentiation experiment). The expression of selected genes was also visualized as a scatter plot per donor.

### Projection of principal components into the bulk RNA-seq dataset

In Micali et al., 2020 ^40^, principal component (PC)3 from bulk RNA-seq data of 6 hiPSC lines (2075-1, 2075-3, 2053-6, 2063-1, 2063-2 and 2053-2) across neuronal differentiation and projection of bulk RNA-seq from dissected human brain regions ^49^ distinguished samples with a dorsal (2075-1, 2075-3, 2053-6) and a ventral telencephalic bias (2063-1, 2063-2 and 2053-2). In Supplementary Fig. 20c, we used projectR ^47^ to project our new bulk RNA-seq data from ASD and Control iPSC lines (ASD #375, #384, and #434 and Cntr #290, #311, and #317), as well as data from bulk RNA-seq from dissected human cortical regions ^49^, into this previously identified DV axis. Similarly, in Supplementary Fig. 18a-c, we show that PC6 from RG_Early_ pseudo-bulk samples derived from our DIV8 ASD and Control NSC scRNA-seq dataset segregated ASD-from control samples. Therefore, we used projectR to project bulk RNA-seq data from this study and Micali et al., 2020 into this PC.

### Projection of macaque patterning centers signatures from Micali et al., 2023 into the bulk RNA-seq dataset

To generate the signature of organizers in the macaque that would also distinguish them from NSCs, markers of organizers compared to NSC subtypes were intersected with the top 200 marker genes of organizers from Micali et al 2023 where patterning centers were compared against each other. We averaged the observed expression of the resulting marker genes in the ASD- and control-derived samples at DIV8 and 17 to measure the obtained signatures in our bulk RNA-seq dataset.

### Expression of patterning-center and cortical region signatures in NSCs in vitro and organoids

Marker genes of cortical region-specific NSCs were obtained from the developing macaque telencephalon dataset ^37^. From these data, genes associated exclusively with tRG cells were removed, since these cells were not present in our new in vitro dataset, and only those genes expressed maximum in 2 cortical regions were kept, in order to avoid unspecific regional markers. We retained the 25 genes in each cortical region with highest expression in our in vitro single-cell data. Gene markers of brain patterning centers were also obtained from Micali et al., 2023. The top 25 genes of each patterning center cluster with lowest p-value was retained. Dotplot visualizations of scaled expression per donor were created with Seurat’s DotPlot function for Mes. prog., RG_Early_ and RG_Late_ subtypes. Genes were ordered by the donor with maximum expression, and we filtered out those genes that did not reach a 3% of expressing cells in at least one donor.

### Intersection with Jourdon et al., 2023 organoid pair dataset

Pairwise ASD versus control DEGs results from Jourdon et al., 2023 scRNA-seq dataset ^34^ were subset to include only NSC subtypes defined in that study on forebrain organoids: radial glia (RG), hem-like radial glia (RG-hem), truncated/dividing (RG-tRG) and outer RG (oRG). For intersection, genes on sexual chromosomes were also excluded. Significance thresholds were used as in Jourdon et al., 2023 (adjusted-p-value below 0.01 and absolute log2FC above 0.25). The dataset consisted of 13 ASD versus control pairs, which were evaluated independently in each cell type at 3 organoid time points (TD0, TD30, TD60). The final datasets included 10,871 genes differentially expressed across any progenitor cell types or stages.

We tested whether DEGs identified in our study were more frequently differentially expressed across the ASD lines from Jourdon et al., regardless of the direction of the change (i.e., considering both upregulation and downregulation as perturbation compared to control). To derive a frequency, we divided the number of pairs with significant changes by the total number of pairs tested in each cell type and stage (which depended on the cell types captured and analyzed in Jourdon et al., 2023). We plotted the cumulative distribution of these frequencies for different sets of genes identified as differentially expressed in ASD in this study. This was then compared to the cumulative distribution of frequencies for all differentially expressed genes from Jourdon et al., The significance of the increase in frequency was tested using the Kolmogorov-Smirnov test.

### Animals

All procedures involving monkeys were performed according to guidelines described in the Guide for the Care and Use of Laboratory Animals and are approved by the Yale University Institutional Animal Care and Use Committee (IACUC). Rhesus macaque monkeys were bred in Rakic primate breeding colony at Yale. Timed pregnant females underwent caesarian section at the indicated embryonic age. E40 and E52 macaque fetal brains were dissected and immerse fixed in 4% paraformaldehyde (PFA) overnight. Fixed brains were cryo-protected in step-gradients of up to 30% sucrose/PBS for several days and then frozen. Sections were prepared at 30 µm on a Leica CM3050S cryostat.

### RNAscope

Single-molecule RNA *in situ* hybridizations were performed by Advanced Cell Diagnostics, Newark, CA, using RNAscopeTM technology. Paired double-Z oligonucleotide probes were designed against target RNA using custom software. All probes used in this study are shown in Supplementary Table 11. RNAscope LS Fluorescent Multiplex Kit (Advanced Cell Diagnostics, Newark, CA, 322800) was used with custom pretreatment conditions following the instruction manual. Fixed frozen monkey fetal brain tissue slides was manually post-fixed in 10% neutral buffered formalin (NBF) at room temperature for 90 minutes. Then the slides were dehydrated in a series of ethanols and loaded onto the Leica Bond RX automated stainer, performing the reagent changes, starting with the pretreatments (protease), followed by the probe incubation, amplification steps, fluorophores, and DAPI counterstain. RNAscope 2.5 LS Protease III was used for 15 minutes at 40°C. Pretreatment conditions were optimized for each sample and quality control for RNA integrity was completed using probes specific to the housekeeping genes *Polr2a*, *Ppib*, and *Ubc*, which are low, moderate, and high expressing genes, respectively. Negative control background staining was evaluated using a probe specific to the bacterial *dapB* gene. Coverslipping was done manually using ProLong Gold mounting media at the end of each run.

### Microscopy and imaging

Fluorescent monkey brain tissue slices were imaged using a Zeiss LSM800 confocal microscope, or a Zeiss 510 Meta confocal microscope. Panoramics of the brain slices were acquired using Zeiss Axioscan 7, equipped with cameras Axiocam 705 color and Orca Flash 4.0 V3. Z-stack and tiled images were processed using Zeiss ZEN2009 and ImageJ (v.2.0.0-rc-69/1.52p). Slight artefactual defects of DAPI intensity were manually corrected with imageJ. When necessary, fluorescence intensity or contrast was slightly adjusted using the same parameters for all the specimens using imageJ.

### Generation of hiPSC lines from control and ASD-derived fibroblasts

Human iPSCs were generated as previously described ^125^ by reprogramming human skin fibroblasts with episomal vectors pCXLE-hOCT3.4-shp53-F, pCXLE-hSK, pCXLE-hUL, and pCXLE-EGFP obtained from Addgene. Human adult fibroblasts collected from unaffected donors (n = 3, #290; #311; #317), and patients affected with idiopathic ASD (n = 3; #375; #384; #434) were nucleofected with 1.5 ug of each episome using Amaxa Nucleofector II and Amaxa NHDF nucleofector kit or Lonza NHDF Nucleofector kit (VPD-1001). Cells (2 x 10^6^) were then seeded onto 100-mm dishes coated with 1:50 diluted growth factor reduced Matrigel. Cultures were grown under hypoxic conditions in Essential 6 medium supplemented with 100 ng/mL bFGF, 100 nM hydrocortisone, and 0.5 mM sodium butyrate. Cells were re-plated 10-14 days after transduction onto 100-mm dishes coated with Matrigel at a density of 5,000 cells/cm^2^ and cultured under hypoxic conditions in Essential 6 medium supplemented with 100 ng/mL bFGF. Colonies were selected for further expansion and evaluation 24–34 days after plating. iPSC lines were cultured under hypoxic conditions in Essential 8 Flex medium on plates coated with Matrigel diluted 1:100 and were passaged using a 0.5 mM EDTA solution for long-term expansion. To validate the iPSC lines, PCR was used to confirm that the electroporated plasmids were not integrated. Karyotyping was performed by the Yale Cytogenetics Lab to ensure no chromosomal rearrangements had occurred. Teratoma assays were conducted by the Yale Mouse Research Pathology Service, and immunofluorescence of marker genes was checked to confirm pluripotency.

### Passaging and differentiation of iPSCs

Unaffected (#290; #311; #317), and idiophatic ASD (#375; #384; #434) iPSC lines were maintained as previously described ^40^, in feeder-free condition, then dissociated in single cells with Accutase (Life Technologies, A11105), plated at a density of 1 X 10^5^ cells/cm^2^ in a Matrigel (BD, 354277)-coated 6 well plates (Falcon, 35-3046) with mTeSR1 (Stem Cell Technology, 05850) containing 5 mM Y27632, ROCK inhibitor (Sigma-Aldrich, Y0503) at 37 °C, 5% CO2. ROCK inhibitor was removed after 24 hours and cells were cultured for 4 days before the next passage. The hiPSC lines were passaged or differentiated into forebrain NSCs as previously described in ^40^.

In the passaging protocol, cells were plated at a density of 6×10^4^ cells in Matrigel-coated 24 well plates (IBIDI, 82406), with mTeSR1 plus ROCK inhibitor at 37 °C, 5% CO2. Cells were switched to Aggrewell medium (Stem Cell Technology, 05893) for 2 days and then to N2 + B27. 100 nM LDN193189 (Stemgent, 04-0074) and 2 mM SB431542 (Sigma-Aldrich, S4317) were added in the medium after ROCK inhibitor withdrawal for 8 days (passage 1). Passage (PS) 1 hNSCs were passaged using Accutase. For expansion, hNSCs were plated at a density of 4×10^5^ cells (from PS1 to PS2) or 2×10^5^ cells (from PS3 to PS8) in a PLO/Fibronectin-coated 24 well plates, cultured in N2 medium with 20 ng/mL FGF2 at 37°C, 5% O2, and 5% CO2 and serially passaged every 6 days, after dissociation with HBSS. RNA was collected at the end of every passage from PS1 to PS8. FGF2 modulation was performed at PS2, PS4 and PS8, exposing cells to 0.1, or 20 ng/mL FGF2 for 6 days which then were processed for RNA extraction.

In the differentiation protocol, iPSC were seeded in mTesR medium + ROCK inhibitor which was gradually switched to N2-B27 in the first days as following: day 0 100%, day 1 75% mTesR + 25% N2-B27, day 3 50% mTesR and N2-B27, day 5 25% mTesR + 75% N2-B27, day 7 100% N2-B27. N2-B27 medium was supplemented with 2 mM XAV939 (Stemgent, 04-0046), LDN193189 (100 nM) and SB431542 (10 mM) (XLSB) for 12 days. NSCs were cultured at 37 °C, 20% O2, passaged on day 8 in N2 + B27 + XLSB; XLSB was withdrawn on day 12. On day 17, NSCs were passaged and terminally differentiated in Neurobasal medium (NB) + B27 until day 38. RNA was collected on day 8, 17, 32, 38 from neurons, using RNAeasy mini kit (Qiagen).

### NeMO Analytics

All of the newly generated RNA-seq studies as well as the existing public gene expression datasets used in this work were uploaded to the NeMO Analytics gene expression exploration environment (nemoanalytics.org) that is built upon the gEAR framework^126,127^. This entailed download of the public datasets from individual repositories and curation of sample metadata. Whenever possible, fully processed gene-tabulated data were captured, so that further analysis and exploration were based on the same version of the data used by the original authors. NeMO Analytics provides interfaces to explore the expression of individual genes across this collection of datasets, as well as visualizations of complex gene signatures, such as disease gene lists, PC analyses, or NMF decompositions. The links which bring up all the studies used in this report are in data availability. Notice that page loading in NeMO may take > 20 sec.

## Notes

### Competing Interest Statement

The authors have declared no competing interest.

